# δ-catenin controls layer-specific transcriptional maturation of astrocytes via Zbtb20

**DOI:** 10.64898/2026.05.12.724361

**Authors:** Gabrielle Séjourné, Christabel X. Tan, Justin T. Savage, Gracie Manigault, Grace Richard Jane Ding, Evelyn J. Hardin, Juan Ramirez, Ryan Baumert, Kristina Sakers, Cagla Eroglu

**Affiliations:** Department of Cell Biology, Duke University Medical Center, Durham, NC 27710, USA; Medical Scientist Training Program, Duke University Medical Center, Durham, NC 27710, USA; Life Sciences Institute, University of Michigan, Ann Arbor, MI 48109, USA; Department of Neurobiology, Duke University School of Medicine, Durham, NC 27710, USA; Department of Psychology and Neuroscience, Trinity College of Arts and Sciences, Duke U 27710, USA; Howard Hughes Medical Institute, Duke University, Durham, NC 27710, USA; Neuroscience Graduate Program, Weill Cornell Medicine, New York, NY, USA; Department of Neuroscience, Baylor College of Medicine, Houston, TX 77030, USA

**Author notes:** Senior author.

**Keywords:** Cerebral cortex, astrocytes, neurodevelopment, Zbtb20, delta-catenin, single-nuclei RNA sequencing, spatial transcriptomics, CUT&RUN, transcriptional regulation

## Abstract

Coordinated maturation of diverse neural cell types drives mammalian cortical circuit development. Disruption of this coordination is a hallmark of human neurodevelopmental disorders, yet mechanisms that synchronize transcriptional maturation across cell types remain poorly understood. Here, we identify δ-catenin (*Ctnnd2*), a component of adherens junctions, that links cell–cell interactions to transcriptional regulation. Using single-nucleus and spatial transcriptomics, we show that δ-catenin loss disrupts transcriptional maturation across neural cell types, particularly in astrocytes. δ-catenin loss impairs acquisition of layer-specific astrocyte identities and prolongs ocular dominance plasticity, indicating impaired circuit stabilization. Mechanistically, we identify the BTB/POZ transcription factor Zbtb20, which is enriched in glial cells, as a key regulator of this process. δ-catenin loss increases Zbtb20 expression, redistributes its genome-wide binding, and dysregulates its target genes. Together, these findings support a model in which δ-catenin regulates Zbtb20-dependent transcriptional programs to establish layer-specific astrocyte identities in coordination with developing cortical circuits.

**SUMMARY:** Sejourne et al report that loss of the *adherens* junction protein δ-catenin prolongs ocular dominance plasticity and disrupts astrocyte and oligodendrocyte transcriptional identity. The underlying mechanism seems to rely on the glia-enriched transcription factor Zbtb20, which is upregulated and redistributed upon δ-catenin loss, resulting in altered expression of its target genes.

## INTRODUCTION

Cortical development requires the coordinated transcriptional and functional maturation of multiple neural cell types, including excitatory (glutamatergic) neurons, inhibitory (GABAergic) interneurons, oligodendrocytes, and astrocytes. Disruption of this coordination is a hallmark of human neurodevelopmental disorders, in which multiple cellular lineages exhibit altered gene expression and function [1, 2]. While the developmental origins and diversification of these cell types have been extensively characterized, the mechanisms that synchronize their transcriptional maturation across lineages during circuit assembly remain poorly understood.

First, excitatory neurons are born from radial glial cells (RGCs) between embryonic day (E)12.5 and E16.5 in mice. They then migrate radially into the growing cortical plate and establish layers that encompass distinct axonal projection targets and unique molecular and morphological properties [3, 4]. Simultaneously (E11.5 to E16.5), inhibitory interneurons arise from a distal pool of progenitors and migrate first tangentially and then radially from the ganglionic eminences (GE) to the cortical plate, thus reaching their final cortical destinations after excitatory neurons establish a six-layered architecture [5, 6]. After the neurogenesis is completed and glutamatergic neuron layers are established, RGCs and GE progenitors switch to produce astrocytes and oligodendroglia (E18.5); to populate the cortex [7]. Each of the later-born and migrating non-glutamatergic cell types (astrocytes, oligodendroglia, and inhibitory interneurons) undergoes layer-specific transcriptional, functional, and morphological diversification throughout the first few postnatal weeks [6-11]. Thus, cortical circuit assembly involves the sequential generation and maturation of multiple neural cell types that must ultimately specialize to function within the same laminar structure, raising the question of how their developmental transcriptional programs are coordinated to generate functional cortical circuits.

An optimal cortical region to study this question is the primary visual cortex (V1), which exhibits a developmental form of experience-dependent plasticity in mammals [12-14]. Ocular dominance (OD), the preference of V1 neurons to respond to stimuli for one eye over the other, emerges during the first two postnatal weeks in mice. However, this dominance can be altered by visual manipulations during a critical period from postnatal day (P)21 to P35 [15-18]. Critical periods of OD plasticity (ODP) have also been characterized in the visual cortices of humans and other primates, though the timing is less well-defined [12, 13, 19]. In mice, the ODP critical period closes by P50, marking a transition from remodeling to stabilization of the binocular vision circuitry [20, 21].

Classical models of critical period plasticity in the visual cortex have emphasized the maturation of excitatory glutamatergic neurons and inhibitory interneurons as primary determinants of circuit refinement and stabilization [18]. In particular, the strengthening of inhibition and the maturation of parvalbumin-positive interneurons have been closely linked to the timing of critical period closure [22, 23]. However, more recent studies indicate that non-neuronal cell types also play essential roles in this process: oligodendrocyte maturation and myelination have been shown to limit neuronal plasticity, while astrocytes regulate the extracellular matrix and perineuronal net formation to promote circuit stabilization [24-27].

Together, these findings suggest that the closure of the critical period reflects the coordinated maturation of multiple neural cell types. Notably, astrocytes are uniquely positioned to integrate neuronal signals and modulate the extracellular environment, raising the question of how their transcriptional maturation is regulated in concert with other cell types to control circuit plasticity.

A growing body of evidence suggests that glutamatergic neurons play an instructive role in establishing the subtype identities of astrocytes, oligodendroglia, and inhibitory interneurons [8, 11, 28-32]. In particular, astrocytes are highly sensitive to layer-specific cues from glutamatergic neurons: disruption of neuronal layering in *Reln* conditional knockout (Reeler), *Satb2* conditional knockout, and *Dab1* conditional knockout mice results in a loss of layer-specific astrocyte identity [8, 10]. These findings support a model in which neuronal architecture provides spatially organized signals that guide the transcriptional specialization of neighboring cell types.

What are the molecular cues through which glutamatergic neurons convey layer-specific information to other cell types? A strong candidate is the *adherens* junction, which consists of transmembrane cell adhesion molecules (cadherins) and intracellular cadherin-associated proteins (catenins) [33]. Cadherins exhibit layer-specific expression in glutamatergic neurons and mediate transcellular interactions with neighboring cells [34], while catenins can transduce adhesion-dependent signals into cytoskeletal and transcriptional responses through interactions with intracellular signaling pathways and nuclear transcription factors [35-37].

One member of the p120-catenin family, δ-catenin (encoded by *Ctnnd2*), is highly enriched in the nervous system and expressed across multiple neural cell types, including astrocytes, oligodendroglia, glutamatergic neurons, and inhibitory interneurons [38]. Mutations in *CTNND2* are strongly associated with neurodevelopmental disorders (NDDs), including intellectual disability, autism spectrum disorders (ASD), schizophrenia, and attention deficit hyperactivity disorder (ADHD) [39-48]. In rodent models, δ-catenin is required for proper synaptogenesis and neuronal circuit development [49-54]. Notably, δ-catenin functions at *adherens* junctions, where it interacts with cadherins to mediate cell–cell adhesion and can also translocate to the nucleus to regulate transcription through interactions with DNA-binding factors [55-60]. This dual localization positions δ-catenin as a potential molecular link between transcellular interactions and transcriptional regulation. Given that cadherin-based adhesion mediates contacts between glutamatergic neurons and neighboring cells, δ-catenin is well positioned to couple neuron-derived cues to transcriptional programs in adjacent cell types, such as astrocytes [29, 61]. However, whether δ-catenin coordinates transcriptional maturation between neurons and glial cells, particularly during astrocyte specification to the cortical layers, remains unknown.

Here, we developed a global *Ctnnd2* knockout (*Ctnnd2*-KO) mouse model to investigate how δ-catenin regulates transcriptional maturation during cortical development. Using single-nucleus and spatial transcriptomic approaches, together with CUT&RUN profiling, we find that loss of δ-catenin disrupts transcriptional maturation across multiple neural cell types, with particularly pronounced effects in astrocytes. In the visual cortex, δ-catenin loss impairs the acquisition of layer-specific astrocyte transcriptional identities and prolongs ocular dominance plasticity, consistent with a failure of circuit stabilization.

Mechanistically, we identify the BTB/POZ transcription factor Zbtb20 as a key transcriptional regulator underlying this phenotype. Canonically, BTB/POZ transcription factors function as transcriptional repressors; however, they are highly versatile due to their protein–protein interaction domains and can also activate gene expression depending on context [62-64]. Zbtb20 is known for its roles in astrocyte differentiation and fate determination [65, 66] and has been linked to neurodevelopmental disorders in humans [67-70]. Despite this, its function in regulating transcriptional programs in astrocytes post-differentiation remains poorly understood.

We find that δ-catenin loss leads to increased Zbtb20 expression and a redistribution of its genome-wide binding in glial cells, accompanied by widespread upregulation of Zbtb20 target genes. Together, these findings support a model in which δ-catenin constrains Zbtb20-dependent transcriptional programs to enable proper astrocyte maturation within developing cortical circuits.

## RESULTS

### δ-catenin loss disrupts transcriptional development in the visual cortex

To determine the functions of δ-catenin in cortical development, we created a global *Ctnnd2* null (*Ctnnd2*-KO) mouse model. To do so, we used CRISPR/Cas9 genome editing to excise *Ctnnd2* exons 4 and 5 (**Fig. 1a**), both of which lie N-terminal to the functionally essential armadillo (ARM) domains and are requisite to all known protein-coding mRNA transcripts [71]. For all experiments, heterozygous mice were bred to produce a mix of wild-type (WT), heterozygous (Het), and knockout (KO) littermates (**Fig. 1b**). We confirmed successful *Ctnnd2* deletion by genomic PCR (**Fig. 1c**) and by immunoblotting in P14 cortical lysates using three independent δ-catenin antibodies (**Fig. 1d-e**). Our heterozygous breeding strategy produced litters of expected Mendelian ratios, size, and survival to weaning (**Fig. S1a-d**). However, *Ctnnd2*-KO mice were smaller than WT littermates throughout development (**Fig. S1e-f**) and exhibited reduced P30 brain weight proportional to the reduction in body weight (**Fig. S1g-h**). To determine if the cellular composition in the *Ctnnd2*-KO cortices was impacted at a gross level, we stained P21 visual cortex for nuclear markers of three main neural cell types: neurons (NeuN+), astrocytes (Sox9+), or oligodendrocyte-lineage cells (Olig2+) (**Fig. 1f**) [72, 73]. Loss of δ-catenin did not affect density or laminar distribution of these three nuclear markers between littermate age- and sex-matched *Ctnnd2*-WT and KO mice (**Fig. 1g**). These results show that loss of δ-catenin causes proportional reduction of body and brain weight and size; however, it does not cause gross deficits in the composition and distribution of major neural cell types in the developing visual cortex.

**Figure 1:**
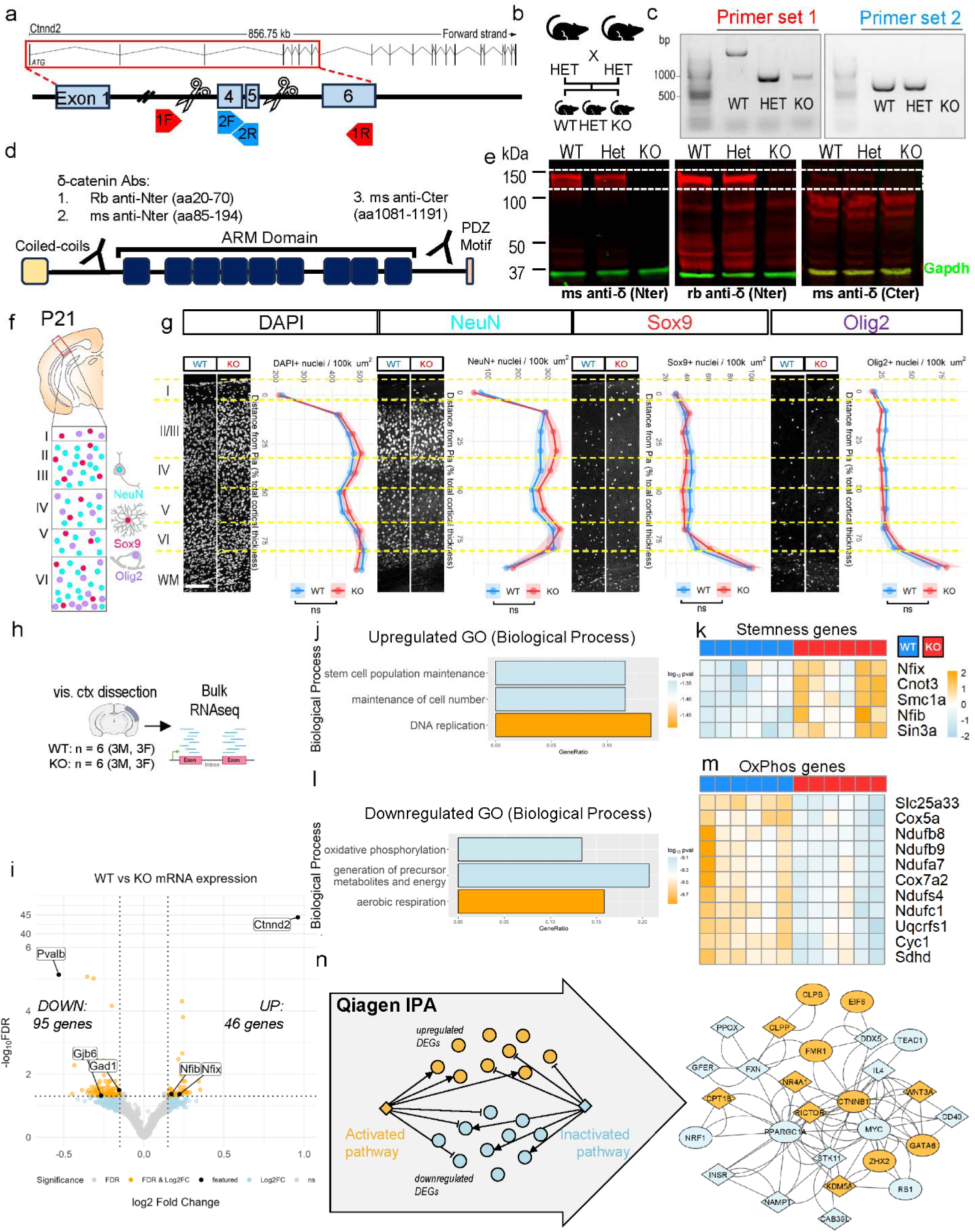
Ctnnd2-KO mice have disrupted transcriptional development in the visual cortex. A) Strategy to create *Ctnnd2*-KO mouse lacking exons 4 and 5. B) Breeding strategy: heterozygous male and female mice were bred to generate littermate wild-type, heterozygous, and knockout mice for all experiments at the ages specified. C) Genomic PCR with flanking primers demonstrates excision of exons 4 and 5 in *Ctnnd2*-KO. D) δ-catenin protein domain structure. E) Western blots from P14 cortical lysates probed with three different antibodies against δ-catenin N-terminus and C-terminus. F) P21 visual cortex was stained for principal neural cell types: nuclei (DAPI), neurons (NeuN+), astrocytes (Sox9+), and oligodendrocytes (Olig2+), and analyzed with respect to relative distance from the pia to the corpus callosum. Dashed lines indicate the boundaries between bins: bin1, 0-0.1; bin2, 0.1-0.2; bin3, 0.2-0.3; bin4, 0.3-0.4; bin5, 0.4-0.5; bin6, 0.5-0.6; bin7, 0.6-0.7; bin8, 0.7-0.8; bin9, 0.8-0.9; bin10, 0.9-1.0. Scale bars: 100_μm. G) Binned density of nuclei (DAPI), neurons (NeuN+), astrocytes (Sox9+), and oligodendrocytes (Olig2+) in the visual cortex is not different between genotypes at P21. N=8 mice of either sex per genotype, n=3 technical replicates/mouse, no significant differences between genotypes (two-way ANOVA). All values represent the mean_±_SEM. H) Visual cortex was dissected from P21 WT and *Ctnnd2*-KO mice (N=6 per genotype, 3 male and 3 female). I) Differential gene expression analysis using DESeq2 (Wald test) identified 46 up-and 95 down-regulated genes in Ctnnd2-KO visual cortex. Criteria for DEG selection: log2 fold change (FC) less than -0.15 or greater than 0.15; false discovery rate (FDR) < 0.05 as calculated by DESeq2 with Benjamini–Hochberg’s correction. J) Gene Ontology (GO) analysis of the upregulated genes identified stemness- and pluripotency-related Biological Processes significantly overrepresented. Criteria for over-represented category: FDR < 0.05, q-value < 0.2 determined by overrepresentation analysis with hypergeometric distribution. Bar length is gene ratio, fill color is FDR (Benjamini-Hochberg correction). K) Heatmap of genes related to stem cell population maintenance (GO category) plotted as read count centered and scaled in the row direction. L) GO analysis of the upregulated genes identified oxidative phosphorylation-related Biological Processes significantly overrepresented. Criteria for over-represented category: FDR < 0.05, q-value < 0.2 determined by overrepresentation analysis with hypergeometric distribution. Bar length is gene ratio, fill color is FDR (Benjamini-Hochberg correction). Bar length is gene ratio, fill color is FDR (Benjamini-Hochberg correction). M) Heatmap of genes related to oxidative phosphorylation (GO category) plotted as read count centered and scaled in the row direction. N) Qiagen Ingenuity pathway analysis was performed on all DEGs to predict upstream regulators. Cytoscape plot shows endogenous molecules that are implicated in transcriptional dysregulation effects of *Ctnnd2* deletion. Fill color is predicted activation state (ellipses are previously linked to catenin pathways).

δ-catenin directly instructs transcription via nuclear translocation and transcription factor interactions [55-60]. Therefore, to study how δ-catenin loss impacts the cortical transcriptome, we performed cell-type agnostic bulk RNA sequencing on 12 prefrontal cortex and 12 visual cortex samples dissected from 6 *Ctnnd2*-WT and 6 KO animals at P21 (**Fig. 1h, Fig. S2a-b**). Differentially expressed genes (DEGs) were defined as fold change greater than + |log2FC| > 0.15 and FDR < 0.05. This relatively permissive threshold was selected to capture distributed transcriptional changes across cell types and enable pathway-level analyses, to capture coordinated shifts in developmental gene programs. We observed more significant changes in the visual cortex samples (141 DEGs) (**Fig. 1i, Table S2**) compared to the prefrontal cortex (22 DEGs) (**Fig. S2c, Table S2**). Sequencing revealed overproduction of reads aligned to non-excised portions of *Ctnnd2* in KO tissue (**Fig. 1i**). However, as expected based on the knockout strategy (**Fig 1a**), *Ctnnd2*-KO tissue lacks reads from the excised exons 4 and 5 (**Fig. S2d**), introducing an early stop codon and abrogating protein expression (**Fig. S2e**).

Among the visual cortex DEGs, we noted upregulation of *Nfib* and *Nfix*, transcription factors with well-established roles in the differentiation and fate determination of neural lineages, including astrocytes and inhibitory interneurons [74]. Concordantly, we observed downregulation of canonical maturation markers in these cell types, including the mature astrocyte marker *Gjb6* (connexin-30) and the mature interneuron markers *Gad1* (GAD-67) and *Pvalb* (parvalbumin) (**Fig. 1i**). Gene Ontology (GO) analysis of visual cortex DEGs revealed increased expression of stemness-associated gene programs (**Fig. 1j-k**) and reduced expression of mitochondria-associated genes (**Fig. 1l-m**). Neurons, astrocytes, and oligodendrocytes all increase mitochondrial biogenesis as they transition from progenitor states to differentiate [75-78]. Consistent with GO analysis, QIAGEN Ingenuity Pathway Analysis (QIAGEN Inc., https://digitalinsights.qiagen.com/IPA) [79] (**Fig. 1n**) inferred activation of growth- and proliferation-associated pathway regulators (e.g., Wnt3a, β-catenin, and Rictor) and deactivation of the mitochondrial biogenesis regulator proliferator-activated receptor gamma coactivator 1-alpha (PGC-1α) in the *Ctnnd2*-KO visual cortex [37, 54]. These results indicate a disruption of neural cell maturation in the *Ctnnd2*-KO visual cortex.

### δ-catenin loss extends visual critical period plasticity

Given that astrocyte and interneuron maturation and specialization critically regulate the timing and extent of experience-dependent visual plasticity, we hypothesized that ODP could be disrupted in *Ctnnd2*-KO mice [20, 24-27, 80]. Depriving one eye of input for 3-5 days during the CP has been shown to induce rearrangements in thalamocortical inputs to the primary visual cortex (V1). Inputs from the non-deprived (ipsilateral) eye are strengthened, displacing the deprived (contralateral) inputs [21]. This manifests as widening of the small “binocular zone” (BZ) that receives input from both eyes. This can be quantified by visualizing the immediate-early gene *Arc* signal following stimulation of the non-deprived eye [21]. Hybridization Chain Reaction (HCR) can be used to detect *Arc* mRNA in V1 neurons, and BZ width is calculated using the zone of peak *Arc* signal in layer 4 as a proxy for BZ [21].

To study δ-catenin’s requirement for developmental ODP and visual circuitry stabilization, we performed monocular deprivation (MD) experiments in *Ctnnd2*-KO mice paired with *Ctnnd2*-WT and/or *Ctnnd2*-Het littermates (*Ctnnd2*-CTRL). The critical period (CP) during which ODP can be observed peaks between P28 and P32; however, the plasticity period is closed after P45, and BZ expansion is no longer observed if MD occurs between P49 and P53 (**Fig. 2a-c**). Consistent with previous studies in WT mice, both *Ctnnd2*-CTRL and *Ctnnd2*-KO mice subjected to 3-4 days of MD during the CP exhibited increased BZ width relative to mice subjected to 12-16 hours of MD (**Fig. 2d-e**). However, while *Ctnnd2*-CTRL mice that underwent MD after the CP (P49-P53) displayed no BZ remodeling plasticity (**Fig. 2f-g**), remarkably, *Ctnnd2*-KO mice still exhibited significant BZ expansion after MD at the same age. In summary, *Ctnnd2*-KO mice exhibit comparable plasticity to littermate controls during the typical CP window, but also remain plastic after the closure of CP. These studies demonstrate that δ-catenin is required for proper closure of critical period plasticity. Taken together with our RNA sequencing results, these findings suggest that δ-catenin controls CP closure by inducing transcriptional maturation of neural cell types, including glia and interneurons.

**Figure 2:**
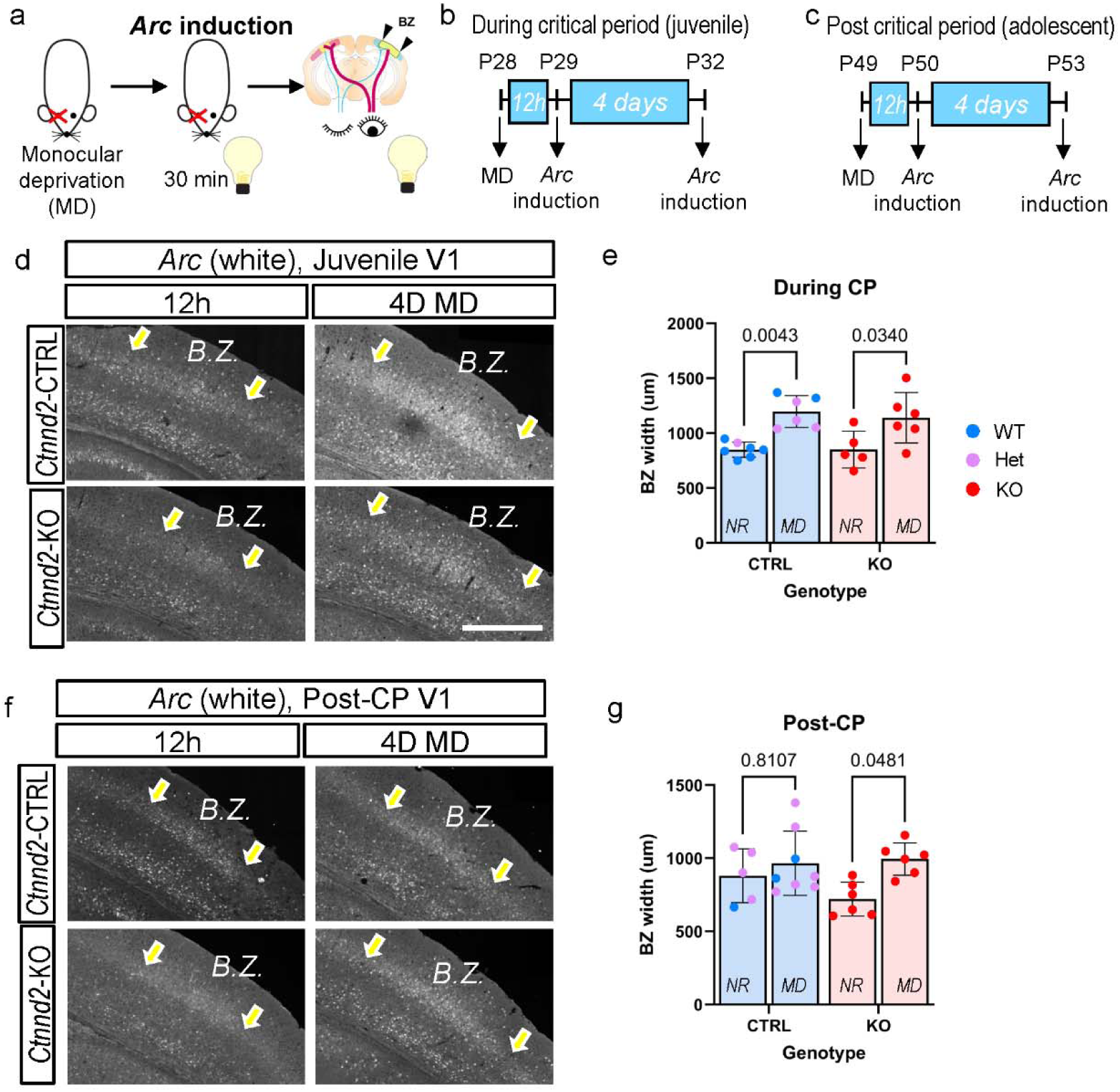
δ-catenin loss extends visual critical period plasticity. A) Experimental paradigm for measuring ocular dominance plasticity (ODP) using *Arc* induction assay. BZ, Binocular Zone. B-C) experimental timeline for measuring ODP in mice during critical period (B) and post-critical period (C). D-E) Representative images of *Arc* smFISH in the BZ of control (*Ctnnd2*-WT or HET) and KO animals 12 hours (top) and 4 days (bottom) after monocular deprivation (MD). Scale bar, 1 mm. Arrows denote *Arc* BZ width. B) Quantification of *Arc* activation width in layer 4 of hemisphere contralateral to deprived eye. N=5-7 mice per group, n=2-5 technical replicates/mouse. 2-way ANOVA with posthoc Tukey tests. NR, normal-reared; MD, monocularly deprived. F) Representative images of *Arc* smFISH in the BZ of control (*Ctnnd2*-WT or HET) and KO animals 12 hours (top) and 4 days (bottom) after monocular deprivation (MD). Scale bar, 500 μm. Arrows denote *Arc* BZ width. G) Quantification of *Arc* activation width in layer 4 of hemisphere contralateral to deprived eye. N=6-8 mice per group, n=2-5 technical replicates/mouse. 2-way ANOVA with posthoc Tukey tests.

### δ-catenin loss alters transcriptional maturation and layer-specific heterogeneity of multiple cortical cell types

To probe the cell-type-specific impact of δ-catenin loss on the transcriptome, we performed single-nucleus RNA sequencing (snRNAseq) from the visual cortices of WT (n = 3) and KO (n = 3) mice at P28. We chose this age because it corresponds to the peak of ODP, and astrocytes, oligodendrocytes, and neurons have fully differentiated into transcriptionally heterogeneous subtypes. To achieve sequencing of large numbers of both neurons and glia, we stained isolated nuclei for the neuronal nuclear marker NeuN (*Rbfox3*) and performed fluorescence-activated nuclei sorting (FANS) to separate NeuN-negative and -positive populations [9]. To enrich the NeuN-negative nuclei, the resulting populations were mixed in a 1:1 ratio (**Fig. 3a**). Nuclei capture rate was robust, with 99,966 total nuclei captured across 6 samples after pre-processing and quality control filtering (**Fig. S3a**). Major cortical cell types were identified using unbiased Leiden clustering and by comparing cluster-defining genes to the literature (**Fig. 3b**). Because *Ctnnd2* is most highly expressed in neural lineages [71], we performed comparative analyses of the clusters corresponding to glutamatergic neurons (GLU), GABAergic interneurons (GABA), astrocytes (ASTRO), and oligodendrocyte lineage cells (OL).

**Figure 3:**
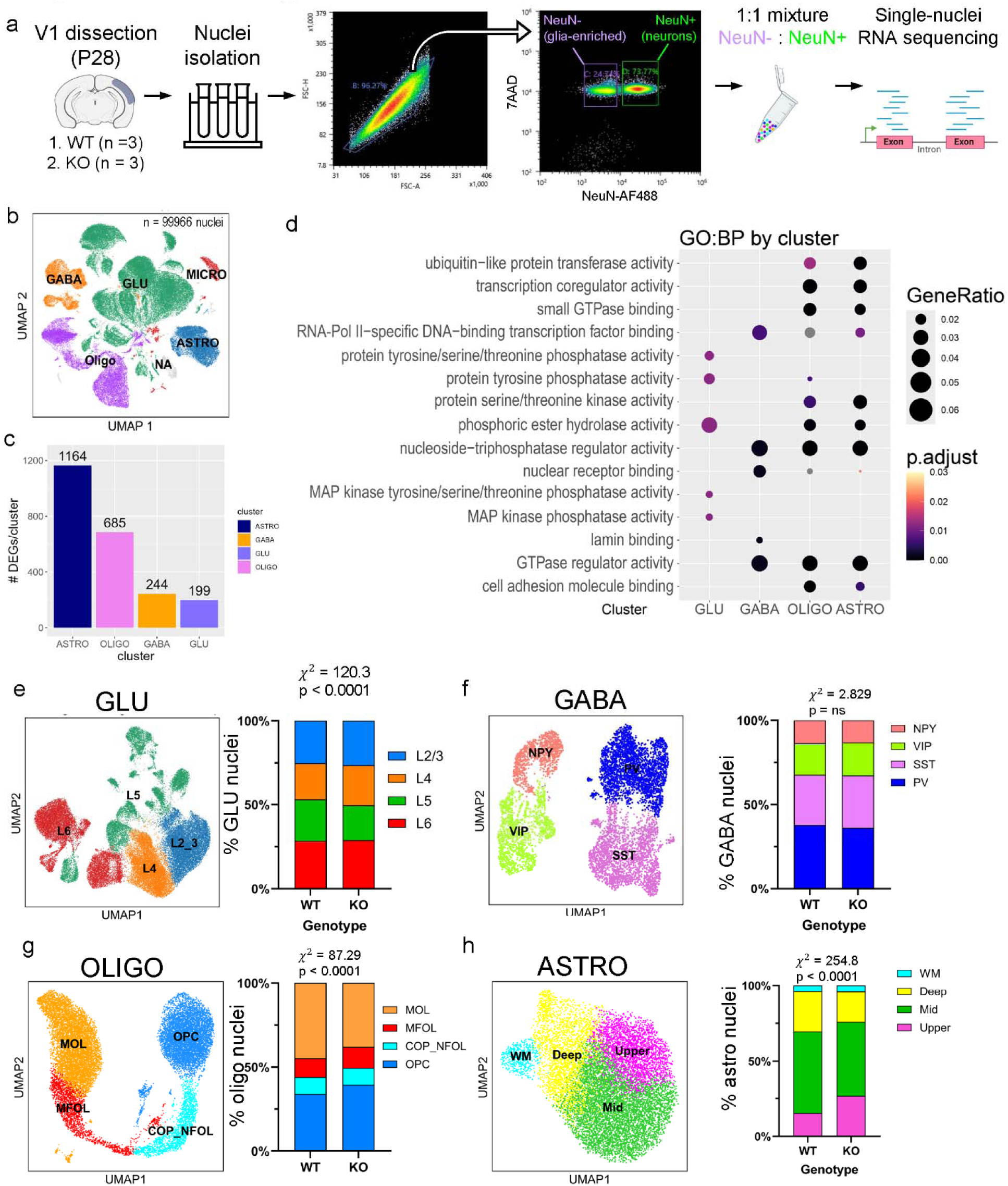
δ-catenin loss alters transcriptional maturation and layer-specific heterogeneity of multiple cortical cell types. A) Experimental paradigm: visual cortex was dissected from *Ctnnd2*-KO and WT animals at P28 (N=6 total samples, 2M and 1F per genotype). Nuclei were isolated and sorted by NeuN expression; NeuN+ and NeuN- nuclei were combined in equal proportions and sequenced using PIPseq platform for barcoding and library preparation. B) UMAP clustering of all nuclei passing quality control from WT and KO mice. Using unbiased clustering followed by manual cell-type annotation, clusters corresponding to 5 major cell types were identified: excitatory neurons (GLU), inhibitory interneurons (GABA), microglia (MICRO), oligodendrocyte lineage (OLIGO), and astrocytes (ASTRO). C) Differentially expressed genes (DEGs) within each major neural cell type, detected by Wilcoxon rank-sum test with Benjamini-Hochberg correction. Criteria for DEG selection: fold change (FC) greater than +1.25 (|log2FC| > 0.322); false discovery rate (FDR) < 0.05. D) Gene Ontology (GO) analysis of Biological Processes significantly overrepresented in the DEGs of each principal cell type. Dot size is gene ratio, fill color is FDR (Benjamini-Hochberg correction). Criteria for over-represented category: FDR < 0.05, q-value < 0.2 determined by overrepresentation analysis with hypergeometric distribution. E-H) Proportional distribution of excitatory neuron (GLU), interneuron (GABA), oligodendrocyte (OLIGO), and astrocyte (ASTRO) subtypes. E) UMAP and Barplot illustrating the proportion of Layer 2/3 (L2_3), Layer 4 (L4), Layer 5 (L5), and Layer 6 (L6) neuron nuclei found in each genotype. Chi-square test of independence, x^2^ (3) = 120.3, p < 0.0001. F) UMAP and Barplot illustrating the proportion of parvalbumin (PV), somatostatin (SST), neuropeptide Y (NPY), and vasoactive intestinal peptide (VIP) interneuron nuclei found in each genotype. Chi-square test of independence, x^2^ (3) = 2.829, p = ns. G) UMAP and Barplot illustrating the proportion of oligodendrocyte progenitor cells (OPCs), committed oligodendrocyte precursor (COP) and newly formed oligodendrocyte (NFOL), myelinating newly formed oligodendrocytes (MFOL), and mature oligodendrocytes (MOL) nuclei found in each genotype. Chi-square test of independence, x^2^ (3) = 87.29, p < 0.0001. H) UMAP and Barplot illustrating the proportion of upper, mid, deep, and white matter (WM) astrocyte subtypes found in each genotype. Chi-square test of independence, x^2^ (3) = 254.8, p < 0.0001.

First, we performed differential gene expression analysis within each lineage. To do so, nuclei were subsetted by lineage (GLU, GABA, ASTRO, OL). For each subset, a Wilcoxon rank-sum test comparing KO to WT was performed with Benjamini-Hochberg correction [81]. DEGs were defined as fold change greater than ±1.25 (|log2FC| > 0.322) and FDR < 0.05. Astrocytes exhibited the most DEGs (1164), followed by oligodendrocyte lineage (685), interneurons (244), and glutamatergic neurons (199) (**Fig. 3c, Table S3**). Comparative Gene Ontology (GO) analysis revealed a striking divergence between how glutamatergic neurons respond to δ-catenin loss compared to astrocytes, oligodendroglia, and interneurons (**Fig. 3d**). In glutamatergic neurons, kinase and phosphatase signaling were among the top perturbed biological processes, whereas in astrocytes, oligodendroglia, and interneurons, categories related to transcriptional regulation were most dysregulated. These findings show that loss of δ-catenin causes robust, cell-type-specific changes in gene expression across the neural cell lineages. Interestingly, its loss causes the most severe gene expression changes in astrocytes, which we previously found depend on cadherin/catenin signaling for their morphological development in a neuronal contact-dependent manner [29].

We next subclustered each of the four major neural cell populations to resolve their transcriptionally distinct subtypes. Single-cell variational inference (scVI) was used to correct for sample variability prior to subclustering [82]. Within glutamatergic neurons, we identified clusters corresponding to layers 1-6 (L1, L2_3, L4, L5, L6). There was a slight but significant increase in the fraction of L4 glutamatergic neurons and a reduction of other layer-specific subtypes in KO samples compared to WT (**Fig. 3e & Fig. S3b**; Chi-square test of independence: x^2^(3) = 120.3, p < 0.0001). To validate this finding, we stained for nuclear markers of layer-specific glutamatergic neuron subtypes in *Ctnnd2*-KO and WT V1 at P21 (**Fig. S4a-d**). We co-stained tissue for the L2-4 marker Cux1, L4-specific marker RORβ, and L5/6 marker Ctip2 (**Fig. S4a**). L2/3 neurons were defined as Cux1+/ RORβ-/Ctip2-, L4 neurons as RORβ+/Ctip2-, and L5/6 neurons as RORβ-/Ctip2+. We assessed the laminar distribution of these glutamatergic neuron subtypes and found no significant differences between genotypes (**Fig. S4b**). *Ctnnd2-*KOs had a minor but significant increase in the proportion of L4 (RORβ+) glutamatergic neurons compared to WT littermates (**Fig. S4c**; Chi-square test of independence: x^2^(3) = 22.28, p < 0.0001), likely due to a trending increase in their density within cortical layer 4 (**Fig. S4d**). snRNAseq resolved 4 molecularly distinct interneuron subtypes, vasoactive intestinal peptide (VIP), neuropeptide Y (NPY), somatostatin (SST), and parvalbumin (PV), which did not differ in proportion between *Ctnnd2*-KO and WT (**Fig. 3f & S3c**; Chi-square test of independence: x^2^(3) = 2.829, ns). Together, these results indicate that Ctnnd2 loss does not substantially disrupt neuronal subtype composition or laminar organization in the visual cortex.

Oligodendrocytes (OLs) are produced constitutively in the cortex by a resident population of proliferating and differentiating OL precursor cells (OPCs) [83, 84]. Thus, sequencing identified clusters corresponding to progenitor (OPC: OL precursor cells), premyelinating (COP: committed OL progenitors; NFOL: newly-formed OL), and myelinating (MFOL: myelin-forming OL, and MOL: Myelinating OL) OL lineage cells (**Fig. 3g & Fig. S3d**). The OPC fraction was significantly increased in *Ctnnd2-*KO compared to WT (Chi-square test of independence: x^2^ (3) = 87.29, p < 0.0001). To validate this finding, we performed two separate stains to differentiate stage-specific oligodendrocyte subtypes in *Ctnnd2*-KO and WT V1 at P30 (**Fig. S4e-k**). First, to differentiate OPCs and MOLs, we co-stained tissue for the pan-oligodendroglia nuclear protein Olig2, together with OPC-specific cytoplasmic protein PDGFRα and MOL-specific cytoplasmic protein ASPA (**Fig. S4e**). We assessed the laminar distribution of these oligodendroglial subtypes and found no significant differences between genotypes (**Fig. S4f**). However, among all Olig2+ cells, KOs exhibited a significant increase in the proportion of PDGFRα+ OPCs and a corresponding decrease in ASPA+ MOLs (**Fig. S4g**; Chi-square test of independence: x^2^ (2) = 63.84, p < 0.0001). These results suggest that *Ctnnd2* loss disrupts OPC progression toward the mature myelinating state.

In separate V1 tissue sections from the same animals, we co-stained for Olig2 and cytoplasmic protein Bcas1, a marker of differentiating OPCs and newly formed OLs. We also stained for Myelin Basic Protein (MBP) to differentiate premyelinating (MBP-) from myelinating (MBP+) cells (**Fig S4i**). *Ctnnd2-*KO animals exhibited no significant change compared to WT in the laminar distributions of newly-formed premyelinating oligodendroglia (NFOLs) and myelination-committed immature oligodendroglia (MFOLs) (**Fig. S4j**). There was also no change in the proportion or density of these oligodendroglia subtypes (**Fig. S4k**; Chi-square test of independence: x^2^ (1) = 0.3524, p = ns). These results suggest that *Ctnnd2* loss disrupts the progenitor-to-oligodendrocyte transition without affecting the progression of committed oligodendrocytes toward myelination.

### δ-catenin loss disrupts the layer-specific gene expression of astrocytes

Unlike glutamatergic neurons or oligodendrocyte lineage cells, astrocyte subtypes cannot be defined by single, layer- or stage-specific molecular markers. However, previous snRNAseq and spatial transcriptomic datasets have identified transcriptomic signatures corresponding to upper-, mid-, and deep-layer (border) astrocyte subtypes and to white matter (WM) astrocyte subtypes [8-10]. In our data, we detected four corresponding astrocyte sub-clusters and found that the proportion of upper-layer-like astrocytes was significantly increased in *Ctnnd2*-KO samples (**Fig. 3h** & **Fig. S3e**; Chi-square test of independence: x^2^ (3) = 254.8, p < 0.0001). To verify that our astrocyte subtype definitions aligned with the literature, we identified cluster-defining genes in WT astrocytes. To do so, WT astrocytes were subsetted, and Wilcoxon rank-sum tests with Benjamini-Hochberg correction were performed comparing each subcluster to the other three subclusters. Indeed, the resulting layer-specific transcriptional signatures from our dataset (**Table S4**) matched well to previous molecular characterizations of layer-specific astrocyte subtypes [8-10].

The significant increase in the proportion of the upper-layer-like astrocytes could be due to either an increase in their densities within upper layers of the cortex or the emergence of upper-layer-like transcriptional signatures in astrocytes in other layers. To distinguish between these possibilities, we performed Xenium spatial transcriptomic analysis of visual cortex sections from *Ctnnd2*-WT (n = 2) and Ctnnd2-KO (n = 2) littermates at P28 (**Fig. 4a**). To do so, we designed a custom 356-gene panel (**Table S5**) with strong coverage of glutamatergic, GABAergic, astrocyte, and oligodendrocyte subtypes (**Fig. S5a-b**). Following cell segmentation, we used an optimal transport-based method (TACCO) to apply cell-type annotations from our snRNAseq dataset to the Xenium dataset (**Fig. 4a**) [85]. Due to the limited number of probes, Xenium could not distinguish deep-layer astrocytes from WM astrocytes. Thus, we merged deep and WM clusters identified by label transfer into a single combined cluster (Deep_WM), resulting in a total of 3 astrocyte subclusters detected in both WT and KO samples (**Fig. 4b-c**). To identify cluster-defining genes in the Xenium dataset, WT astrocytes were subsetted, and Wilcoxon rank-sum tests with Benjamini-Hochberg correction were performed comparing each subcluster to the other two subclusters (**Table S6**). Cluster-defining genes, including astrocyte subtype-defining genes (**Fig. S5d**), closely corresponded to those in the snRNAseq dataset (**Fig. S5c**), confirming successful label transfer.

**Figure 4:**
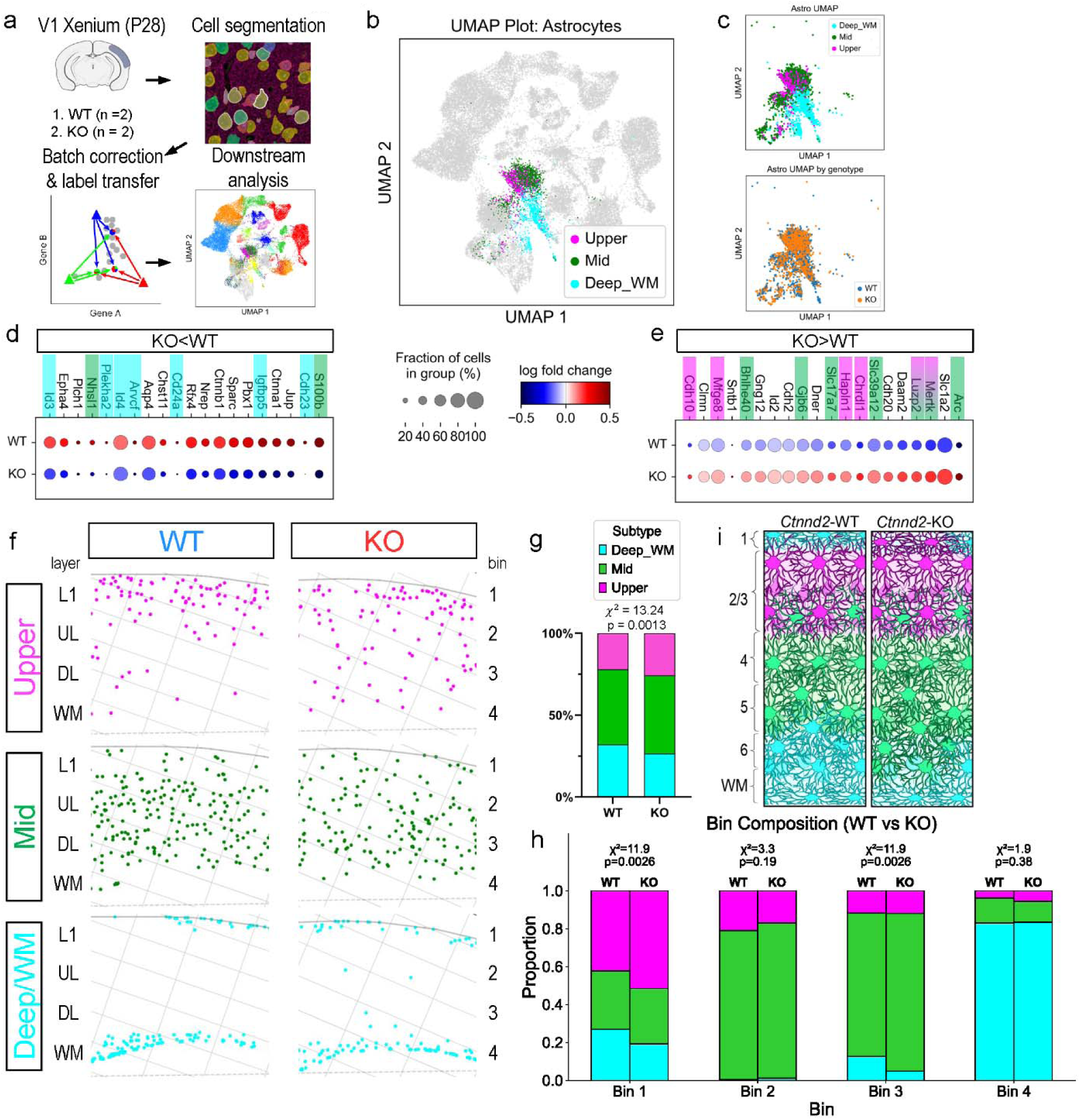
l)-catenin loss disrupts the layer-specific organization of astrocytes. A) Experimental paradigm: visual cortex was sectioned from *Ctnnd2*-KO and WT control animals at P28 (N=4 total samples, 1M and 1F per genotype). Transcripts were detected using 10X Xenium probes, followed by cell segmentation, label transfer from snRNAseq data (Fig. 3), and downstream analysis. B) UMAP of all cells detected by Xenium after selecting ROIs containing visual cortex from single hemisphere of 4 independent samples. 31469 total cells detected (15222 WT, 16247 KO). Highlighted are astrocyte subtype clusters identified by label transfer from snRNAseq. C) Higher resolution UMAP of astrocyte subtype clusters annotated by subtype (top) and genotype (bottom). 2749 total astrocytes detected (1378 WT, 1371 KO) and identified as Upper (347 WT, 396 KO), Mid (723 WT, 736 KO), and Deep_WM (506 WT, 409 KO) subtypes. D-E) Dotplots show differentially expressed genes in *Ctnnd2*-KO compared to WT astrocytes (top and bottom 20 scores by Wilcoxon rank-sum test with Benjamini Hochberg correction comparing KO to WT). For D-E, color of dot is log2 fold-change (log2FC), size of dot is fraction of cells expressing transcript. Highlight colors denote layer-specific enrichment. Criteria for layer-specific enrichment: fold change (FC) greater than 1.8 (log2FC > 0.85); false discovery rate (FDR) < 0.05. F) Spatial positioning of upper-, mid-, and deep-layer astrocyte molecular subtypes in P28 posterior neocortex sections. L1: layer 1;UL: upper layer; LL: lower layer; WM: white matter. G) Proportion of upper, mid, and deep/WM astrocyte subtypes in each genotype. Chi-square test of independence, x^2^ (3) = 13.24, p = 0.0013. H) Barplots show proportional contributions of each molecular subtype to the total astrocyte population within each of 4 equally sized cortical bins. I) Summary schematic of layer-specific changes in astrocyte subtypes.

Differential gene expression analysis on astrocytes of all layers revealed that a large proportion of the downregulated DEGs (7 out of the top 20 ranked by Wilcoxon score) were deep-layer or WM astrocyte-defining transcripts in the snRNAseq dataset (**Fig. 4d** & **Table S7**). On the other hand, a large proportion of the upregulated DEGs (11 out of the top 20) were upper- or mid-layer astrocyte-defining transcripts (**Fig. 4e** & **Table S7**). As previously shown, the “deep-layer” transcriptional subtype is present at gray matter borders, including pia and L6-to-corpus callosum transition, whereas the “upper-layer” subtype follows a gradient with the highest enrichment in upper layers (L1– L4), and the “mid-layer” subtype is enriched in L2/3-L5 [8, 86] (**Fig. 4f**). Mirroring the snRNAseq dataset, Xenium detected an increased proportion of upper-and mid-layer astrocytes in *Ctnnd2*-KO compared to WT visual cortex (**Fig. 4g**; Chi-square test of independence, x^2^ (3) = 13.24, p = 0.0013). In the absence of δ-catenin, the proportion of astrocytes with upper- and mid-like transcriptional signatures increased in upper and deep layers of the cortex, respectively (**Fig. 4h**). Correspondingly, the proportion of astrocytes with Deep_WM-like signatures decreased in these layers. These results show that *Ctnnd2* loss increases upper- and mid- layer transcriptomic signatures at the expense of deep and WM astrocyte signatures (**Fig. 4i**). This finding indicates that δ-catenin is required to specify the layer-specific transcriptional identity of cortical astrocytes.

Xenium also detected layer-specific glutamatergic neuron subtypes (L2_3, L4, L5, L6), stage-specific oligodendrocyte subtypes (OPC, COP_NFOL, MFOL, MOL), and molecularly-defined interneuron subtypes (PV, SST, VIP, NPY) (**Fig. S5e& g, Fig. S6a**). In concordance with the snRNAseq and IHC validations (**Fig. 3e-f; Fig. S4a-d**), Xenium showed a change in the proportional distribution of glutamatergic neurons with an increase in L4 subtypes (**Fig. S5e-f**), and no change in the proportions of GABAergic interneuron subtypes (**Fig. S5g-h**) in the *Ctnnd2*-KO compared to WT visual cortex. With respect to oligodendrocytes, the Xenium dataset was incongruent with our snRNAseq and IHC findings (**Fig. 3g; Fig. S4e-k**), showing no change in the overall proportion or distribution of oligodendrocyte subtypes (**Fig. S6b**). However, binned analysis of oligodendrocyte subtype proportions revealed an increased OPC fraction and reduced MOL fraction within deep cortical layers **Fig. S6c**). These results show that *Ctnnd2* loss increases OPC-like transcriptional programs at the expense of mature programs in oligodendrocytes localized to deep cortical layers (**Fig. S6d**). This finding suggests that δ-catenin is required in deep cortical layers to promote oligodendrocyte lineage progression.

In summary, using single-nuclei and spatial transcriptomics, we found that *Ctnnd2* loss impairs the acquisition of subtype-specific transcriptomic signatures. This phenotype was observed across multiple neural lineages, with the strongest effect on layer-specific astrocyte subtype specification.

### δ-catenin loss modulates Zbtb20 expression and transcription regulatory activity in glia

To identify candidate regulators of the transcriptional changes observed upon δ-catenin loss, we compared gene expression between *Ctnnd2*-KO and WT nuclei across all cell types. Among the most strongly upregulated transcripts in KO samples was *Zbtb20* (**Fig. 5a**), which was predominantly expressed in NeuN-negative (Rbfox3 gene name) nuclei at P28 (**Fig. 5b–c**). *Zbtb20* encodes a Broad-Complex, Tramtrack, and Bric-à-brac/Pox virus and Zinc finger (BTB/POZ) transcription factor with established roles in astrocyte differentiation and fate determination [65, 66]. While Zbtb20 function has been studied primarily in neural stem cells and astrocyte progenitors, its role in transcriptional regulation in fully differentiated cortical cells remains largely unexplored.

**Figure 5:**
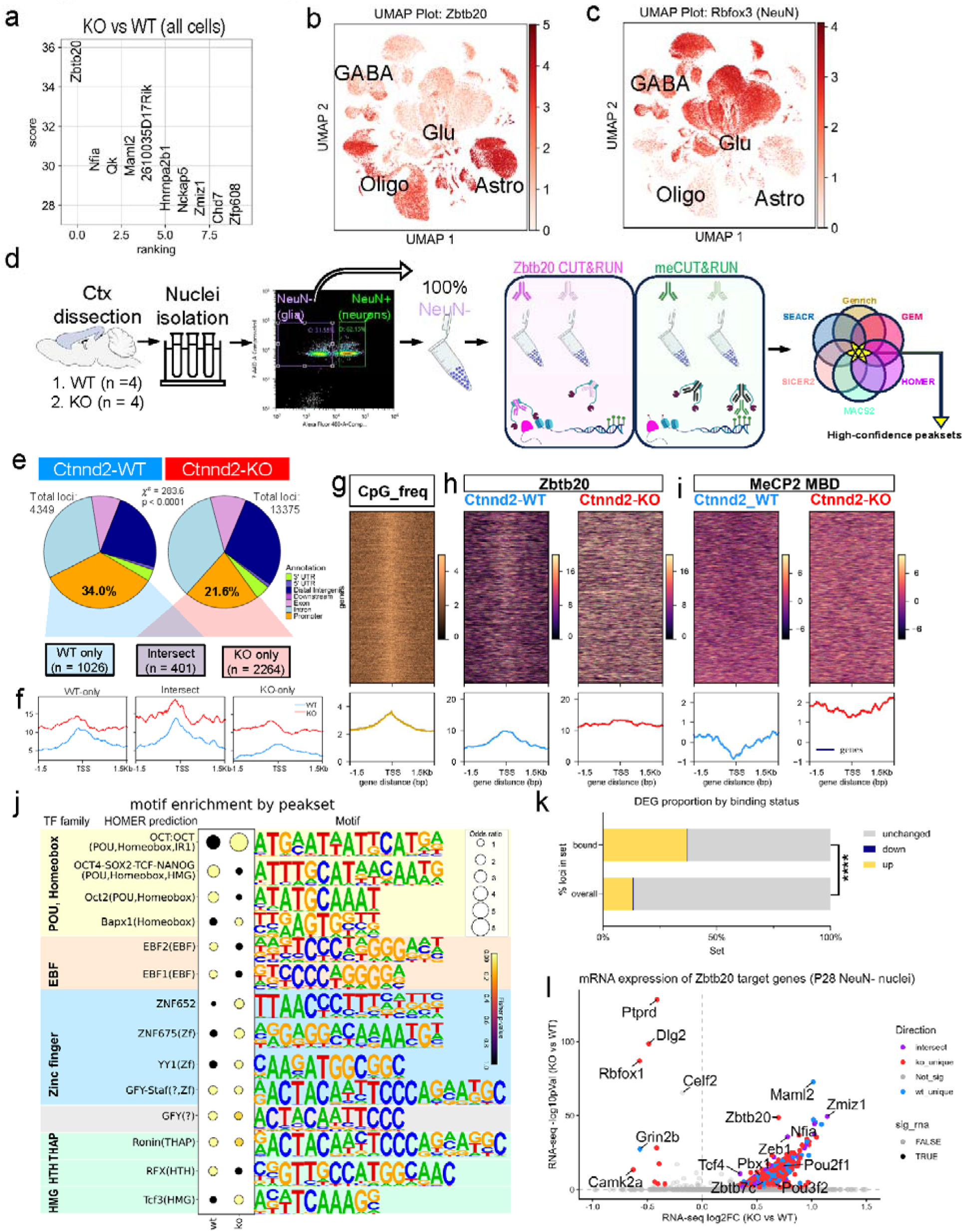
δ-catenin loss dysregulates Zbtb20 expression and transcription regulatory activity in non-neuronal cells. A) Top 10 differentially expressed genes across all nuclei from P28 snRNAseq ranked in order of z-score (Wilcoxon rank-sum test w/ Benjamini–Hochberg’s correction). B-C) UMAP plots of all captured nuclei annotated by *Zbtb20* (B) and *Rbfox3* (NeuN) (C) expression. D) Experimental paradigm for NeuN-negative *Ctnnd2*-WT vs KO comparative CUT&RUN. Conditions: Rabbit anti-Zbtb20 Antibody, Rabbit IgG Negative Control Antibody, GST-MeCP2 Methyl Binding Domain (MBD) to pull down methylated chromatin, Rabbit Anti-GST Tag Negative Control Antibody. Peak-calling was performed using 6 independent algorithms, and high-confidence peaksets were defined as those called by at least 5 out of 6. E) Feature distribution of high-confidence Zbtb20 peaksets. Chi-square test of independence: x^2^(6) = 283.6, p < 0.0001. F) Aggregate profiles of Zbtb20 coverage surrounding the TSS at Zbtb20-bound promoters. Each profile represents merged signal of N=4 biological replicates, following spike-in normalization and subtraction of corresponding negative control. G-I) Heatmaps and aggregate profiles at Zbtb20-bound promoters with CpG islands showing CpG frequency (G), Zbtb20 (H) and Mecp2 MBD (I) CUT&RUN read coverage. Each heatmap represents merged signal of N=4 biological replicates, following spike-in normalization and subtraction of associated negative control. Rows are sorted based on MecP2 MBD signal in WT condition. J) HOMER TF motif enrichment in WT-only and KO-only promoter sets. Dot size represents odds ratio of proportion of promoters in set with a given motif compared to background promoters; color represents p-value. K-L) Comparing Zbtb20-bound promoters to differentially expressed genes (DEGs) among NeuN-negative nuclei from Ctnnd2-WTvKO snRNAseq. Criteria for DEG selection: log2 fold change (FC) between -0.32 and 0.32 (|FC| > 1.25) and p < 0.05 comparing KO to WT within NeuN-negative subset. K) Proportion of DEGs among all genes vs. genes with Zbtb20-bound promoters, Fisher’s exact test: p<0.0001. L) Volcano plot of Zbtb20-bound DEGs, colored by promoter set and significance (criteria for significant DEGs as above).

To determine how δ-catenin loss impacts Zbtb20 DNA-binding activity in glial cells, we isolated NeuN-negative nuclei from WT and *Ctnnd2*-KO cortex at P28 using FANS and profiled Zbtb20 binding by performing Cleavage Under Targets & Release Using Nuclease (CUT&RUN) with a Zbtb20 antibody (**Figs. 5d & S7a**). In parallel, because DNA methylation at CpG island promoters is known to influence occupancy and regulatory function of other BTB/POZ TFs, we assessed DNA methylation by performing CUT&RUN with GST-MeCP2 Methyl Binding Domain (MBD) to pull down methylated chromatin [87, 88]. Rabbit IgG and Rabbit anti-GST Tag antibodies were used as negative controls to assess nonspecific background signal in Zbtb20 and MBD CUT&RUN, respectively. Normalization and quality control analyses were performed for both Zbtb20 and MBD CUT&RUN, as well as their respective negative controls. The CUT-and-RUN Analysis Suite (CaRAS) was used to perform peak calling to identify putative Zbtb20 binding sites. The CaRAS pipeline was run separately for each genotype (*Ctnnd2*-WT and KO), with each run comprising 4 target and 4 negative-control replicates. Zbtb20 peaks detected by at least 5 of 6 peak-calling algorithms (**Fig. S7b-c**) were selected to form consensus peak sets, which were used for downstream analyses.

A greater overall number of Zbtb20 peaks were detected in *Ctnnd2*-KO compared to WT samples (13375 vs 4349); however, a smaller proportion of KO peaks were located in promoter regions within 1.5kb of a Transcription Start Site (TSS) compared to WT (21.6% compared to 34.0%, Chi-square test of independence: x^2^(6) = 283.6, p < 0.0001) (**Fig. 5e**). We identified 1026 Zbtb20-bound promoters unique to WT, 2264 unique to KO, and 401 shared. Notably, Zbtb20 binds its own promoter in both *Ctnnd2*-WT and *Ctnnd2-*KO conditions (**Fig. S7d**), suggesting it regulates its own expression. Read counts from Zbtb20 CUT&RUN were higher in *Ctnnd2*-KO samples across all Zbtb20-bound promoters (**Fig. 5f**), in line with increased Zbtb20 expression in the KO. Promoters with Zbtb20 binding peaks called only in WT still showed Zbtb20 binding in KO, but it was dispersed. On the other hand, promoters with peaks called only in KO exhibited low binding signals in WT samples. For the following downstream analyses, we combined all Zbtb20-bound promoters into a single union set. After filtering for genes expressed in the P28 visual cortex snRNAseq, the final union promoter set consisted of 3568 loci.

We next set out to determine if Zbtb20-bound promoters contained CpG islands and whether their methylation differed between *Ctnnd2*-WT and KO samples. To do so, the union promoter set was cross-referenced with the UCSC table browser CpG island track. CpG islands are defined as having a GC content of at least 50%, a length greater than 200 base-pairs, and a ratio greater than 0.6 of observed to expected CG dinucleotide count [89]. Among 3568 Zbtb20-bound promoters, 2118 had a CpG island overlapping with the TSS (**Fig. 5g**). We analyzed MeCP2 MBD and Zbtb20 binding at these CpG island promoters (**Fig. 5h-i**). In WT, methylation at CpG islands displayed a local minimum at the TSS, and Zbtb20 binding inversely correlated with DNA methylation (**Fig. 5h**). In contrast, this correlation was lost in KO samples, with uniform methylation and Zbtb20 binding throughout the TSS and surrounding region (**Fig. 5i**). These results show that Zbtb20 binds to unmethylated CpG islands at TSSs in wild-type glia and loss of *Ctnnd2* uncouples Zbtb20 binding from DNA methylation at CpG island promoters.

In addition to CpG content and methylation pattern, TF binding also depends on DNA sequence motifs surrounding the TSS. It is possible that δ-catenin loss changes which DNA motifs Zbtb20 binds. To probe this possibility, we screened the WT- and KO-specific Zbtb20-bound promoters for enrichment of known TF-binding motifs (**Fig. 5j** & **Table S8**). Both WT and KO promoter sets showed enrichment of several zinc-finger binding motifs, consistent with previous reports that Zbtb20’s first three zinc finger domains (ZF1-3) are critical for recognizing and binding to gene promoters [90]. Both WT and KO promoters were also enriched for octamer (OCT) motifs, which are targeted by POU-III TFs (e.g., Brn-1, Brn-2) that critically regulate vertebrate neurogenesis [91]. Interestingly, the Wnt-responsive element (WRE), which is highly associated with β-catenin signaling via TCF/LEF family transcription factors was enriched over background in the KO promoter set but not in wild-type. Taken together, these findings demonstrate that δ-catenin deletion reduces Zbtb20-binding specificity, resulting in a disrupted binding pattern at CpG island promoters and suggest enhanced occupancy of Zbtb20 at sites normally bound by repressive TFs.

These findings also suggest that the gene expression changes we have observed in the *Ctnnd2* KOs may be at least in part due to dysregulated Zbtb20 function. To test this possibility, we subsetted non-neuronal (*Rbfox3*/NeuN-negative) nuclei from the snRNAseq dataset (**Fig. S7e**) and performed differential expression analysis, detecting 4343 DEGs with a fold change greater than ±1.25 (|log2FC| > 0.322) and FDR < 0.05 (Wilcoxon rank-sum test with Benjamini-Hochberg correction). Of genes with Zbtb20 binding peaks within their upstream promoters, 37.14% (1325 of 3568) are differentially expressed, compared to 13.48% of all genes detected by snRNAseq (p < 0.0001, Fisher’s exact test) (**Fig. 5k**). Almost all (1315 of 1325) of the Zbtb20-bound DEGs are upregulated in *Ctnnd2*-KO samples, regardless of whether Zbtb20 binding is lost (WT-only promoters), gained (KO-only promoters), or maintained (shared promoters) in *Ctnnd2*-KO (**Fig. 5l**), suggesting that δ-catenin is required for Zbtb20-mediated gene repression or suppresses Zbtb20-mediated gene activation. Notably, many of the DEGs whose TSSs are bound by Zbtb20 are TFs with glia-enriched expression patterns (**Fig. 5l**, **Fig. S7f-g**), suggesting that Zbtb20 and δ-catenin together regulate a TF network underlying glial development.

Taking the above evidence together, we propose the following working model (**Fig. S7h**): Zbtb20 normally binds demethylated CpGs within CpG island promoters, repressing gene transcription in a δ-catenin-dependent manner. In the absence of δ-catenin, methylation and Zbtb20 binding patterns are disrupted at CpG island promoters, and the associated genes are derepressed. δ-catenin loss also induces an increase in Zbtb20 occupancy at sites normally bound by repressive TFs, such as the TCF/LEF family. At these ectopic binding sites, Zbtb20 functions as a transcriptional activator, turning on expression of the associated genes.

### δ-catenin instructs layer-specific transcriptional identity of cortical astrocytes via Zbtb20

Because δ-catenin loss profoundly disrupted layer-specific astrocyte gene expression, we hypothesized that Zbtb20 may regulate these transcripts in astrocytes. To probe this possibility, we performed astrocyte-specific Zbtb20 CUT&RUN in cortices of P21 WT mice (**Fig. 6a**). In the same nuclei preparations, we also performed trimethylated histone H3 lysine 4 (H3K4me3) CUT&RUN to identify actively transcribed gene promoters. Rabbit IgG was used as a negative control to assess nonspecific background signal in both the Zbtb20 and H3K4me3 CUT&RUN assays. Astrocyte nuclei were labeled with GFP fused to the nuclear membrane protein SUN1 using the CAG-Sun1/sfGFP transgenic mouse line and injecting the astrocyte-specific Cre AAV (GfaABC1d-Cre-4x6T) at P3 [92]. GFP+ astrocyte nuclei were isolated from P21 cortices using FANS (**Figs. S8a**). We validated the astrocyte-specificity of this approach by isolating RNA from GFP+ nuclei and performing quantitative Real-Time PCR (qPCR) for cell-specific transcripts, which showed a greater than 4-fold increase in astrocyte-specific *Aqp4* and greater than 4-fold reduction in neuron-specific *Snap25* (**Fig. S8b**).

**Figure 6:**
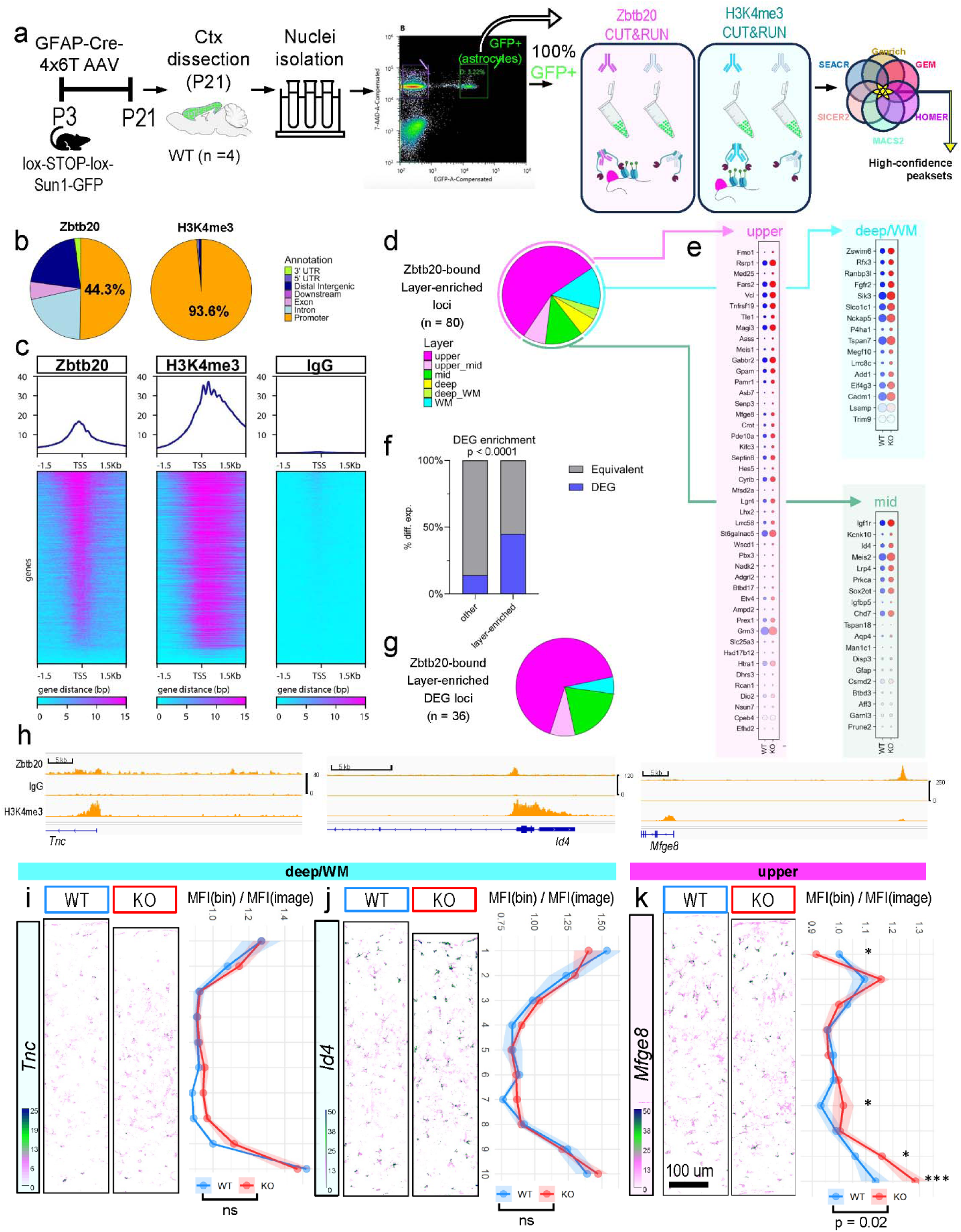
δ-catenin cooperates with Zbtb20 to control layer-specific gene expression in astrocytes. A) Experimental paradigm for astrocyte-specific CUT&RUN. Peak-calling was performed using 6 independent algorithms, and high-confidence peaksets were defined as those called by at least 5 out of 6. B) Feature distribution of Zbtb20 (left) and H3K4me3 (right) consensus peaksets. C) Heatmap of Zbtb20, H3K4me3, or IgG control CUT&RUN read coverage surrounding TSS of Zbtb20-bound promoters (N=4 biological replicates each). D-G) Analysis of layer enrichment and δ-catenin-dependent expression of Zbtb20 target genes. D) Pie charts showing layer distribution for all layer-enriched Zbtb20 target genes. Layer enrichment defined as greater than 1.8-fold enriched in at least one layer. E) Dotplots of KO vs WT expression data from snRNAseq for layer-enriched genes bound by Zbtb20. Color of dot is log2 fold-change (log2FC), size of dot is fraction of cells expressing transcript. Wilcoxon rank-sum test with Benjamini-Hochberg correction. F) Barplot showing proportion of all Zbtb20 targets that are differentially expressed in *Ctnnd2*-KO compared to WT astrocytes (snRNAseq). Criteria for DEG selection identical to previous figures. G) Pie charts showing layer distribution for all layer-enriched Zbtb20 target genes that are differentially expressed in *Ctnnd2*-KO compared to WT astrocytes (snRNAseq). H) Zbtb20, IgG, and H3K4me3 CUT&RUN read coverage at loci of transcripts enriched in layer-specific astrocyte subtypes: *Tnc* (WM), *Id4* (deep), *Mfge8* (upper). Merged tracks from N=4 biological replicates. I-K) Hybridization chain reaction (HCR) and quantification of *Tnc*, *Mfge8,* and *Id4* mean signal intensity within astrocyte cell bodies (S100b+). I) N=3 mice per genotype of either sex, n=2-6 technical replicates/mouse. No significant differences between genotypes (two-way ANOVA). K) N=3 mice per genotype of either sex, n=2-6 technical replicates/mouse. 2-way ANOVA for pial distance bin and genotype: bin (p <0.0001), genotype (p = 0.02) & bin:genotype (p = 0.005). Tukey posthoc tests: bin 1, p = 0.0321; bin7, p = 0.0355; bin 9, p = 0.0138; bin 10, p = 0.0004. For I-K, scale bar: 100 um. All values represent the mean_±_SEM.

Normalization and quality control analyses were performed for Zbtb20 and H3K4me3 CUT&RUN and their respective negative controls. Peak-calling was performed separately for each probe (Zbtb20 and H3K4me3), with each run comprising 4 target and 4 negative-control replicates. Peaks detected by at least 5 of 6 peak-calling algorithms (**Fig. S8c-d**) were selected to form consensus peak sets, which were used for downstream analyses. Peak annotation (**Fig. 6b**) revealed that promoter regions are the most highly represented genomic feature among Zbtb20-bound loci (44.3%). As expected, H3K4me3 was found almost exclusively at promoter regions (93.6% of all bound loci). We identified 3343 high-confidence Zbtb20-bound promoter regions within 1.5kb of a Transcription Start Site (TSS) (**Fig. 6c**). Most of the Zbtb20-bound promoters exhibited high H3K4me3 signal, indicating they are active promoters in at least a subset of cortical astrocytes.

In agreement with Zbtb20 regulating layer-specific gene expression in astrocytes, we identified 80 astrocyte subtype-enriched genes among the high-confidence Zbtb20-bound promoters (**Fig. 6d**). Next, to determine if δ-catenin cooperates with Zbtb20 to control astrocyte layer-specific transcriptional specialization, we compared the expression of Zbtb20-bound layer-enriched genes in *Ctnnd2*-WT and KO astrocytes (**Fig 6e**). Indeed, 45% (36 out of 80) of the Zbtb20-bound layer-enriched genes are differentially expressed in *Ctnnd2*-KO compared to WT astrocytes from snRNAseq (**Fig. 6f**). Conversely, only 14.8% (507 of 3343) of all Zbtb20 binding targets are differentially expressed in astrocytes (Fisher’s exact test: p<0.0001). Interestingly, most (24 of 36) of the layer-specific Zbtb20-bound DEGs are enriched in upper-layer astrocytes (**Fig. 6g**), suggesting that the δ-catenin and Zbtb20 cooperate to regulate upper-layer gene expression in astrocytes.

To further test this possibility, we selected 3 exemplary transcripts (**Fig. 6h**) differing with respect to Zbtb20 binding, layer of enrichment, and differential expression in *Ctnnd2*-KO compared to WT astrocytes. *Tnc* (Tenascin-C) is not bound by Zbtb20, is enriched in WM astrocytes, and is not differentially expressed in KO. *Id4* (Inhibitor of DNA-binding 4) is bound by Zbtb20, enriched in deep-layer astrocytes, and upregulated in KO. *Mfge8* (Milk Fat Globule-EGF Factor 8) is bound by Zbtb20, enriched in upper-layer astrocytes, and upregulated in KO. Hybridization chain reaction (HCR) was performed in V1 of *Ctnnd2*-WT and KO animals at P30 (**Fig. 6i-k**). The WT samples demonstrated appropriate layer enrichment of each transcript, with *Tnc* and *Id4* enriched in border regions and *Mfge8* in upper layers. Consistent with a specific role for δ-catenin and Zbtb20 cooperating to regulate upper-layer genes, we found no change in *Id4* and *Tnc* expression level or spatial distribution in *Ctnnd2-*KO visual cortex (**Fig. 6i-j**). In contrast, the laminar distribution of *Mfge8* is dysregulated, with increased expression at border regions in KO cortex (**Fig. 6k**). Taken together with the spatial transcriptomic findings, these results show that δ-catenin is required to regulate upper-layer-specific expression of Zbtb20 target genes in astrocytes.

## DISCUSSION

### δ-catenin coordinates transcriptional development of multiple cortical cell types

In this study, we identify δ-catenin as a regulator of transcriptional maturation in the developing cortex, linking cell–cell interactions to gene expression programs that shape circuit development. Previous mechanistic studies of δ-catenin have primarily focused on its roles in glutamatergic neurons, including synaptogenesis, dendritic branching, and dendritic spine maintenance [51, 52, 93, 94]. Multiple groups have documented functional neurological deficits in *Ctnnd2* knockout and truncated mutant mouse models; however, whether *Ctnnd2* plays critical roles in the development and function of other neural cell types has remained understudied [52, 54, 94, 95]. Here, by taking a holistic approach to neurodevelopment, we find that loss of δ-catenin disrupts transcriptional maturation across multiple neural cell types, with particularly pronounced effects in astrocytes. These transcriptional changes are accompanied by impaired acquisition of layer-specific astrocyte identities and prolonged ODP, consistent with a failure of circuit stabilization. Notably, these transcriptomic alterations are reminiscent of molecular signatures observed in the brains of individuals with neurodevelopmental disorders (NDDs) [1, 96].

At the mechanistic level, our findings suggest that δ-catenin operates within cadherin-based signaling pathways that mediate cell–cell interactions during cortical development. δ-catenin functions downstream of cadherins, which are key components of a “chemoaffinity code” that governs neuronal subtype-specific interactions and circuit assembly in the developing nervous system [61, 97, 98]. Glutamatergic neurons populate the cortex first and establish its laminar architecture, thereby positioning them to instruct the molecular specialization of astrocytes, oligodendroglia, and interneurons [19, 29–31, 104, 105]. Our findings imply that transcellular cadherin–catenin signaling between glutamatergic neurons and neighboring cells coordinates the transcriptional maturation of glia and interneurons.

Why astrocytes rely on neuron-derived cues for maturation remains an important question. The maturation of these glial cell types has been linked to the closure of the ocular dominance plasticity (ODP) critical period, which marks the transition from plasticity to stabilization of visual circuitry [18]. By engaging a shared cadherin–catenin-dependent mechanism, astrocytes may synchronize their developmental trajectories to underlying circuits to ensure coordinated maturation. Consistent with this model, we find that loss of δ-catenin prolongs critical-period plasticity. Together, these findings suggest that coordinated transcriptional maturation of glia and interneurons is required for the timely closure of critical periods.

### How does δ-catenin loss impact transcription?

The transcriptional changes induced by δ-catenin deletion were most pronounced in NeuN-negative cells, including astrocytes and OLs, all of which highly express the BTB/POZ TF Zbtb20. Zbtb20 was among the top upregulated genes in *Ctnnd2*-KO visual cortex and is highly enriched in glia compared to neurons, implicating it as a genetic interactor of δ-catenin with a specialized role in glial development. Given that δ-catenin is known to modulate the transcription regulatory activity of another BTB/POZ TF, Zbtb33 [99], we investigated a possible link between δ-catenin and Zbtb20-dependent transcriptional activity. We found that, in NeuN-negative cells, Zbtb20 binds to lowly methylated TSSs within CpG island promoters, and δ-catenin loss disrupts both methylation and Zbtb20 binding at these sites. At the same time, δ-catenin loss increases Zbtb20 expression and DNA binding frequency, notably enhancing its association with promoters containing the Wnt Response Element (WRE) (**Fig S7h**).

What are the mechanisms by which δ-catenin loss induces these changes in Zbtb20 binding location? One possibility is that δ-catenin translocates to the nucleus and directly interacts with Zbtb20. In non-neuronal systems, δ-catenin and the related p120-catenin have been shown to bind the closely related BTB/POZ transcription factor Zbtb33 (Kaiso) and modulate its DNA occupancy and transcriptional activity [59, 99-101]. Zbtb20 shares structural homology with Zbtb33, raising the possibility that δ-catenin may similarly influence Zbtb20-dependent transcriptional regulation. δ-catenin interacts with Zbtb33 near its DNA-binding zinc finger domains, typically disrupting its association with DNA and displacing it from target gene promoters, although it can enhance binding at select loci [59, 99-101]. By analogy, δ-catenin binding to Zbtb20 could displace it from non-specific promoters while enhancing its specificity at CpG island promoters.

Notably, loss of δ-catenin predominantly induces upregulation of Zbtb20 target genes. This could be explained by the fact that BTB/POZ TFs can act bivalently as either transcriptional repressors or activators depending on the type of cofactor complex recruited [63, 64]. Perhaps δ-catenin promotes Zbtb20 recruitment of transcriptional corepressors; conversely, in the absence of δ-catenin, Zbtb20 recruits coactivators and activates gene expression (**Fig S7h**). Each of these hypotheses presents exciting avenues for future mechanistic studies.

In addition to potential protein–protein interactions, DNA sequence and methylation context are known to influence the binding of BTB/POZ transcription factors. Zbtb33 (Kaiso) preferentially binds methylated CpG dinucleotides and functions as a methylation-dependent transcriptional repressor [88, 99, 102, 103]. In contrast, we found Zbtb20 binding in cortical glial cells is enriched at non-methylated CpG island promoters. This distinction suggests that, despite their structural similarity, Zbtb20 and Zbtb33 operate in distinct epigenetic contexts *in vivo*. Notably, δ-catenin loss disrupts this relationship, leading to a loss of the normal coupling between binding and methylation state. Together, these findings raise the possibility that δ-catenin regulates Zbtb20-dependent transcription not only through protein interactions but also by influencing the chromatin context in which Zbtb20 binds.

It is also possible that δ-catenin regulates Zbtb20 indirectly via β-catenin. When complexed with cadherins and other catenins, including α- and δ-catenin, β-catenin is sequestered away from the nucleus [104]. Phosphorylation-dependent dissociation of δ-catenin destabilizes the cadherin-catenin membrane complex [29, 105], promoting β-catenin nuclear translocation and transcriptional activity [106-108]. Consistent with this, bulk RNA sequencing of P21 visual cortex showed a strong upregulation of Wnt-related and stemness-like transcriptomic signatures in the absence of δ-catenin (**Fig. 1h-n**). Therefore, global loss of δ-catenin may also destabilize the cadherin-catenin complex and promote β-catenin nuclear shuttling. When β-catenin is in the nucleus, it interacts with WRE-binding TCF/LEF repressive TFs, either promoting their recruitment of coactivators [109] or displacing them from DNA [110, 111]. Therefore, increased β-catenin nuclear shuttling in *Ctnnd2-*KO may displace TCF/LEF TFs and leave WRE promoters open for Zbtb20 binding (**Fig. S7h**). Alternatively, nuclear β-catenin could promote cooperative WRE binding and coactivator recruitment by Zbtb20 and TCF/LEF TFs. In either case, δ-catenin loss would primarily activate gene expression because β-catenin translocation to the nucleus would relieve TCF/LEF TF-dependent repression. Future studies are needed to disentangle the direct and indirect mechanisms by which δ- and β-catenin control Zbtb20’s transcription regulatory activity.

### δ-catenin links astrocyte morphogenesis to layer-specific transcriptional identity

Astrocytes were profoundly affected by δ-catenin loss, exhibiting disrupted layer-specific gene expression. Astrocytes populate the cortex during the late embryonic to early postnatal period, after glutamatergic neurons establish their six-layer architecture [7]. Astrocytes undergo layer-specific molecular specialization after colonizing the parenchyma, suggesting they rely on local cues to decide their identity. Indeed, previous studies have demonstrated that genetic disruption of glutamatergic neuronal layering results in a loss of astrocyte layer-specific transcriptional specialization [8, 10]. Remarkably, our results reveal δ-catenin loss disrupts layer-specificity in astrocytes (**Fig. S4a-d**) without causing a major change in glutamatergic neuron layer organization (**Fig. 4g-i**). One possible explanation for this finding is that δ-catenin transduces layer-specific cues from glutamatergic neurons to astrocytes.

One outstanding question is how δ-catenin-based contact conveys layer-specific information, given that δ-catenin itself is expressed across all cortical layers in astrocytes and neurons [29]. We propose that δ-catenin derives layer specificity from its interactions with cadherins, a large superfamily of transmembrane cell adhesion molecules with layer-restricted expression patterns in CNS neurons [97, 98, 112]. In agreement, we previously found that δ-catenin, through cadherin-based interactions with neurons, controls layer-specific astrocyte morphogenesis [29]. These studies link astrocyte morphological development to layer-specific molecular specialization. Therefore, it is plausible that neuronal cadherin-based contact instructs δ-catenin transcriptional signaling in astrocytes in a layer-specific manner as well (**Fig. S8e**). Future studies linking cadherins to layer-specific astrocyte gene expression are needed to explore this exciting possibility.

### Zbtb20 regulates astrocyte layer identity in a δ-catenin-dependent manner

Our single-nucleus and spatial transcriptomic experiments revealed that δ-catenin loss increases the proportion of astrocytes in an upper-layer-like transcriptional state. Moreover, a significant proportion of Zbtb20-bound upper-layer astrocyte genes were upregulated in *Ctnnd2*-KO. Based on these results, our working model is that δ-catenin and layer-specific neuronal cadherins together control Zbtb20 in astrocytes of different cortical layers (**Fig. S8e**). In the absence of stable cadherin-based adhesion with a neighboring neuron, Zbtb20 binds to promoters of upper-layer astrocyte genes and exerts a positive (activating) effect on gene expression. Once stable cadherin-based adhesion is established, changes in cadherin or catenin phosphorylation trigger δ-catenin release from the transcellular cadherin complex, as has been previously demonstrated [35]. δ-catenin shuttles to the nucleus, where it inhibits Zbtb20-mediated activation of layer-enriched transcripts via mechanisms described in the previous section. Such a model could explain how certain genes (e.g., *Mfge8*) are activated in upper-layer astrocytes, which are surrounded by N-cadherin-low neurons, while they are repressed in deep-layer astrocytes, which are surrounded by N-cadherin-high neurons. Moreover, this model explains why astrocytes in deep layers and border regions adopt an upper/mid-layer-like transcriptional program in *Ctnnd2*-KO. Fluctuating extracellular Wnt abundance, which controls β-catenin nuclear signaling, can also intersect δ-catenin/Zbtb20 signaling to provide further spatial and temporal specificity. In addition to Wnt and cadherins, various other signaling axes exhibit layer-specific gradients, creating a highly complex multimodal system. The findings presented here provide inroads to begin dissecting both catenin-dependent and independent mechanisms controlling layer-specific gene expression in astrocytes.

### Zbtb20’s sequential functions during astrocyte subtype specification

Although there is strong evidence that local cues instruct astrocyte layer specification, it is also possible that subtypes are partially specified within progenitor pools. In agreement, lineage tracing combined with single-cell transcriptomics shows that two molecularly distinct astrocyte-producing RGC populations coexist from the outset of cortical astrocytogenesis [56]. Strikingly, *Zbtb20* was highly enriched in one of the two RGC populations, which gave rise to astrocytes closely resembling our upper-layer astrocyte cluster [56]. In light of our findings, these results suggest that Zbtb20 begins to push astrocytes toward an upper-layer-like transcriptional program during the pre-mitotic progenitor phase. Indeed, prior to our study, Zbtb20 was primarily studied for its role in astrocytogenesis, where it controls initial differentiation and fate determination events [65, 66]. Our astrocyte-specific CUT&RUN dataset provides a comprehensive survey of Zbtb20 target genes in postnatal astrocytes, establishing a link between Zbtb20 and astrocyte layer specification. Profiling these newly identified Zbtb20 targets throughout development will be a powerful approach to address the longstanding debate of when and how astrocyte layer-specific subtypes arise.

### A multicellular role for δ-catenin in neurodevelopmental disorders

Although glutamatergic neurons are the fundamental drivers of cortical activity and long-range output, they are not alone competent to form and maintain functional synaptic networks. Indeed, many essential features of a well-constructed cortical network—sparse encoding, long-range conduction efficiency, and critical-period plasticity—depend on non-glutamatergic cell types [113-117]. However, it is unknown how these multiple cell types coordinate their cellular maturation to set the pace and sequence of cortical network refinement. Here, we propose that δ-catenin, encoded by the human disease-linked *CTNND2* gene, performs this function by coupling glial development to their glutamatergic neuron neighbors. Remarkably, loss of δ-catenin both decouples cellular maturation and prolongs experience-dependent plasticity, suggesting the two processes are interdependent. Though we assayed plasticity in the visual circuit, other cortical areas may also exhibit mistimed cellular maturation and circuit plasticity in *Ctnnd2*-KO. Indeed, each of the behavioral phenotypes previously observed in *Ctnnd2*-KO mice, social deficits, fear learning, repetitive behaviors, memory deficits, and sleep-wake disturbances, relies on cortical circuitry that undergoes a distinct period of experience-dependent plasticity [52-54, 95, 118, 119]. Correspondingly, we propose that plasticity deficits across various brain regions may explain the wide range of neurodevelopmental and neurodegenerative disorders observed in cases of human *CTNND2* mutations. The findings presented here provide strong rationale to study how astrocytes and oligodendrocytes, which are profoundly affected by *Ctnnd2* loss, contribute to these other behavioral phenotypes.

### Limitations of the study

Due to the use of a global deletion model, we cannot definitively say whether transcriptional dysregulation in the absence of δ-catenin is cell-intrinsic, cell-extrinsic, or a transcellular effect. We previously used astrocyte- and neuron-specific *in vitro* and *in vivo* knockdown models to show that δ-catenin impacts astrocyte morphology through a transcellular adhesion-based mechanism. Therefore, we propose that δ-catenin is also required in both cell types to instruct astrocyte transcription. In the future, cell-type-specific manipulations can test this hypothesis.

We did not pinpoint which cortical cell type is responsible for the extension of ODP we observed in the *Ctnnd2*-KO mouse model. Our interpretation is that all affected neural cell types could be involved: indeed, layer 4 neurons, inhibitory interneurons, oligodendroglia, and astrocytes have all been linked to the closure of the critical period [20, 24, 26, 27, 120, 121]. Future studies targeting *Ctnnd2* in each cell type are needed to dissect the respective contributions of these cell types to the ODP phenotype.

Our study does not differentiate between the possibilities that the ODP critical period is prolonged or remains open indefinitely. Interestingly, δ-catenin was initially discovered as an interactor of the Alzheimer’s gene presenilin-1 [122], and *CTNND2* mutations are linked to neurodegenerative disease in humans [123]. Therefore, future studies investigating circuit plasticity and non-neuronal cell development in older mice will enhance our understanding of δ-catenin function and its contributions to human disease.

## ACKNOWLEDGMENTS

This work was supported by Adelson Medical Research Foundation Award and National Institutes of Health grants to C.E., U19NS123719 and NS102237, and a grant from the Duke Precision Genomics Collaboratory. G.S. was supported by NIGMS T32GM007171-47, NIGMS T32GM145449-01, and AHA 25PRE1357000. J.T.S. was supported by NIH funding (F31NS134252), G.A.R. was supported by the Duke Summer Neuroscience Program, and J.D. was supported by the Duke Dean’s Summer Research Fellowship. Biorender was used to generate elements of figures 5 and 6. We thank the Eroglu lab, Dr. Nicole Calakos, Dr. Chantell Evans, Dr. Scott Soderling for their valuable feedback. We thank Dr. Debra Silver, Dr. Brooke D’Arcy, Dr. Rohit Singh, and Dr. Anne West for their critical reading of the manuscript. C.E. is an HHMI Investigator.

## EXPERIMENTAL MODEL AND STUDY PARTICIPANT DETAILS

### Mice

All mice in this study were used in accordance with the Institutional Animal Care and Use Committee (IACUC) and the Duke Division of Laboratory Animal Resources (DLAR) oversight (IACUC Protocol Numbers A103-23-04 and A110-23-04). Mice were housed in groups of 2–5 per cage with a 12 h dark-light cycle and free access to food and water. Wild type CAG-Sun1/sfGFP reporter mice (Strain #030952, RRID:IMSR_JAX:030952) and wild-type C57BL6/J mice (Strain #000664, RRID:IMSR_JAX:000664) were obtained through MMRRC. WT CD1 mice used for PALE and in utero electroporation experiments were purchased through Charles River Laboratories (RRID:IMSR_CRL:022). Ctnnd2 knockout (Ctnnd2-KO) mice were generated in collaboration with the Duke Transgenic Mouse Facility using CRISPR-EZ protocol from Modzelewski et al [124]. Briefly, single guide RNAs (sgRNAs) specific to intronic regions flanking exons 4 and 5 (Table 1) were synthesized by in-vitro transcription of a short DNA template.

**Table 1.**
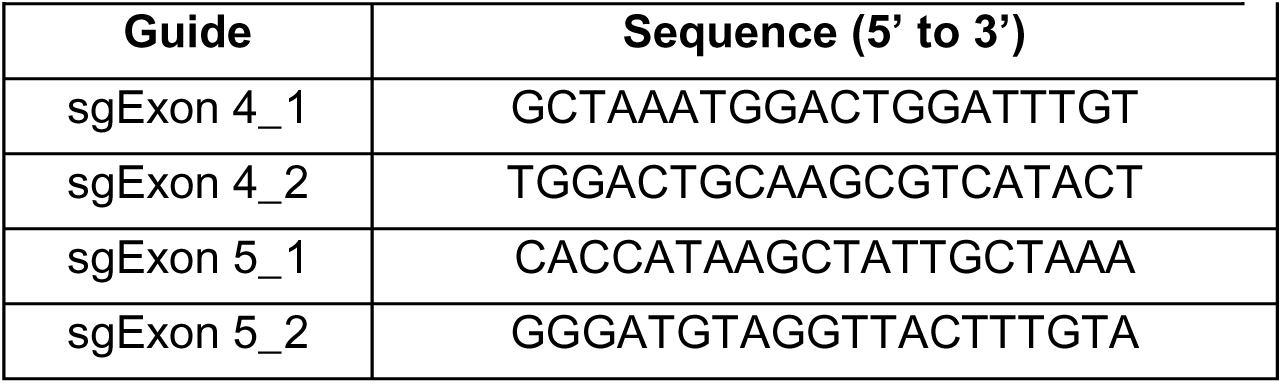
List of sgRNAs used in targeted exon deletion of Ctnnd2-KO mice.

After donor superovulation, pronuclear-stage embryos were collected, cultured, and electroporated with a ribonucleoprotein (RNP) complex prepared by incubating sgRNAs with Cas9 protein (IDT, Alt-R Cas9 v3 Cat#1081059) at 37°C for 10 minutes. Viable embryos were cultured and genotyped ex vivo to test the sgRNA efficiency. Successful embryos were oviduct transferred to pseudopregnant C57BL6/J females. After birth, founders were screened by PCR of genomic DNA using custom primers (Table 2).

**Table 2.**
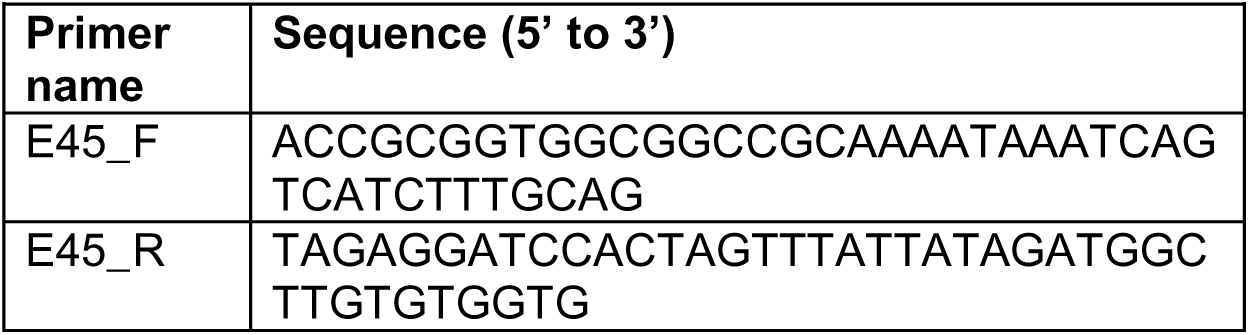
List of primer sequences used in allelic subcloning.

PCR products were subcloned into pL253 at the NotI-SpeI restriction sites. From each founder, independent subclones were sequenced by Sanger sequencing to identify the targeted mutation and select putative positive chimeras. For line maintenance, heterozygous males were crossed to wild-type C57BL6/J females from Jackson Laboratories, and F1 heterozygous offspring were used as experimental breeders. For experimental breeding pairs, mice were bred in a heterozygous x heterozygous paradigm to produce a mix of wild-type, knockout, and heterozygous littermates. Mice were genotyped using genomic PCR with custom primers (Table 3).

**Table 3.**
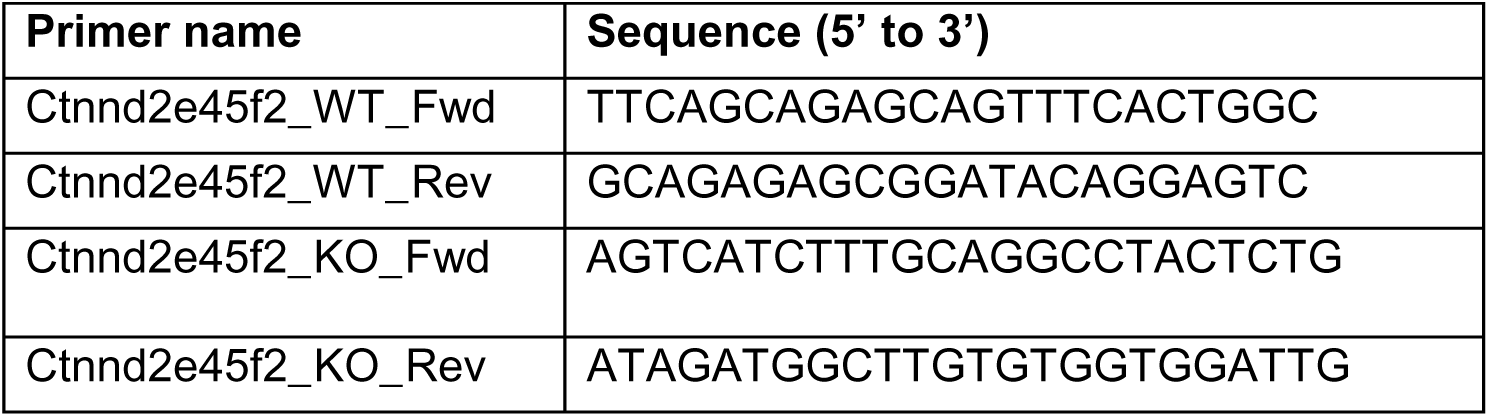
List of primer sequences used in Ctnnd2-KO genotyping.

P21, P28, P30, P49, P50, and P53 mice were used for experiments or as specified in the text and figure legends. Age-matched wild-type were used as controls in all experiments except monocular deprivation, where both wild-type and heterozygous littermates were used as controls. For all experiments both male and female mice were used with sex ratios equivalent to or not significantly different from 1:1. No significant differences in outcome measures between males and females were detected.

## METHOD DETAILS

### SURGICAL PROCEDURES

#### Monocular Deprivation

P28 or P49 mice were anesthetized using 5% isoflurane in oxygen. Animals were visually monitored for respiration and signs of distress. Once deeply anesthetized, mice were maintained anesthetized with 1.5% isoflurane in oxygen. Prior to surgery, mice were given meloxicam (2 mg/kg) subcutaneously as pre-operative analgesic (0.4mL). Depilatory cream was used to remove fur around the eye. Top and bottom eyelash bed was removed from one eye. A single horizontal mattress suture was used to seal the eyelids shut and Vetbond was used to aid with wound closure. Following eyelid closure, mice were monitored on heating pad until they successfully recovered and were returned to home cage. Mice used for normal rear condition were collected ≤12 hours following surgery, and mice used for monocular deprivation condition were deprived for 3-4 days. These mice received meloxicam (2 mg/kg) subcutaneously once daily for the first 48-72 hrs as post-surgical analgesia (0.4mL first day, 0.25 mL second day, 0.25 mL as needed third day). Eyelid closure was assessed each day to ensure wound healing and full closure of eyelid. If needed, additional Vetbond was added to protect suture.

#### Adeno-associated virus (AAV) production and administration

Purified AAV was produced in house and in collaboration with the Duke University Viral Vector Core facility. Briefly, HEK293T cells were co-transfected with 3 plasmids: pAd-DELTA F6 (plasmid No. 112867; Addgene), serotype plasmid AAV PHP.eB (plasmid No. 103005), and AAV plasmid (GEARBOCS, Addgene #218181). After transfection, cells were lysed and the replicated viral vectors were collected, purified and stored at −80°C. The viral titer for each vector was calculated by real-time PCR. To induce astrocyte-specific expression of GFP on the nuclear envelope, CAG-Sun1/sfGFP mice were injected intracortically with GEARBOCS AAV (AAV-U6-sgRNA-GfaABC1d-Cre-6xmiRT) between postnatal days 3 and 4 (P3-4). Pups were anesthetized by hypothermia and 1μl of concentrated AAV vector was injected bilaterally at 1 rostral and 1 caudal site (4 sites total) using a Hamilton syringe. The pups were monitored until recovered on a heating pad and returned to the cage for parental care.

### MOUSE TISSUE PREPARATION

#### Immunohistochemistry & smFISH

Mice were anesthetized with 200 mg/kg tribromoethanol (Avertin) and perfused with 1X TBS or TBS/heparin and 4% PFA. Brains were collected and post-fixed in 4% PFA for 1 overnight, then cryoprotected in 30% sucrose in 1X TBS. Brains were frozen in solution of 2 parts 30% sucrose and 1 part O.C.T. (TissueTek) and stored at −80°C until time of sectioning. For immunohistochemistry experiments, coronal tissue sections of 40 μM (P30 staining for oligodendrocyte subtypes) or 30 μM (P21 staining for neural subtypes, neuronal subtypes, and Zbtb20 expression) thickness were collected and stored in a 1:1 solution of TBS/glycerol at −20°C. For smFISH experiments, 20 μM sections were directly mounted on Superfrost Plus slides and stained and imaged within 1 week.

#### *Arc* induction

All mice for *Arc* induction were collected between 4:00 and 6:30am at the end of the animal’s dark cycle (active phase). Mice stimulated monocularly through their open eye with white light for ∼ 30 minutes just prior to brain collection. Mice were then anesthetized with 200 mg/kg tribromoethanol (Avertin) and transcardially perfused with 1X TBS followed by 4% PFA. Brains were collected and post-fixed in 4% PFA for 1 overnight, then cryoprotected in 30% sucrose in 1X TBS. Brains were frozen in solution of 2 parts 30% sucrose and 1 part O.C.T. (TissueTek) and stored at −80°C until time of sectioning. 20 μM sections were directly mounted on Superfrost Plus slides (VWR) and stained and imaged within 1 week.

#### Xenium

All mice for Xenium were collected between 9:00 and 10:00am. The morning of collections, 10% NBF (pH 7.2) was prepared fresh: combine 10 mL formalin (37% formaldehyde) with 90 mL diH2O, 0.65 g dibasic/anhydrous Sodium Phosphate (Na2HPO4) and 0.4 g monobasic NaH2PO4, adjust pH to 7.2.Mice were anesthetized with 200 mg/kg tribromoethanol (Avertin) and transcardially perfused with 1X PBS followed by 10% neutral buffered formalin (NBF). Brains were collected and dehydrated in 70% ethanol until paraffin embedding. Paraffin was performed by the Duke BioRepository and Precision Pathology Center (BRPC), according to standard protocol. A single coronal section containing posterior neocortex (bregma ∼-2.5) was collected for each sample at 5 μm thickness onto Xenium slides (10X Genomics 1000659), dried at 42C for 3 hours and stored at room temperature in a desiccator. Each sample (1 WT female, 1 WT male, 1 KO female, 1 KO male) was mounted on a separate slide. Two sex-matched littermate pairs were processed in parallel for all steps of the procedure.

#### Isolation and enrichment of non-neuronal nuclei from mouse brain

The following protocol was used to isolate non-neuronal nuclei from juvenile (P28) *Ctnnd2*-WT and KO mice prior to downstream CUT&RUN and snRNA-seq experiments. Mice were deeply anesthetized with 200 mg/kg tribromoethanol (Avertin), acutely decapitated, and the brain was immediately extracted. For CUT&RUN, each cortical hemisphere was bluntly dissected from underlying structures using forceps. For snRNAseq, visual cortex was microdissected by using a scalpel to make 2 transverse and 2 longitudinal cuts on the dorsal surface of the brain (corresponding to the anterior, posterior, medial, and lateral borders) then bluntly separated from underlying structures using forceps. Tissue samples were immediately flash-frozen in liquid nitrogen and stored at -80°C. Nuclei were extracted and magnetically enriched from frozen cortex samples (single full cortical hemisphere for CUT&RUN or bilateral visual cortex for snRNA-seq) using Miltenyi Biotec reagents and technology, according to manufacturer protocol. Briefly, tissue was transferred to gentleMACS C Tube containing nuclei extraction buffer (Miltenyi Nuclei Extraction buffer), C tube was placed on genteMACS Dissociator and covered with gentleMACS Octo Cooler, and the preset gentleMACS program “4C_nuclei_1” was run. Crude nuclei were strained using MACS SmartStrainer (70 µm), centrifuged 300 × g at 4 °C for 5 minutes, resuspended in ice-cold resuspension buffer (Miltenyi Nuclei extraction buffer, DPBS), and strained again using MACS SmartStrainer (30 µm). Anti-nucleus Microbeads were added to crude nuclei (50 uL for bilateral visual cortex, 75 uL for unilateral whole cortex) and incubated at 4 °C for 15 minutes on a nutator. Magnetic separation was performed using LS columns and MACS Separator according to standard manufacturer protocol. Magnetically enriched nuclei were centrifuged 300 × g at 4 °C for 5 minutes, resuspended in 50 uL Alexa Fluor 488-conjugated anti-NeuN antibody (MAB377X, diluted 1:1000 in DPBS with 1% BSA), and incubated for 1 hour on ice in the dark. Finally, samples were washed in 450 uL of sorting buffer (DPBS with 1% BSA), centrifuged 300 × g at 4 °C for 5 minutes, resuspended in sorting buffer, and strained using MACS SmartStrainer (30 µm) immediately prior to sorting. For snRNAseq, RNase inhibitor was added to buffers at a final concentration of 0.2 U/uL. For CUT&RUN, Roche complete protease inhibitor cocktail was added to buffers at a final concentration of 1X.

#### Isolation and enrichment of astrocyte nuclei from mouse brain

The following protocol was used to isolate astrocyte nuclei from juvenile (P21) CAG-Sun1/sfGFP mice with AAV-induced expression of astrocyte-specific Cre (GEARBOCS, Addgene #218181) prior to downstream CUT&RUN experiments. Mice were deeply anesthetized with 200 mg/kg tribromoethanol (Avertin), acutely decapitated, and the brain was immediately extracted. Each cortical hemisphere was bluntly dissected from underlying structures using forceps. Tissue samples were immediately flash-frozen in liquid nitrogen and stored at -80°C. Nuclei were extracted from a single frozen cortical hemisphere using Miltenyi Biotec reagents and technology without magnetic enrichment. Briefly, tissue was transferred to gentleMACS C Tube containing nuclei extraction buffer (Miltenyi Nuclei Extraction buffer), C tube was placed on genteMACS Dissociator and covered with gentleMACS Octo Cooler, and the preset gentleMACS program “4C_nuclei_1” was run. Crude nuclei were strained using MACS SmartStrainer (70 µm), centrifuged 300 × g at 4 °C for 5 minutes, resuspended in ice-cold resuspension buffer (Miltenyi Nuclei extraction buffer, DPBS), and strained again using MACS SmartStrainer (30 µm) immediately prior to sorting. A protease inhibitor cocktail was added to buffers at a final concentration of 1X.

#### Flow cytometry

Fluorescence-activated nuclei sorting (FANS) was performed in the Duke Cancer Institute Flow Cytometry Shared Resource (DCI-FCSR) using a Sony MA900 Multi-Application Cell Sorter with 1X PBS for sheath fluid at room temperature in 2-way (snRNAseq) or 1-way (CUT&RUN) purity mode. A 70 μm nozzle was used for these experiments with sheath pressure set to 40 PSI and event rate was maintained at approximately 1000-1600 events per second. Exclusion of debris was performed first using forward and side scatter pulse area parameters (FSC-A and BSC-A), followed by exclusion of aggregates using pulse height (FSC-A and BSC-H), before gating populations based on 7AAD and GFP or Alexa Fluor 488 fluorescence. 7-AAD was excited with 561 nm laser, and GFP and AF488-NeuN were excited with 488 nm laser. To isolate both neuronal and non-neuronal populations (snRNAseq), nuclei with supra- and sub-threshold FITC signal (NeuN+ and NeuN-, respectively) were collected separately. To isolate only the non-neuronal population (WT vs KO glia CUT&RUN), nuclei devoid of FITC signal (NeuN-) were collected. To isolate the astrocyte population (WT astrocyte CUT&RUN), nuclei positive for FITC signal (GFP+) were collected. Nuclei were purified using a one-drop single-cell sort mode (for counting accuracy); these were directly deposited into a 1.5 ml Eppendorf pre-coated with 250-500 uL additional buffer. Data were analyzed using MA900 software (Sony). FSC was plotted in linear scale, while SSC and fluorescence in logarithmic scale to facilitate gating relying on strong side scatter and respective fluorochrome signals.

#### Bulk RNA sequencing tissue preparation

Tissue samples were harvested from WT and *Ctnnd2*-KO adult mice (6 animals per genotype, including 3 males and 3 females) at postnatal day 21. Animals were anesthetized with 200 mg/kg tribromoethanol (avertin) and acutely decapitated. The brain was immediately extracted, 4 transverse cuts were made with a clean, chilled razor: 1) Just anterior to the cerebellum, 2) Bisecting the cortex just anterior to half anterior-posterior distance, 3) Bisecting the anterior half of the cortex, 4) Bisecting the posterior half of the cortex. The cerebellum was discarded, leaving 4 coronal sections of cortex, numbered 1-4 in anterior-posterior order. Prefrontal cortex was isolated from section 2, and visual cortex from section 4, in each case using forceps to bluntly dissect each cortical hemisphere from underlying structures using forceps. Tissue samples were immediately flash-frozen in liquid nitrogen and stored at -80°C until RNA extraction.

#### Protein extraction

P14 Ctnnd2 E4/5 or Ctnnd2 E6 WT, HET and KO sex-matched littermate trios were anesthetized by injection of 200 mg/kg tribromoethanol (avertin) and perfused with 1X TBS. The cortex was quickly dissected out on ice, divided into hemispheres, snap frozen in liquid nitrogen, and stored at -80°C. 2ml dounce tissue grinders (Sigma-Aldrich #D8938) were pre-cooled on ice prior to tissue lysis. 1 hemisphere was dounced in 1 mL of ice-cold lysis buffer (25 mM HEPES (Sigma-Aldrich #H0887), 150 mM KCl, 1.5 mM MgCl2, 10% glycerol, 1X Protease Inhibitor (Roche cOmplete Protease Inhibitor Cocktail #4693132001), 1% NP-40 (Thermo Scientific #28324)). The tissue suspension was vortexed for 10 seconds and returned to ice for 5 minutes a total of 3 times, then centrifuged at 18,000g for 10 minutes at 4°C to pellet non-solubilized debris. The supernatant containing solubilized protein was collected. Protein concentration was measured by a Micro BCA assay (Thermo Scientific #23235 and denatured in 2X Laemmli Sample Buffer (Bio-rad #1610737) containing 5% beta-mercaptoethanol for 45mins at 45°C. Samples were stored at -80°C.

### BIOCHEMICAL ASSAYS

#### Western blot

Protein samples were loaded into a 4–15% Mini-PROTEAN® TGX Stain-Free™ Protein Gel (Bio-rad #4568084) and ran at 100V for 100 minutes. The gel was then transferred to a PVDF membrane at 100V for 7 minutes using a Power Blotter (Invitrogen). Membranes were then washed once with 0.1% Tween20 in 1X TBS (TBST) and blocked for 1 hour in 1% Bovine Serum Albumin in 0.1% TBST. Membranes were incubated in primary antibody in 0.1% TBST overnight at 4°C. The following combinations of primary antibodies were used: δ-catenin (Abcam #ab184917) and GAPDH (Abcam #ab8245), δ-catenin (BD Biosciences #611536) and GAPDH (Abcam #ab9485), or δ-catenin (Santa-Cruz #81793) and GAPDH (Abcam #ab9485). The next day, membranes were washed three times with 0.1% TBST, incubated in secondary antibody (LI-COR) for 2 hours at room temperature, washed 3 times with 0.1% TBST, and imaged on an Odyssey Infrared Imaging System on the Image Studio software.

### HISTOLOGY

#### Nuclear staining for neural cell types

Three coronal sections containing the visual cortex were selected from each sample. Sections were permeabilized by washing 3 x 10 min in 1 x TBS containing 0.2% Triton X-100 (TBST) at room temperature (RT) on a shaker. Sections were blocked in 10% NGS diluted in TBST for 1h at RT on a shaker. Following blocking, the sections were incubated with primary antibody mix diluted in 10% NGS in TBST (Rb Sox9, 1:2000; Ms IgG1 NeuN, 1:1000; Ms IgG2a Olig2, 1:500) for 2 overnights at RT with shaking. After primary incubation, sections were washed in 3 x 10 min in TBST at RT on a shaker, then incubated in Alexa Fluor conjugated secondary antibody mix containing goat anti-Rb 488, goat anti-Ms IgG1 594, and goat anti-Ms IgG2a 647 all diluted 1:200 in 10% NGS in TBST for 3 hours at RT in the dark with shaking. After this incubation, the sections were washed 3 x 10 min with TBST, with the addition of DAPI (1:10,000) in the second wash. Sections were then mounted in TBS onto glass slides using a homemade mounting media (90% glycerol, 20 mM Tris pH 8.0, 0.5% n-propyl gallate) and sealed with nail polish.

#### Nuclear staining for neuronal subtypes

To optimize nuclear signal over background, Heat Induced Epitope Retrieval (HIER) was required. Sections were first mounted on Superfrost Plus slides (VWR) and baked for 10 minutes on a slide warmer set to 60°C. Slides were immersed 10 min in 1X PBS at room temperature in a slide mailer on a shaker. Slides were transferred to a coplin jar with Sodium Citrate Buffer (10mM Sodium Citrate, 0.05% Tween 20, pH 6.0) and placed in a boiling water bath for 15 min immersion. The coplin jar containing slides was removed from the water bath and allowed to cool for 10 minutes, then slides were transferred to fresh 1X PBS at RT for additional 10 min of gradual cooling. Slides were next transferred to slide mailers for 3 x 5 min immersions in 1X PBS, followed by permeabilization in 1x PBS containing 0.25% Triton X-100 (PBST) for 10 min at RT on a shaker. Sections were blocked in 5% NGS diluted in 1X PBS for 1h at RT. Following blocking, the sections were incubated with primary antibodies diluted in 5% NGS in 1X PBS (Rt Ctip2 diluted 1:500, Ms IgG2b ROR-β 1:150, and Rb Cux1 1:500) for 2 overnights at 4°C. After primary incubation, sections were immersed 3 x 10 min in 1X PBS at RT, incubated with Alexa Fluor conjugated secondary antibodies, all diluted 1:400 in 5% NGS in 1X PBS (goat anti-Rt 488, goat anti-Ms IgG2b 594, and goat anti-Rb 647) for 2 hours at RT in the dark. After this incubation, the sections were immersed 3 times for 10 min in 1X PBS at RT, with the addition of DAPI (1:10,000) in the second wash. Sections were then mounted using a homemade mounting media (90% glycerol, 20 mM Tris pH 8.0, 0.5% n-propyl gallate) and sealed with nail polish.

#### Staining for oligodendrocyte subtypes

##### For OPCs and MOLs

Three coronal sections containing the visual cortex were selected from each sample. Sections were placed in 1X L.A.B. solution for 5 minutes at room temperature. Sections were then placed in 3 x 10 min immersions in 1X PBS on a shaker at room temperature. Sections were blocked in PBS containing 10% normal goat serum and 0.2% Triton X-100 for one hour on a shaker at room temperature. Overnight, sections were placed in a primary antibody solution containing Rt Pdgfra (dilution 1:200), Ms IgG2a Olig2 (dilution 1:500), and R ASPA (dilution 1:500) in PBS with 10% normal goat serum and 0.2% Triton X-100 at 4°C. The following day, sections were washed in 3 x 10 min immersions in PBS with 0.2% Triton X-100 on a shaker at room temperature. Sections were incubated in secondary antibody solution containing Alexa Fluor conjugated antibodies GT anti-RB 488 (dilution 1:250), GT anti-RT 594 (dilution 1:250) and GT anti-MS IgG2a 647 (dilution 1:250) in 0.2% Triton PBST on a shaker at room temperature for one hour. Next, the sections were placed in 3 x 10 min immersions in PBS containing 0.2% Triton X-100 on a shaker at room temperature, the second wash containing DAPI (dilution 1:10000). Finally, sections were placed on glass microscope slides, dried, and mounted with homemade mounting media (90% glycerol, 20 mM Tris pH 8.0, 0.5% n-propyl gallate) and sealed with nail polish.

##### For NFOLs and MFOLs

Three coronal sections containing the visual cortex were selected from each sample. Sections were washed in 3 x 10 min immersions in 1X PBS on a shaker at room temperature. Sections were blocked in PBS containing 10% normal goat serum and 0.2% Triton X-100 for one hour on a shaker at room temperature. Sections were incubated overnight in a primary antibody solution containing Ms IgG1 Bcas1 (dilution 1:300), Ms IgG2a Olig2 (dilution 1:500), and Ms IgG2b MBP (dilution 1:500) in PBS with 10% normal goat serum and 0.2% Triton X-100 at 4°C. The next day, sections were placed in 3 x 10 min immersions in PBS containing 0.2% Triton X-100 on a shaker at room temperature. Next, sections were incubated in secondary antibody solution containing Alexa Fluor conjugated antibodies Gt anti-Ms IgG1 594 (dilution 1:250), Gt anti-Ms IgG2b 488 (dilution 1:250), and Gt anti-Ms IgG2a 647 (dilution 1:250) in 0.2% Triton PBST on a shaker at room temperature for one hour. Next, the sections were placed in 3 x 10 min immersions in PBS containing 0.2% Triton X-100 on a shaker at room temperature, the second wash containing DAPI (dilution 1:10000). Finally, sections were placed on glass microscope slides, dried, and mounted with homemade mounting media (90% glycerol, 20 mM Tris pH 8.0, 0.5% n-propyl gallate) and sealed with nail polish.

#### smFISH for detection of *Arc*

Custom-made probes for *Arc* were acquired from Molecular Instruments. HCR RNA-FISH (v3.0) buffers were purchased from Molecular Instruments and used according to a modified version of the manufacturer protocol. Briefly, 20 μm sections of visual cortex from P29 (juvenile normal-reared), P32 (juvenile monocularly deprived), P50 (adolescent normal-reared), or P53 (adolescent monocularly deprived) Ctnnd2-WT and Ctnnd2-KO brains were directly mounted onto glass slides. All sections were kept at −80°C until use. First, sections were thawed to room temperature and baked for 10 minutes on a slide warmer set to 60°C. Slides were then postfixed in 500 μl of 4% PFA for 15 min at 4°C and progressively dehydrated by 4 x 5 min immersions in sequentially increasing concentrations of EtOH diluted with UltraPure H2O (50%, 70%, 100%, 100%). To prime the sections for RNA probe hybridization, slides were immersed 2 x 5 min in 1X PBS at room temperature, followed by HCR™Probe Hybridization Buffer (v3.0) for 10 min at 37°C. 100 μl of 8 nM probe solution was applied to the sections for incubation overnight in a 37°C humidified chamber. The following day, the sections were incubated in a mixture of probe wash buffer and 5× sodium chloride sodium citrate containing 0.1% Tween 20 (5× SSCT) at 37°C to remove the probe excess. Each incubation had an increasing concentration of 5× SSCT, ending with a final incubation of 100% 5× SSCT. Next, 200 μl of HCR™ Amplifier Buffer (v3.0) was added to the sections for 30 min at room temperature. Snap-cooled hairpins specific to Arc (B1594) were mixed with the amplification buffers resulting in a 120 nM hairpin solution. The sections were incubated with 100 μl of this mixture at room temperature overnight. Finally, the sections were immersed 5× SSCT for a total of 65 min to remove excess hairpins. Sections were dried, and mounted with homemade mounting media (90% glycerol, 20 mM Tris pH 8.0, 0.5% n-propyl gallate) and sealed with nail polish.

#### Multiplexed immunohistochemistry and smFISH for detection of layer-specific astrocyte genes

Custom-made probes for *Mfge8, Tnc,* and *Id4* were acquired from Molecular Instruments. HCR Gold RNA-FISH buffers were purchased from Molecular Instruments and used according to a modified version of the manufacturer protocol. Briefly, 20 μm sections of visual cortex from P30 *Ctnnd2*-WT and *Ctnnd2*-KO brains were directly mounted onto glass slides. All sections were kept at −80°C until use. First, sections were thawed to room temperature and baked for 10 minutes on a slide warmer set to 60°C. Slides were then postfixed in 500 μl of 4% PFA for 15 min at 4°C and progressively dehydrated by 4 x 5 min immersions in sequentially increasing concentrations of EtOH diluted with UltraPure H2O (50%, 70%, 100%, 100%). To prime the sections for RNA probe hybridization, slides were immersed 2 x 5 min immersions in 1X PBS at room temperature, followed by HCR™ HiFi Probe Hybridization Buffer for 10 min at 37°C. 100 μl of 16 nM probe solution (Mfge8 + Tnc or Mfge8 + Id4) was applied to the sections for incubation overnight in a 37°C humidified chamber. The following day, the sections were incubated 4 x 15 min in HCR™ HiFi Probe Wash Buffer at 37°C to remove the probe excess. Next, 200 μl of HCR™ Gold Amplifier Buffer was added to the sections for 30 min at room temperature. Snap-cooled hairpins specific to *Mfge8* (X8594) and *Tnc* (X2647) or *Mfge8* (X8594) and *Id4* (X2647) probes were mixed with the amplification buffers resulting in a 60 mM hairpin solution. To each solution, rabbit anti-S100beta primary antibody (1:200) was added. The sections were incubated with 100 μl of this mixture at room temperature overnight. The sections were immersed 4 x 15 min in HCR™ Gold Amplifier Wash Buffer at room temperature to remove excess hairpins. Finally, sections were immersed 2 x 5 min in 1x PBS containing 0.2% Tween 20 (PBST). Sections were incubated in Alexa Fluor 488 conjugated goat anti-rabbit secondary antibody diluted 1:250 in PBST with DAPI 1:10000. Finally, sections were dried and mounted with a homemade mounting media (90% glycerol, 20 mM Tris pH 8.0, 0.5% n-propyl gallate) and sealed with nail polish.

#### Xenium

Following tissue collection, Xenium slides were processed by the Duke Molecular Genomics Core (MGC), according to 10X Xenium protocol. A total of 346 genes were included in the analysis, including the 247-gene mouse brain gene expression panel (1000462) plus a 99-gene custom addon panel specific to our study (1000651, **Table S5**). Gene panel probe hybridization occurred overnight for 18 hours at 50°C. Slides were washed the next day to remove unbound probes. Ligase was added to circularize the paired ends of bound probes (2 hours at 37°C) followed by enzymatic rolling circle amplification (2 hours at 30°C). Slides were washed in TE buffer before background fluorescence was chemically quenched.

### GENOMIC AND TRANSCRIPTOMIC SEQUENCING

#### Bulk RNA sequencing (Bulk RNA-seq)

##### RNA extraction

Frozen samples were weighed and ice cold TRIzol Reagent (15596026, Thermo Fisher Scientific, Waltham, MA) was added at a ratio of 1 mL per 0.1 mg. Tissue was homogenized using a plastic pestle (Fisher Scientific 50-197-8234) and trituration with a P1000 pipette, followed by 5 min incubation at room temperature. Chloroform (Sigma-Aldrich, C2432, St. Louis, MO) was added to each tube at a ratio of 200 uL per 0.1 mg starting tissue, and incubated 3 min at room temperature or until a separation became clearly visible. Samples were centrifuged at 12000 x g for 10 min at 4°C, after which the top clear aqueous phase was separated into a fresh RNase free tube. Isopropanol was added at a ratio of 500 uL per 0.1 mg starting tissue, and samples were vortexed at 2000 rpm for 1 min, incubated at room temperature for an additional 10 mins, and then centrifuged at 12000 x g for 10 min at 4°C. The supernatant was discarded, and the RNA pellet was washed three times with 1 ml of ice-cold 75% ethanol, centrifuging at 7500 x g for 5 min at 4°C in between each wash. After decanting the third wash, the RNA pellet was air-dried and resuspended in 40μl of RNase-free water. RNA samples were treated with DNaseI, cleaned, and concentrated using the Zymo RNA Clean & Concentrator Kit (Zymo Research R1013) according to manufacturer instructions. A minimum of 7000ng of RNA was obtained from each tissue sample.

##### Library preparation & Sequencing

All RNA samples were coded numerically. Sequencing was performed blind to sample identity by MedGenome. RNA-sequencing libraries were generated using the Illumina TruSeq stranded mRNA kit. The NovaSeq 6000 was used to target an average read depth of 20 million paired (40 million total), 2x100 bp reads per replicate; actual read depth was between 25 and 28 million paired (50 – 56 million total) 2 x 100 bp reads per replicate.

#### Single-nuclei RNA sequencing (snRNA-seq)

##### Capture & Lysis

After FANS, NeuN+ and NeuN- nuclei were recombined in a 1:1 ratio and pelleted by centrifuging at 1000 x g for 10 min at 4°C in a swinging bucket rotor with custom 1.5 mL tube adaptors. Nuclei suspension buffer (NSB) was prepared by diluting 6X NSB in ultrapure H2O and adding 25% bovine serum albumin (BSA). Nuclei were resuspended in a final volume of 20 uL 1X NSB. From this resuspension, 2 uL were used to confirm final nuclei counts using Trypan Blue staining with the Countess 3 Cell Counter. The remaining 18 uL was diluted in 1X NSB to reach a final concentration of 2.5 X 10^3^ nuclei per uL, and 16 uL of this final dilution (40 X 10^3^ total nuclei) were added directly to the thawed PIPs (1 sample per PIP tube), then 2 uL of RNase inhibitor (40 units) were added. Nuclei capture and lysis was performed with the PIPseq V T20 Capture & Barcoding Kit (Fluent Biosciences FB0005358), according to manufacturer instructions for nuclei sample preparation.

##### Library preparation & Sequencing

All RNA samples were coded numerically. Sequencing was performed blind to sample identity by MedGenome. RNA-sequencing libraries were generated using the PIPseq V Library Prep Kit (Fluent Biosciences FB0005372). The NovaSeq X Plus produced was used to target an average read depth of 400 million paired (800 million total), 2x100 bp reads per replicate; actual read depth was between 380 and 470 million paired (760 - 940 million total), 2x100 reads per replicate.

##### Quality control

Quality control analyses were performed after FANS (cell viability and counting) and after library preparation (RNA concentration and size) (**Fig. S3a**). Four samples (WT1, WT2, WT3, WT4, KO1, KO2, KO3, KO4) were processed through library preparation. However one WT sample (WT1) failed QC after library preparation due to low RNA concentration. To maintain comparable sample and cell numbers, both WT1 and KO1 were excluded from sequencing and downstream analyses.

### CUT&RUN

#### Isolation of target chromatin

After FANS, NeuN- nuclei (WT vs KO glial CUT&RUN) or GFP+ nuclei (WT astrocyte CUT&RUN) were pelleted by centrifuging at 1000 x g for 10 min at 4°C in a swinging bucket rotor with custom 1.5 mL tube adaptors. Target chromatin was isolated using the CUTANA™ ChIC/CUT&RUN Kit Version 5 (EpiCypher 14-1048) or CUTANA™ meCUT&RUN Kit for DNA Methylation Sequencing (Epicypher 14-1060-24), according to manufacturer instructions. Primary antibody dilutions were scaled to maintain a consistent antibody mass (0.5 μg) per reaction: Zbtb20 (1:5), H3K4me3 (1:50), IgG (1:50), and anti-GST (1:50) were selected based on manufacturer recommendations. GST-MeCP2 was diluted 1:20. *E. coli* spike-in DNA dilutions were scaled according to expected yield of each reaction. For WT vs KO glia CUT&RUN: 0.01 ng per reaction for all targets (Zbtb20, IgG, GST-MeCP2, anti-GST). For WT astrocyte CUT&RUN: 0.05 ng per reaction for low-abundance targets (Zbtb20 and IgG) and 0.1 ng per reaction for higher-abundance targets (H3K4me3).

#### Library preparation & Sequencing

All DNA samples were coded numerically. Sequencing was performed blind to sample identity by MedGenome. DNA-sequencing libraries were generated using the CUTANA™ CUT&RUN Library Prep Kit (EpiCypher 14-1001, 14-1002). For WT vs KO glia CUT&RUN, the NovaSeq X Plus was used to target an average read depth of 12 million paired (24 million total) 2x150 bp reads per replicate; actual read depth was between 6.5 and 16 million paired (13 to 32 million total), 2x150 reads per replicate. For WT astrocyte CUT&RUN, the the NovaSeq X Plus was used to target an average read depth of 8 million paired (16 million total) 2x50 bp reads per replicate; actual read depth was between 5.5 and 9.5 million paired (11 to 19 million total), 2x50 reads per replicate.

### BIOINFORMATICS AND DATA ANALYSIS

#### Imaging data analysis

##### For neural cell counting

The counting of DAPI, Sox9, Olig2, and NeuN-labeled cells was performed as previously described [72]. Tile scan images (11 z-stacks with 1µm step-size; 10µm total) containing the primary visual cortex (V1) from P21 WT and *Ctnnd2*-KO mice were acquired rapidly at 30X objective using the resonant scanner of an Olympus FV 3000. The resolution of the stitched images was digitally restored to improve resolution using a trained content-aware image restoration (CARE) network [125]. The ROI of stitched images used for quantifying cell numbers spanned L1 through L6 of V1 (912µm x 1326µm). A machine-learning-based method [126] was used to segment and identify cells labeled by each marker. The complete source code for this method is available at https://github.com/ErogluLab/CellCounts. 3 images per animal were used for segmentation and analysis. Segmented images were reviewed and adjusted, when necessary, for accuracy. The layer-specific density of each cell type was calculated by dividing count within bin by bin area. The density of each cell type was also calculated within the entire image.

##### For neuronal subtype counting

Sections were imaged using a 20× objective to capture tilescans of the primary visual cortex spanning pia to corpus callosum with high resolution (1 μm step size, 5 μm z-stack) on a Leica Stellaris 8 confocal. 32 images were taken from the entire dataset (4 images per animal, 4 animals per genotype for WT and KO). Images were then cropped to rectangles of 200 pixel width and variable height, excluding the corpus callosum. Image analysis was performed in QuPath (v0.6.0) and cell detection of Cux1+, RORβ+, or Ctip2+ cells was performed using QuPath’s built-in “Positive Cell Detection.” The data for each image and cell were exported.

Proportions of each neuronal subtype were calculated and Chi-square tests performed in Graphpad Prism. The spatial frequency distribution of each neuronal subtype was calculated by dividing subtype count within bin by total subtype count in image. The density of each neuronal subtype was calculated within an ROI containing 75% (Layer 2/3) or 90% (Layer 4 and 5/6) of the subtype of interest. The complete source code for this analysis is available upon request.

##### For OPC and MOL counting

sections were imaged using a 20× objective to capture tilescans of the primary visual cortex spanning pia to corpus callosum with high resolution (1 μm step size, 7 μm z-stack) on a Leica Stellaris 8 confocal. 18 images were taken from the entire dataset (3 images per animal, 3 animals per genotype for WT and KO). Images were then cropped to rectangles of 200 pixel width and variable height, excluding the corpus callosum. Image analysis was performed in QuPath (v0.6.0) and cell detection of ASPA+ or Pdgfra+ cells was performed using QuPath’s built-in “Positive Cell Detection.” The data for each image and cell were exported.

Proportions of each oligodendroglia subtype were calculated and Chi-square tests performed in Graphpad Prism. The spatial frequency distribution of each oligodendroglia subtype was calculated by dividing subtype count within bin by total subtype count in image. The density of each oligodendroglia subtype was calculated within the entire image. The complete source code for this analysis is available upon request.

##### For NFOL and MFOL counting

sections were imaged using a 20× objective to capture tilescans of the primary visual cortex spanning pia to corpus callosum with high resolution (1 μm step size, 7 μm z-stack) on a Leica Stellaris 8 confocal. 35 images were taken for the entire data set (2-3 images per animal, 6 animals per genotype for WT and KO). The images were cropped into rectangles of 200 pixel width and variable height to include only the cortex (excluding the corpus callosum) in ImageJ. The images were then processed using ImageJ’s Cell Counter to manually quantify Bcas1+ and MBP+/Bcas1+ cells. The data for each section and cell were exported.

Proportions of each oligodendroglia subtype were calculated and Chi-square tests performed in Graphpad Prism. The spatial frequency distribution of each oligodendroglia subtype was calculated by dividing subtype count within bin by total subtype count in image. The complete source code for this analysis is available upon request.

##### For smFISH *Arc* detection

Slides were imaged using a 20X objective on the Leica Stellaris 8 system. 2-5 ROIs were imaged per animal of binocular primary visual cortex contralateral to the eye closure. Images were then analyzed blinded to condition using FIJI to measure *Arc* signal length in binocular zone of primary visual cortex. Unblinded data was analyzed using Graphpad Prism. The complete source code for this analysis is available upon request.

##### For multiplexed IF/smFISH

Sections were imaged within 2 days using a 20× objective to capture tilescans of the primary visual cortex spanning pia to corpus callosum with high resolution (1 μm step size, 3 μm z-stack) on an Leica Stellaris 8 microscope. In total, 21 images were taken for the entire dataset (2-3 sections per animal per probeset, three animals per genotype for WT and KO). The images were then processed using a Fiji custom pipeline that includes thresholding and masking of S100-beta signal and quantification of *Mfge8, Tnc,* and *Id4* intensity with S100-beta-positive astrocyte area. The normalized signal was calculated by dividing mean signal within bin by mean signal within entire image. The complete source code for this analysis is available upon request.

#### Bulk RNA-seq data processing and analysis

Trimmomatic (v0.39) was used to trim the raw reads of adapters, and STAR (v2.7.10a) [127] was used to align the reads to the reference mouse genome (mm10). Subread (featureCounts v2.0.3) was used to count the reads. The raw data was uploaded to the NCBI Gene Expression Omnibus and is available upon request. Counts were then imported into R (v4.3.2). DESeq2 (v1.42.1) was used to conduct differential gene expression analysis in R. ClusterProfiler (v4.10.1) was used to identify GO terms for differentially expressed genes [128]. The complete source code for this analysis is available upon request.

#### snRNA-seq data processing and analysis

Raw reads were processed following the PIPseeker 3.3.0 pipeline for v4.0PLUS chemistry. Cell calling sensitivity 4 was used and counts matrices were then imported into Jupyter Notebook (Python v3.12.7) using Scanpy (v1.11.5). For each dataset, quality control and filtering were performed as described in Best Practices for Single-Cell Analysis Across Modalities [129]. In brief, cells were filtered for abnormally high number of genes by counts, total counts, and percent mitochondrial counts to eliminate low quality cells. Counts were normalized and highly variable genes were identified using Scanpy. Counts were then scaled and clusters were identified following PCA, UMAP, and neighbor calculations.

Major neural lineages (astrocytes, oligodendrocyte lineage, interneurons, and excitatory neurons) were identified by performing unbiased Leiden clustering using Scanpy. To annotate clusters, Wilcoxon rank-sum tests were performed comparing each cluster to every other cluster, and top cluster-defining genes were cross-referenced with literature-based gene lists. Wilcoxon Rank-sum tests were next performed across genotypes for each of the 4 lineages, and significant DEGs were used for GO analysis in R using clusterProfiler.

Prior to subclustering the major neural lineages, batch effects were corrected using scVI (v1.3.3) [82]. Next, nuclei belonging to each major lineage previously identified were subsetted, and new anndata objects were created for each. Unbiased Leiden clustering was repeated for each new anndata object, and clusters were identified by cross-referencing literature-based gene lists.

Raw counts, raw count matrices, and filtered count matrices were uploaded to NCBI Gene Expression Omnibus and is available upon request. The complete source code for this analysis is available upon request.

#### CUT&RUN data processing and analysis

The Cut-and-RUN Analysis Suite (CaRAS) was used to perform QC, cleanup, alignment, and peak-calling. Briefly, FastQC (v0.12.1) was used to perform an initial quality assessment of raw unaligned reads, clumpify (bbtools v39.01) was used for deduplification of reads, BBDuk (bbtools v39.01) for adapter trimming, and Trimmomatic for quality trimming [130]. Repeat quality assessment of pre-processed reads was performed using FastQC (v0.12.1). Reads were aligned to the target organism genome (mm10) and normalization organism genome (eColiK12) using the mem algorithm (BWA v0.7.18-r1243-dirty). Samtools (v1.18) was used to perform quality filtering, sorting, and indexing of aligned reads. bamCoverage (deeptools v3.5.5) was used to generate aligned read bedgraph files and visualization tracks (bigwig files). For each sample, a custom script (samtools v1.18) was used to normalize aligned reads based on the number of eColiK12 reads. Repeat quality assessment of aligned reads was performed using FastQC (v0.12.1). Peak-calling was performed using MACS2 (v2.2.9.1), GEM (v2.7-0), HOMER, Genrich (v0.6.1), SEACR (v1.3), and SICER2 [131]. Peaks identified by the six peak callers were merged using mergePeaks (HOMER). A custom script (samtools v1.18) was used to calculate weighted peak center fold enrichment, corrected based on ChIP-to-control normalization factor. A custom script (IDR v2.0.4.2) was used to perform irreproducibility rate calculation; final IDR value shows the chance of a peak being unable to be reproduced by different peak calling algorithms. Peaks identified by at least 5 peak callers and annotated as promoters were used for downstream analysis.

Chipseeker (v1.38.0) was used to annotate peaks, with promoters defined as within 1.5kb of transcription start sites (TSSs) [132, 133]. computeMatrix, plotHeatmap, and plotProfile (deeptools v3.5.6) were used to visualize Zbtb20 and MBD coverage surrounding transcription start sites of promoter peaks. HOMER was used to perform motif enrichment analysis separately on *Ctnnd2*-WT and *Ctnnd2*-KO Zbtb20-bound promoter sets. CpG island track was downloaded in BED format from the UCSC table browser and cross-referenced with union set of *Ctnnd2*-WT and *Ctnnd2*-KO Zbtb20-bound promoters. To infer RNA expression changes at genes with Zbtb20-bound promoters, P28 snRNAseq was subsetted to isolate either *Rbfox3*-negative nuclei (glia CUT&RUN) or astrocyte nuclei (astrocyte CUT&RUN). Differential expression analysis was performed on each subset using Scanpy (v1.11.5) to perform Wilcoxon Rank-sum tests across genotypes.

The raw data was uploaded to the NCBI Gene Expression Omnibus and is available upon request. The complete source code for this analysis is available upon request.

#### Xenium data processing and analysis

Imaging processing, decoding, quality score of gene transcripts, and DAPI-based cell segmentation was automatically performed by the Xenium Analyzer system. Data was acquired on instrument software version 4.0.1.4 and analysis version xenium-4.0.1.0.

After pre-processing, spatialdata (v0.5.0) and spatialdata_io (v0.3.0) were used to read .zarr stores into spatialdata objects, which were subsequently converted to anndata objects containing count matrices, cell, and gene annotations. Quality control and filtering was performed using Scanpy (v1.11.5) on a merged anndata object containing all detected cells from all 4 biological replicates. In brief, cells were filtered for abnormally low number of genes by counts. Counts were normalized using scanpy pp.normalize_total function.

Separate QC-filtered anndata objects were generated for each sample. Each anndata object was further filtered for cells contained within a hand-drawn visual cortex ROI selected in the Xenium Explorer. Anndata objects from each animal were combined into a single anndata object, scVI (v1.3.3) was used to perform batch correction, and Tacco (v0.4.0.post1) was used to perform label transfer from the P28 visual cortex snRNAseq dataset.

The raw and processed data were uploaded to the NCBI Gene Expression Omnibus and is available upon request. The complete source code for this analysis is available upon request.

#### Quantification and statistical analysis

Statistical analyses were performed using Graphpad Prism (v11.0.0) and R Statistical Software (v4.3.2). Details on sample size and the tests used can be found in the figure legends or methods subsection for specific experiments. In all cases, “N” refers to biological replicates and “n” represents the number of images or cells analyzed within biological replicates.

## KEY RESOURCES TABLE

| REAGENT or RESOURCE | SOURCE | IDENTIFIER |
| --- | --- | --- |
| Antibodies |  |  |
| Mouse Monoclonal anti-NeuN Antibody, clone A60 | Millipore | Cat# MAB377,<br>RRID:AB_2298772 |
| Rabbit polyclonal anti-Sox9 | Millipore | Cat# AB5535,<br>RRID:AB_2239761 |
| Mouse IgG2a monoclonal anti-Olig2, clone 211F1.1 | Millipore | Cat# MABN50<br>RRID: AB_10807410 |
| Rabbit Anti-S100 beta Monoclonal, Unconjugated, Clone EP1576Y | Abcam | Cat# ab52642,<br>RRID:AB_882426 |
| Mouse anti-S100 beta Monoclonal, Clone SH-B1 | Sigma-Aldrich | Cat# S2532;<br>RRID:AB_477499 |
| Rabbit polyclonal anti-ZBTB20 | Sigma-Aldrich | Cat# HPA016815;<br>RRID:AB_1858947 |
| Rat IgG2a monoclonal anti-Ctip2 [25B6] | Absolute Antibody | Cat# Ab00616-7.4 |
| Mouse IgG2b monoclonal anti-ROR-beta [N7927] | R and D Systems | Cat# PP-N7927-00,<br>RRID:AB_3659610 |
| Rabbit polyclonal anti-CUX1 | Proteintech | Cat# 11733-1-AP,<br>RRID:AB_2086995 |
| Anti-NeuN, clone A60, Alexa Fluor 488 conjugated | Millipore | Cat# MAB377X,<br>RRID:AB_2149209 |
| Mouse IgG2b monoclonal anti-MBP | Thermo Fisher Scientific | Cat# MA1-10837<br>RRID:AB_1077025 |
| Rabbit polyclonal anti-ASPA | GeneTex | Cat# GTX113389<br>RRID: AB_2036283 |
| Mouse IgG1 monoclonal anti-Bcas1 | Santa Cruz | Cat# SC-136342<br>RRID AB_10839529 |
| Rat monoclonal anti-Pdgfrα | BD Biosciences | Cat# 558774<br>RRID: AB_397117 |
| Rabbit monoclonal Anti-Delta-catenin antibody | Abcam | Cat# ab184917,<br>RRID:AB_3661881 |
| Mouse monoclonal Anti-Delta-catenin antibody | Santa Cruz Biotechnology | Cat# sc-81793,<br>RRID:AB_1120808 |
| Mouse monoclonal Anti-Delta-catenin antibody | BD Biosciences | Cat# 611537,<br>RRID:AB_398995 |
| Mouse monoclonal anti-Gapdh antibody | Abcam | Cat# ab8245,<br>RRID:AB_2107448 |
| Rabbit anti-Gapdh antibody | Abcam | Cat# ab9485,<br>RRID:AB_307275 |
| H3K4me3 Antibody, SNAP-Certified™ for CUT&RUN | EpiCypher | Cat# 13-0060 |
| CUTANA™ Rabbit IgG CUT&RUN Negative Control Antibody | EpiCypher | Cat# 13-0042,<br>RRID:AB_2923178 |
| CUTANA™ Anti-GST Tag Antibody | EpiCypher | Cat# 13-0073 |
| IRDye 800CW Goat anti-Mouse IgG Secondary Antibody | LICORbio | Cat# 926-32210,<br>RRID:AB_621842 |
| IRDye 680RD Goat anti-Mouse IgG Secondary Antibody | LICORbio | Cat# 926-68070,<br>RRID:AB_10956588 |
| IRDye 680RD Goat anti-Rabbit IgG Secondary Antibody | LICORbio | Cat# 926-68071,<br>RRID:AB_10956166 |
| IRDye 680RD Goat anti-Mouse IgG Secondary Antibody | LICORbio | Cat# 925-68070,<br>RRID:AB_2651128 |
| Alexa Fluor 488 conjugated Goat anti-rabbit | Thermo Fisher Scientific | Cat# A-11008,<br>RRID:AB_143165 |
| Alexa Fluor 488 conjugated Goat anti-mouse IgG1 | Thermo Fisher Scientific | Cat# A-21121,<br>RRID:AB_2535764) |
| Alexa Fluor 488 conjugated Goat anti-Mouse IgG2b | Thermo Fisher Scientific | Cat# A-21141<br>RRID: AB 2535778 |
| Alexa Fluor 488 conjugated Goat anti-Rat | Thermo Fisher Scientific | Cat# A-11006,<br>RRID:AB_2534074 |
| Alexa Fluor 594 conjugated Goat anti-Chicken IgY | Thermo Fisher Scientific | Cat# A-11042<br>RRID: AB_2534099 |
| Alexa Fluor 594 conjugated Goat anti-Mouse IgG1 | Thermo Fisher Scientific | Cat# A-21125<br>RRID: AB 2535767 |
| Alexa Fluor 594 conjugated Goat anti-Rat | Thermo Fisher Scientific | Cat# A-11007<br>RRID: AB_10561522 |
| Alexa Fluor 594 conjugated Goat anti-rabbit | Thermo Fisher Scientific | Cat# A-11012,<br>RRID:AB_2534079 |
| Alexa Fluor 594 conjugated Goat anti-Mouse IgG2b | Thermo Fisher Scientific | Cat# A-21145,<br>RRID:AB_2535781 |
| Alexa Fluor 647 conjugated Goat anti-Mouse IgG1 | Thermo Fisher Scientific | Cat# A-21240,<br>RRID:AB_2535809 |
| Alexa Fluor 647 conjugated Goat anti-Mouse IgG2a | Thermo Fisher Scientific | Cat# A-21241<br>RRID: AB 2535810 |
| Alexa Fluor 647 conjugated Goat anti-rat | Thermo Fisher Scientific | Cat# A-21247,<br>RRID:AB_141778 |
| Alexa Fluor 647 conjugated Goat anti-rabbit | Thermo Fisher Scientific | Cat# A-21245,<br>RRID:AB_2535813 |
| Bacterial and virus strains |  |  |
| One Shot™ Stbl3™ Chemically Competent *E. coli* | Thermo Fisher Scientific | Cat# C737303 |
| Chemicals, peptides, and recombinant proteins |  |  |
| Tissue-Tek® O.C.T. Compound | Sakura® Finetek | Cat# 4583 |
| Triton™ X-100 Surfact-Amps™ Detergent Solution | Thermo | Cat# 28314 |
| VWR® Superfrost® Plus Micro Slide | VWR | Cat# 48311-703 |
| DAPI | Thermo Fisher Scientific | Cat# D1306 |
| Tween® 20 (Polysorbate), Molecular Biology Grade | Promega | Cat# H5151 |
| Formaldehyde (37% by Weight/Molecular Biology) | Thermo Fisher Scientific | Cat# BP531-25 |
| Nuclei Extraction Buffer | Miltenyi Biotec | Cat # 130-128-024 |
| DPBS, no calcium, no magnesium | Thermo-Fisher | Cat# 14190144 |
| 7-AAD (7-Aminoactinomycin D) | Thermo Fisher Scientific | Cat# A1310 |
| Bovine Serum Albumin (BSA), lyophilized powder | Sigma-Aldrich | Cat# A4161 |
| cOmplete™ Protease Inhibitor Cocktail tablets | Roche | Cat# 04693132001 |
| SUPERase• In™ RNase Inhibitor (20 U/μL) | Thermo Fisher Scientific | Cat# AM2696 |
| L.A.B. Solution (Liberate Antibody Binding Solution) | Polysciences | Cat# 24310 |
| TRIzol Reagent | Thermo Fisher Scientific | Cat# 15596026 |
| GST-MeCP2 for meCUT&RUN | EpiCypher | Cat# 15-2002 |
| Critical commercial assays |  |  |
| HCR Gold RNA-FISH Kit | Molecular Instruments | N/A |
| HCR RNA-FISH Kit v3.0 | Molecular Instruments | N/A |
| PIPseq V T20 Capture & Barcoding Kit | Fluent Biosciences | Cat# FB000538 |
| PIPseq V Library Prep Kit | Fluent Biosciences | Cat# FB0005372 |
| CUTANA™ ChIC/CUT&RUN Kit Version 5 | EpiCypher | Cat# 14-1048 |
| CUTANA™ meCUT&RUN Kit for DNA Methylation Sequencing | EpiCypher | Cat# 14-1060-24 |
| CUTANA™ CUT&RUN Library Prep Kit | EpiCypher | Cat# 14-1001, 14-1002 |
| Xenium Slides & Sample Prep Reagents | 10X Genomics | Cat# 1000460 |
| Xenium Mouse Brain Gene Expression Panel | 10X Genomics | Cat# 1000462 |
| Xenium Add-on Custom 51 to 100 Gene Panel (4 rxn) | 10X Genomics | Cat# 1000651 |
| Anti-Nucleus MicroBeads | Miltenyi Biotec | Cat# 130-132-997 |
| Zymo RNA Clean & Concentrator Kit | Zymo Research | Cat# 1013 |
| 4–15% Mini-PROTEAN® TGX Stain-Free™ Protein Gel | Bio-Rad | Cat #4568084 |
| Deposited data |  |  |
| Bulk RNA sequencing of WT and *Ctnnd2*-KO visual and prefrontal cortex | This paper | Upon Request |
| Single-nuclei RNA sequencing of WT and *Ctnnd2*-KO visual cortex | This paper | Upon Request |
| Xenium analysis of WT and *Ctnnd2*-KO posterior neocortex | This paper | Upon Request |
| CUT&RUN sequencing of WT and *Ctnnd2*-KO glial nuclei | This paper | Upon Request |
| CUT&RUN sequencing of astrocyte nuclei | This paper | Upon Request |
| Experimental models: Organisms/strains |  |  |
| Mouse: *Ctnnd2* KO | This paper | N/A |
| Mouse: CAG-Sun1/sfGFP | JAX | RRID:IMSR_JAX:021039 |
| Mouse: C57BL6/J | JAX | RRID:IMSR_JAX:000664 |
| Oligonucleotides |  |  |
| sgRNA sequences for targeted *Ctnnd2* exon 4/5 deletion, see Table 1 | This paper | N/A |
| Primer sequences for allelic subcloning, see Table 2 | This paper | N/A |
| Primer sequences for *Ctnnd2*-KO genotyping, see Table 3 | This paper | N/A |
| Recombinant DNA |  |  |
| GEARBOCS | Addgene | 218181 |
| pAD-ΔF6 | Addgene | 112867 |
| pUC-mini-iCAP-PHP.B | Addgene | 103005 |
| Software and algorithms |  |  |
| ImageJ (v1.54j) | NIH | https://imagej.net/software/fiji/ |
| QuPath (v0.6.0) | Queen's University,<br>Belfast, Northern<br>Ireland | https://github.com/qupath/qupath/releases/tag/v0.6.0 |
| Xenium Analyzer Software (4.0.1.4 and xenium-4.0.1.0) | 10X Genomics | https://www.10xgenomics.com/support/software/xenium-explorer/downloads |
| R Statistical Software (v4.3.2) | RStudio | https://www.r-project.org/ |
| Graphpad Prism (v11.0.0) | GraphPad Software<br>Inc., La Jolla, CA,<br>USA | www.graphpad.com |
| Python (v3.10.18 and v3.13.7) | Python programming<br>language | https://www.python.org/downloads/ |
| Other |  |  |
| gentleMACS Octo Dissociator | Miltenyi Biotec | Cat# 130-095-937 |
| gentleMACS Octo Coolers | Miltenyi Biotec | Cat# 130-130-533 |
| gentleMACS C Tubes | Miltenyi Biotec | Cat# 130-096-334 |
| MACS SmartStrainers (70 µm) | Miltenyi Biotec | Cat# 130-110-916 |
| MACS SmartStrainers (30 µm) | Miltenyi Biotec | Cat# 130-098-458 |
| PolyPestle Double-Sided Pestle | Thermo Fisher<br>Scientific | Cat# 50-197-8234 |
| DNA LoBind® Tubes, 1.5 mL | Eppendorf | Cat# 022431021 |
| MA900 Multi-Application Cell Sorter | Sony |  |
| Mfge8 HCR HiFi probe | Molecular Instruments | N/A |
| Tnc HCR HiFi probe | Molecular Instruments | N/A |
| Id4 HCR HiFi probe | Molecular Instruments | N/A |
| Arc HCR probe | Molecular Instruments | N/A |
| HybEZ hybridization system | ACDBio |  |
| 10X Genomics Xenium Analyzer | 10X Genomics | RRID:SCR_023910 |
| Power Blotter–Semi-dry Transfer System | Invitrogen |  |
| Odyssey Infrared Imaging System | LICORbio |  |

## AUTHOR CONTRIBUTIONS

Conceptualization, G.S., C.T., C.E., K.S., R.B.; methodology, G.S., C.T., E.H., J.S., K.S., J.R.; Investigation, G.S., C.T., J.S., G.R., J.D., G.M, K.S.; writing—original draft, G.S. and C.E.; writing—review/editing, G.S., C.T., J.S., G.M., G.R., J.D., E.H., J.R., R.B., K.S., and C.E.; funding acquisition, G.S. and C.E.

## DECLARATION OF GENERATIVE AI AND AI-ASSISTED TECHNOLOGIES

During the preparation of this work, the authors used ChatGPT (OpenAI) and Claude (Anthropic) in order to troubleshoot code. After using this tool or service, the authors reviewed and edited the content as needed and take(s) full responsibility for the content of the publication.

## SUPPLEMENTARY INFORMATION

### Supplementary Figures

**Figure S1:**
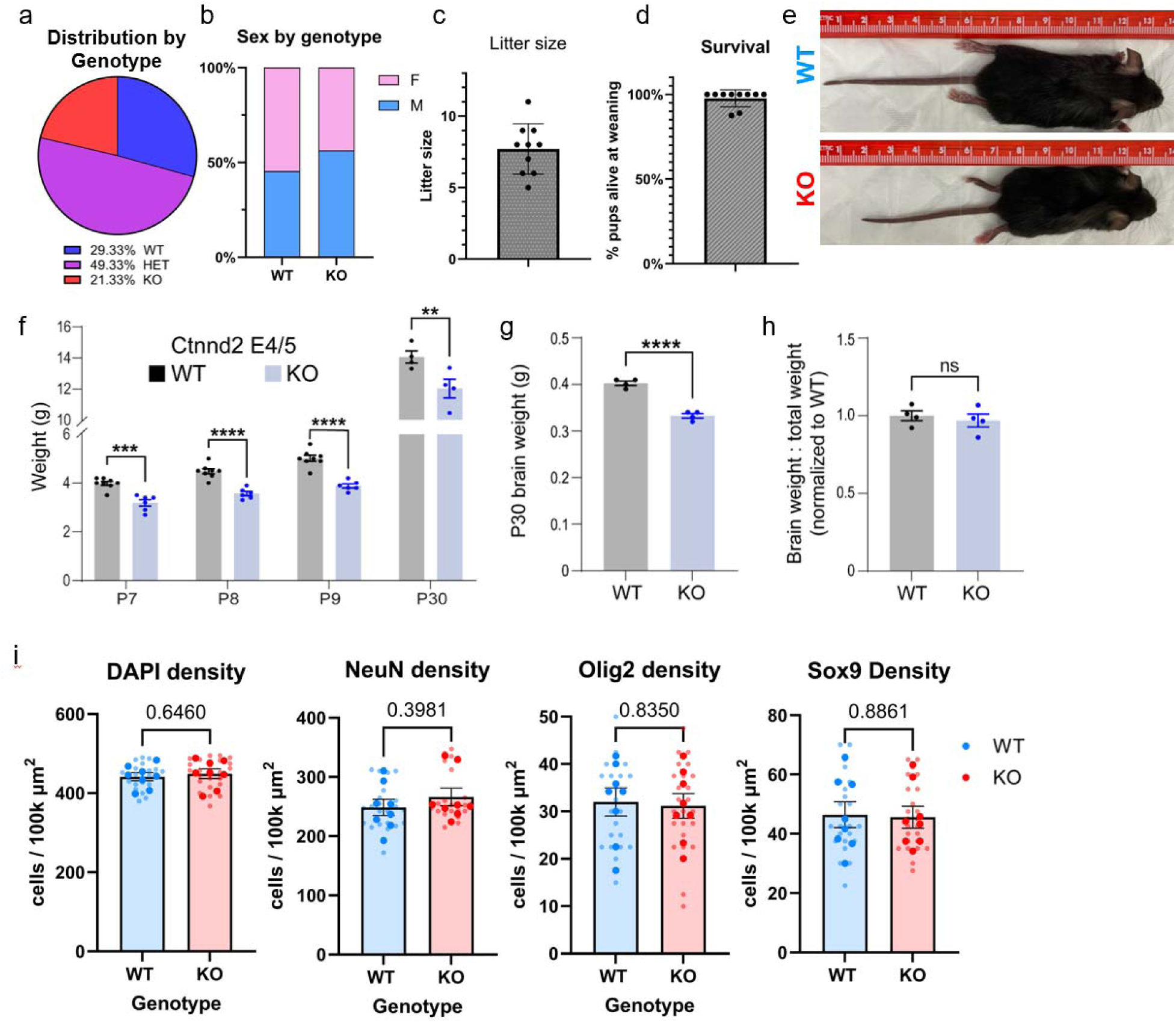
Characterization of Ctnnd2-KO mouse model (related to Figure 1). A-D) 10 independent litters from *Ctnnd2*-Het breeding pairs: (A) Genotype distribution, Chi-square test of independence, (2) = 0.9733, p = ns. B) Sex distribution, Chi-square test of independence, (1) = 0.4318, p = ns. C) Litter size and D) survival rate for offspring of *Ctnnd2*-Het breeding pairs. E) Photograph illustrating size difference between P30 Ctnnd2 E4/5 WT and KO sex-matched (male) littermates. F) Analysis of Ctnnd2 WT and KO body weight. N=4-8 littermate pairs per age. Data points represent mouse averages. Bars are mean ± s.e.m. Multiple unpaired two-tailed t-tests with Welch’s correction. P7 (p=6.03e-4), P8 (p=1.8e-6), P9 (p=6e-6) and P30 (p=3.5e-2). N=4-8 animals per genotype per timepoint. G-H) Analysis of P30 *Ctnnd2* WT and KO brain weight (G) and brain weight normalized to body weight (H). N=4 sex-matched littermates, Welch’s t-test. Brain weight (p=6e-4), normalized brain weight (ns). I) Unbinned densit analysis of nuclei (DAPI), neurons (NeuN+), astrocytes (Sox9+), and oligodendrocytes (Olig2+) in the P21 visual cortex. N=8 mice of either sex per genotype, n=3 technical replicates/mouse. Welch’s unpaired t-test.

**Figure S2:**
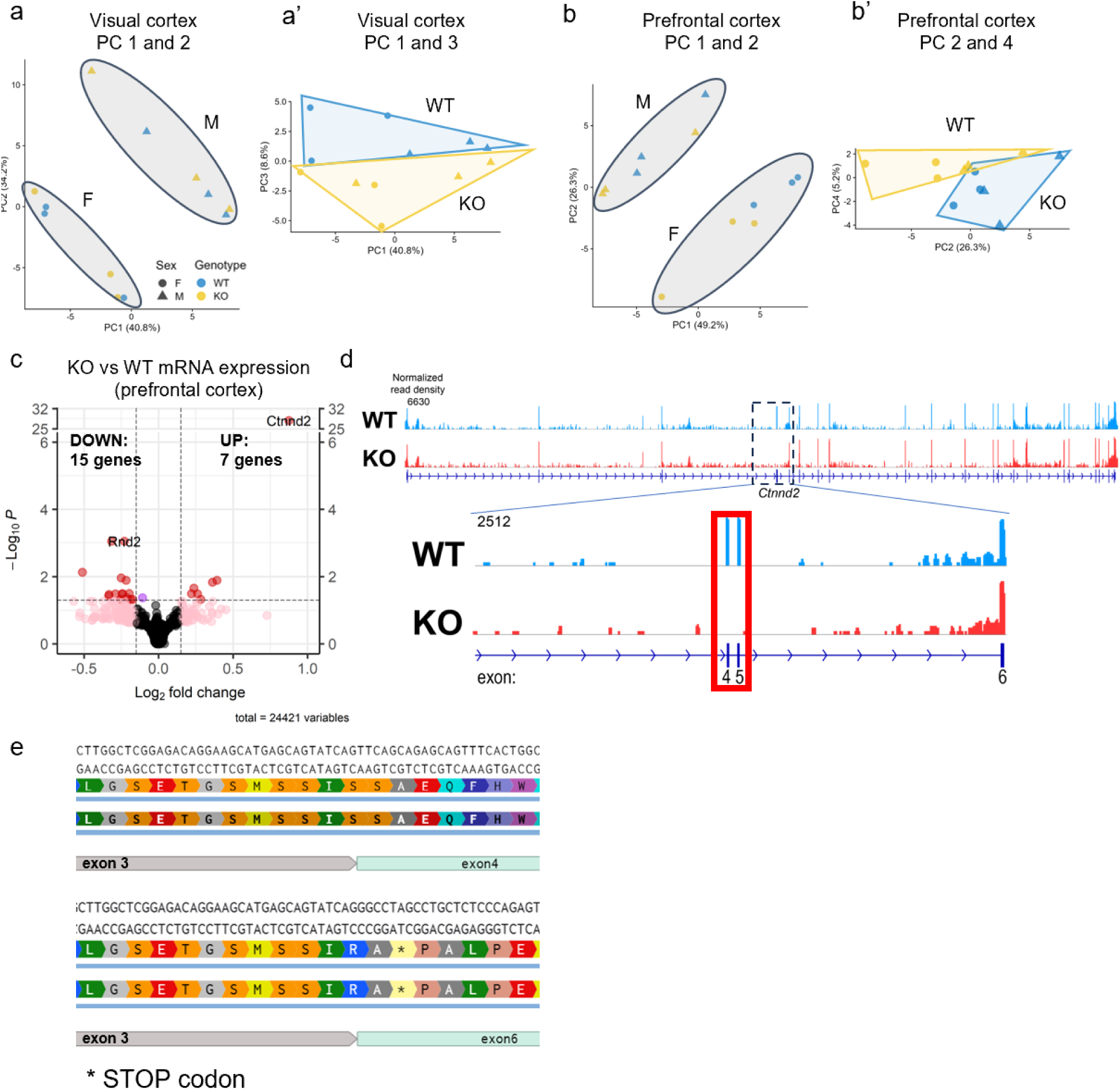
Additional analysis and quality control of RNA sequencing (related to Figure 1). A) PCA plots showing segregation of visual cortex samples by sex (A) and genotype (A’). B) PCA plots showing segregation of prefrontal cortex samples by sex (B) and genotype (B’). C) Volcano plot of DEGs in the prefrontal cortex of KO (N=6, 3 males, 3 females) compared to WT (N=6, 3 males, 3 females). Criteria for DEG selection: log2 fold change (FC) less than -0.15 or greater than 0.15; false discovery rate (FDR) < 0.05 as calculated by DESeq2 with Benjamini–Hochberg’s correction. D) RNA sequencing reads aligned to *Ctnnd2* genomic DNA sequence. Reads aligned to exons 4 and 5 are absent in KO samples (red rectangle). E) Protein translation of KO allele showing early stop codon with excision of C*tnnd2* exons 4 and 5.

**Figure S3:**
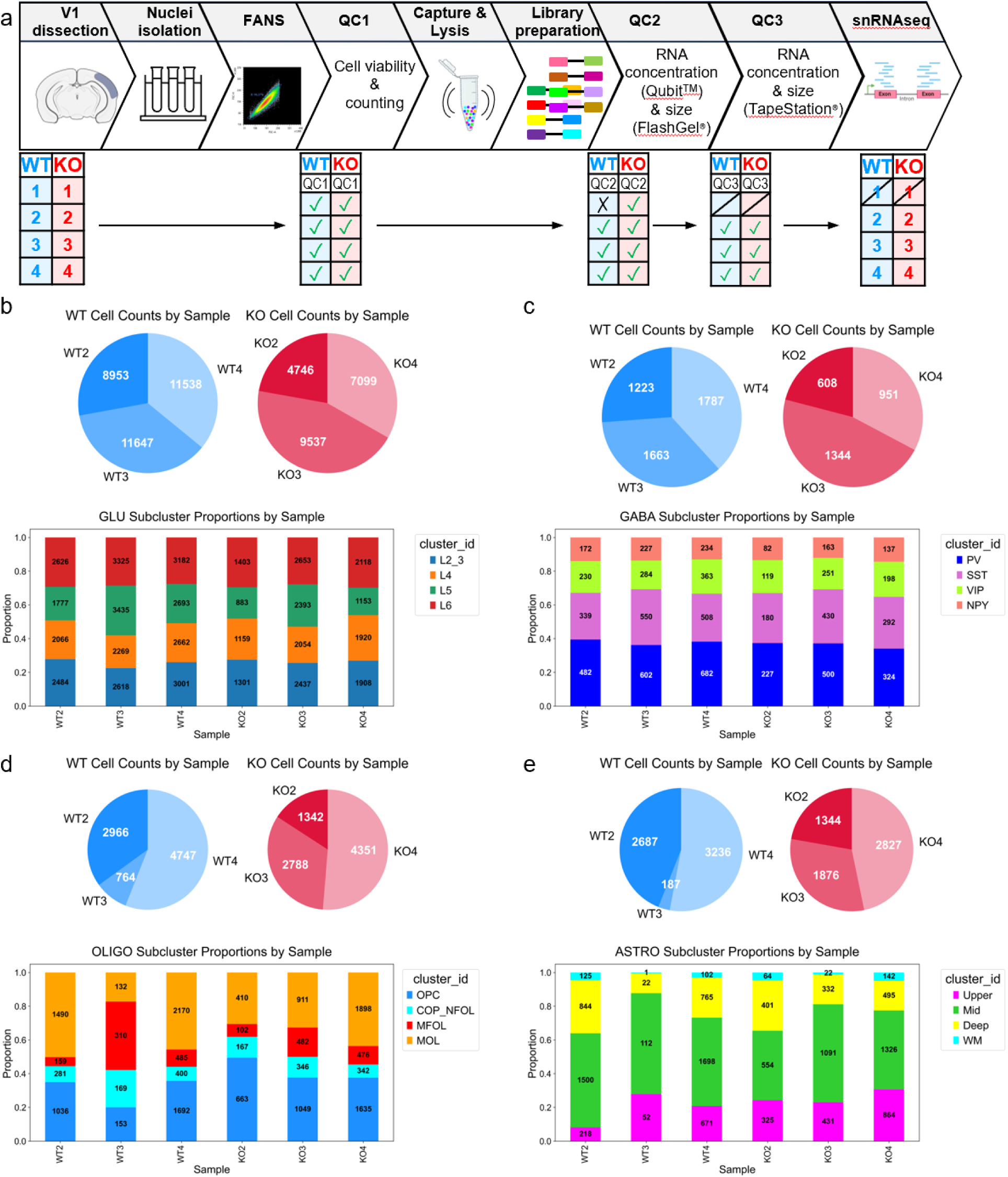
Additional analysis and quality control of snRNA sequencing (related to Figure 3). A) Detailed flow chart of snRNAseq experimental design, including sample filtering at successive qualit control (QC) steps. B-E) Pie charts of sample distribution (top) and contingency plots (bottom) of cellular subtype distribution in each individual sample for glutamatergic neurons (B), GABAergic neurons (C), oligodendrocytes (D), and astrocytes (E).

**Figure S4:**
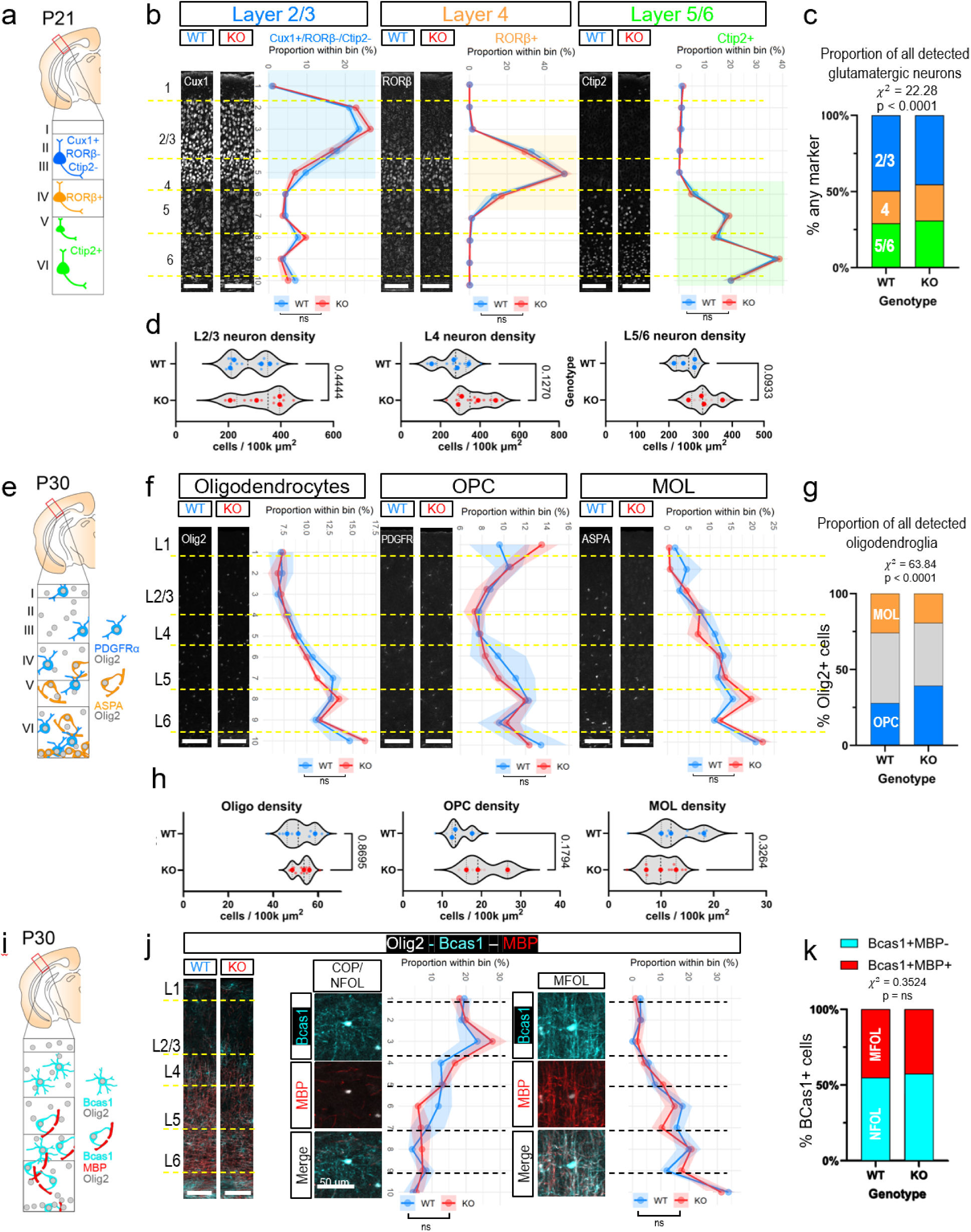
The effects of 6-catenin loss on glutamatergic neurons and oligodendroglia in the visual cortex (related to Figure 3). A) P21 visual cortex was stained for layer-specific markers (Cux1, RORβ, Ctip2) to identify glutamatergic neuron molecular subtypes and analyzed with respect to relative distance from the pia to corpus callosum. B) Representative images of Cux1 (left), RORβ (middle), and Ctip2 (right) staining and histograms of Layer 2/3 (Cux1+/RORβ-/Ctip2-), Layer 4 (RORβ+/Ctip2-), and Layer 5/6 (RORβ-/Ctip2+) molecular subtype distributions across V1 in WT and *Ctnnd2*-KO. No significant differences between genotypes (two-way ANOVA). Scale bars: 100 μm. All values represent the mean[±[SEM. C) Barplot illustrating the proportion of L2/3, L4, and L5/6 neuron nuclei found in each genotype. Chi-square test of independence, **x**^2^ (3) = 22.28, p < 0.0001. D) Density analysis of L2/3, L4, and L5/6 neuron nuclei within ROIs corresponding to L2/3, L4, and L5/6, respectively. Welch’s unpaired t-test. For B-D: N=4 animals per genotype from 3 litters, n=2-3 technical replicates/mouse. E) P30 visual cortex was stained for stage-specific markers of oligodendrocyte lineage cells (Olig2, PDGFRα, ASPA) and analyzed with respect to relative distance from the pia to corpus callosum. OPC, Oligodendrocyte precursor cells; MOL, mature oligodendrocytes; NFOL, newly formed oligodendrocytes; MFOL, myelinating newly formed oligodendrocytes. F) Representative images of Olig2 (left), PDGFRα (middle), and ASPA (right) staining and histograms of oligodendrocytes (left), OPCs (middle), and MOLs (right) distribution across V1 in WT and *Ctnnd2*-KO. No significant differences between genotypes (two-way ANOVA). Scale bars: 100 μm. All values represent the mean[±[SEM. G) Barplot illustrating the proportion of OPCs (PDGFRα+/ASPA-) and MOL (PDGFRα-/ASPA+) found in each genotype. Chi-square test of independence, **x**^2^ (2) = 63.84, p < 0.0001. H) Unbinned density analysis of OPCs and MOLs in the P30 visual cortex. Welch’s unpaired t-test. For F-H: N=3 animals per genotype from 2 litters, n=3 technical replicates/mouse. I) P30 visual cortex was stained for myelination status of newly formed oligodendrocytes (Bcas1, MBP) and analyzed with respect to relative distance from the pia to corpus callosum. J) Representative images of Bcas1 and MBP staining across V1 cortex (scale bars: 100 μm) with high magnification insets (scale bars: 50 μm) and histograms of NFOL (left), MFOL (right) distribution across V1 in WT and *Ctnnd2*-KO. No significant differences between genotypes (two-way ANOVA). All values represent the mean[±[SEM. K) Barplot illustrating the proportion of NFOL (Bcas1+/MBP-) and MFOL (Bcas1+/MBP+) found in each genotype (**x**^2^ (1) = 0.3524, p = ns). For J-K: N=4 animals per genotype from 4 litters, n=2-3 technical replicates/mouse.

**Figure S5:**
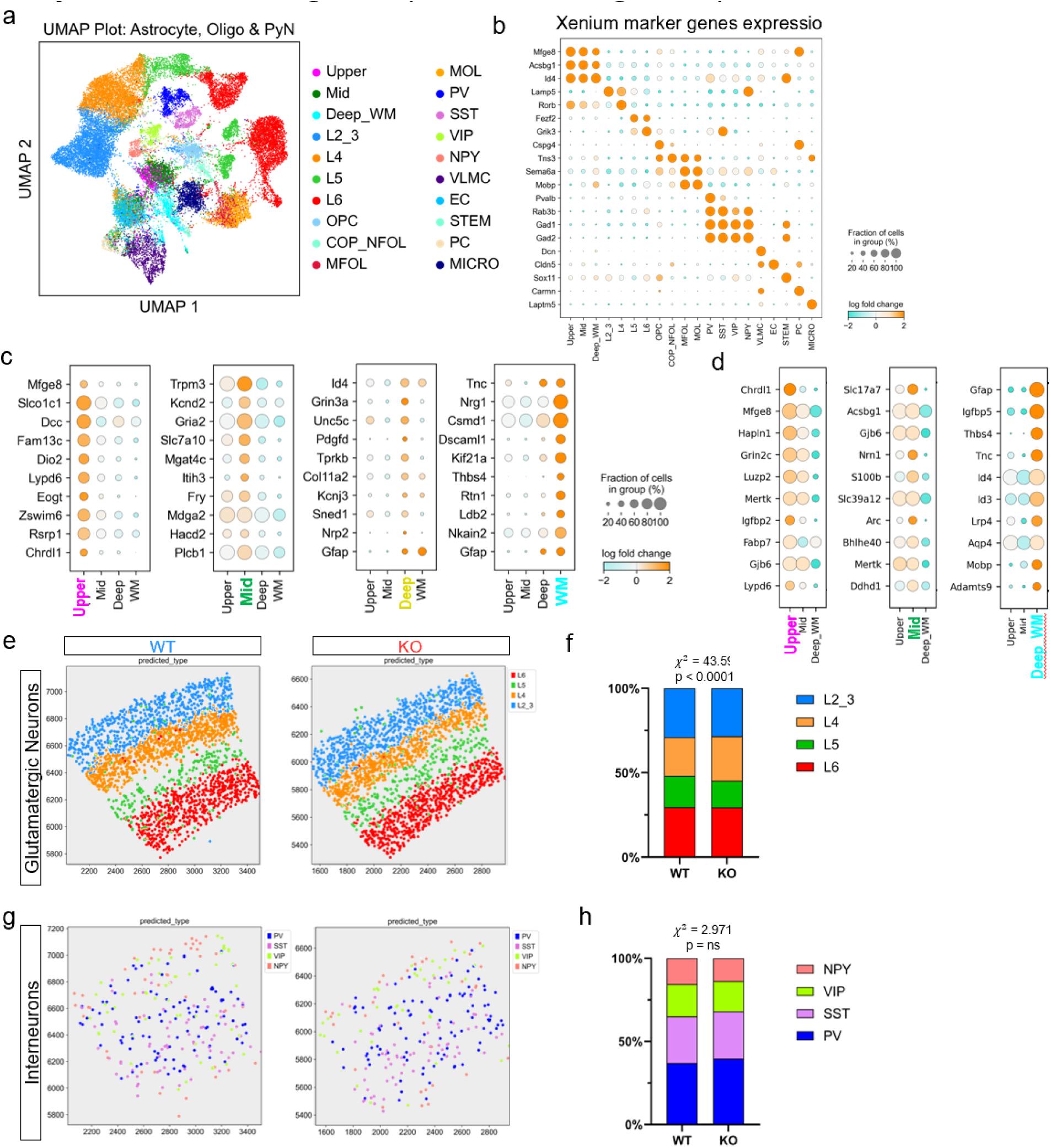
Spatial transcriptomic analysis of WT and Ctnnd2-KO visual cortex (related to Figure 4). A) UMAP V1 Xenium dataset, colored by cell identity. 31469 total cells detected (15222 WT, 16247 KO). B) Dotplot of marker gene expression across V1 cell types. C-D) Dotplots show cluster-defining genes for astrocyte subtypes from WT snRNAseq (C) and Xenium (D) data (Wilcoxon rank-sum test with Benjamini Hochberg correction comparing each cluster to every other cluster). Color of dot is log2 fold-change (log2FC), size of dot is fraction of cells expressing transcript. E-F) Spatial distribution (E) and proportion (F) of glutamatergic neuron subtypes in each genotype. Chi-square test of independence, (3) = 56.14 p < 0.0001. G-H) Spatial distribution (E) and proportion (F) of GABAergic interneuron subtypes in each genotype. Chi-square test of independence, (3) = 5.679 p = ns.

**Figure S6:**
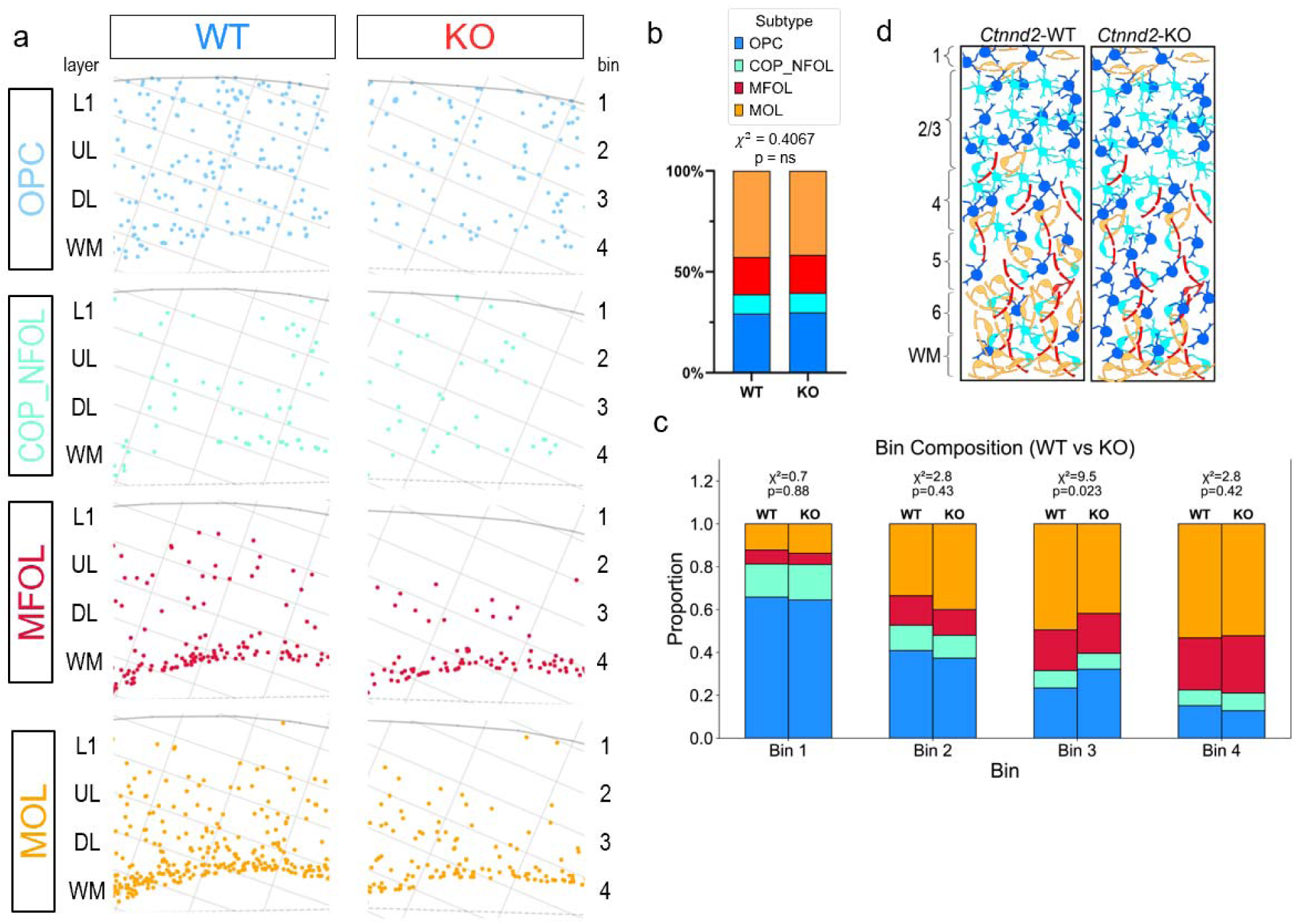
Spatial transcriptomic analysis of oligodendrocytes (related to Figure 4). A-B) Spatial distribution (A) and proportion (B) of oligodendrocyte lineage cell subtypes in each genotype. Chi-square test of independence, (3) = 0.4067, p = ns. C) Barplots show proportional contributions of each stage-specific subtype to the total oligodendroglia population within each of 4 equally sized cortical bins. D) Summary schematic of layer-specific changes in oligodendroglia subtypes.

**Figure S7:**
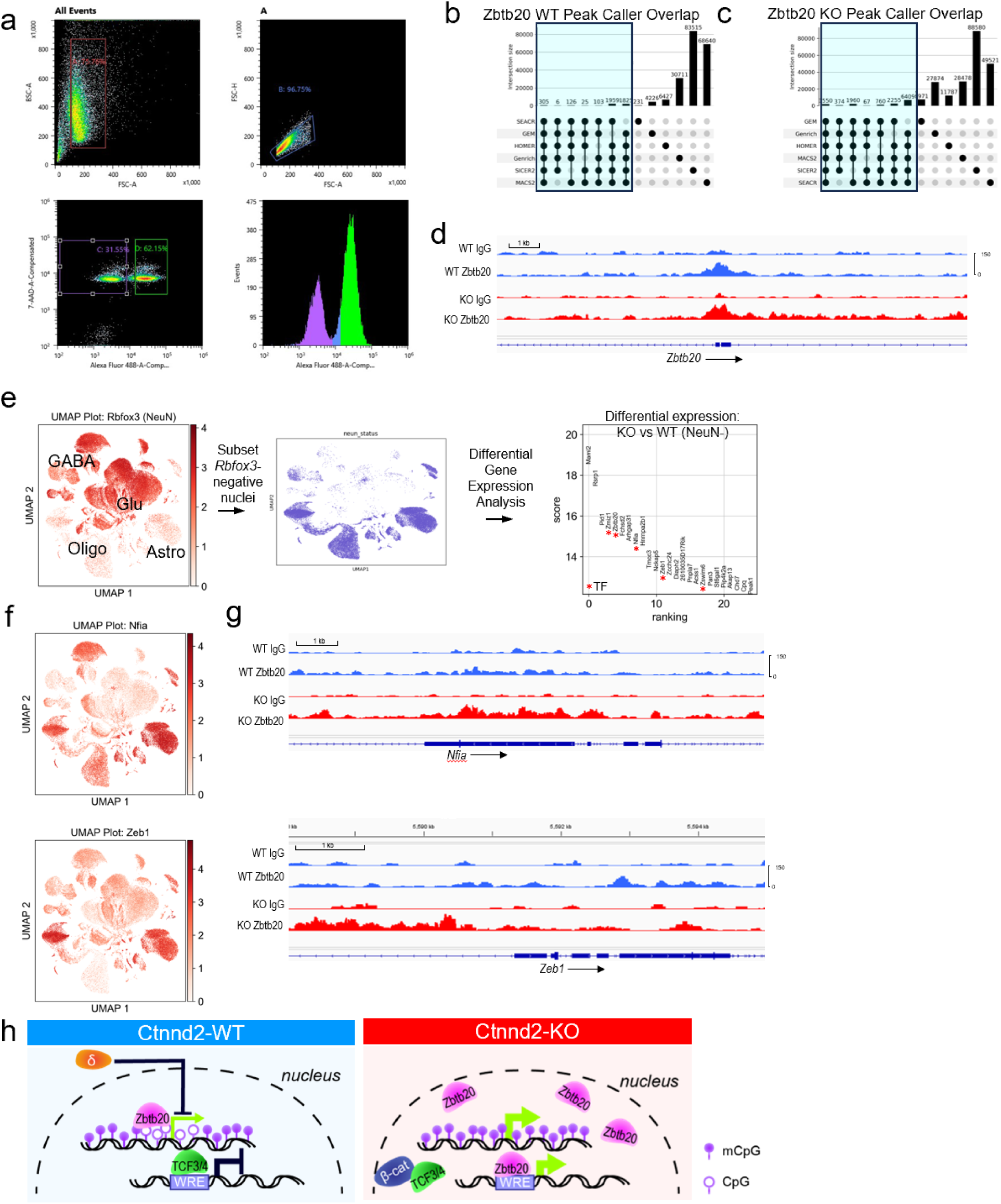
Quality control and additional multiomic CUT&RUN analyses (related to Figure 5). A) Fluorescence-activated nuclei sorting (FANS) gating strategy to isolate AF488-negative singlet glial nuclei from P28 cortex labeled by staining nuclei with AF488-conjugated NeuN antibody. Far left, debris exclusion gate; near left, singlet gate; near right, 7AAD+/AF488- nuclei gate; far right, AF488 histogram. B-C) UpSet plots showing overlap between the called peaks for WT (B) and KO (C) samples across all peak callers in the CARAS pipeline for Zbtb20 CUT&RUN. High confidence peak sets highlighted. D) Zbtb20 and IgG CUT&RUN read coverage at *Zbtb20* promoter. Merged tracks from N=4 WT and N=4 KO biological replicates. E) Bioinformatic pipeline for differential gene expression analysis of NeuN-negative nuclei. Asterisk indicates transcription factor or coactivator. F) UMAPs of all nuclei, annotated by expression of transcription regulator of interest. G) Zbtb20 and IgG CUT&RUN read coverage at *Nfia* (top) and *Zeb1* (bottom) promoters. Merged tracks from N=4 WT and N=4 KO biological replicates. H) Working model for cooperative transcriptional regulation by δ-catenin and Zbtb20.

**Figure S8:**
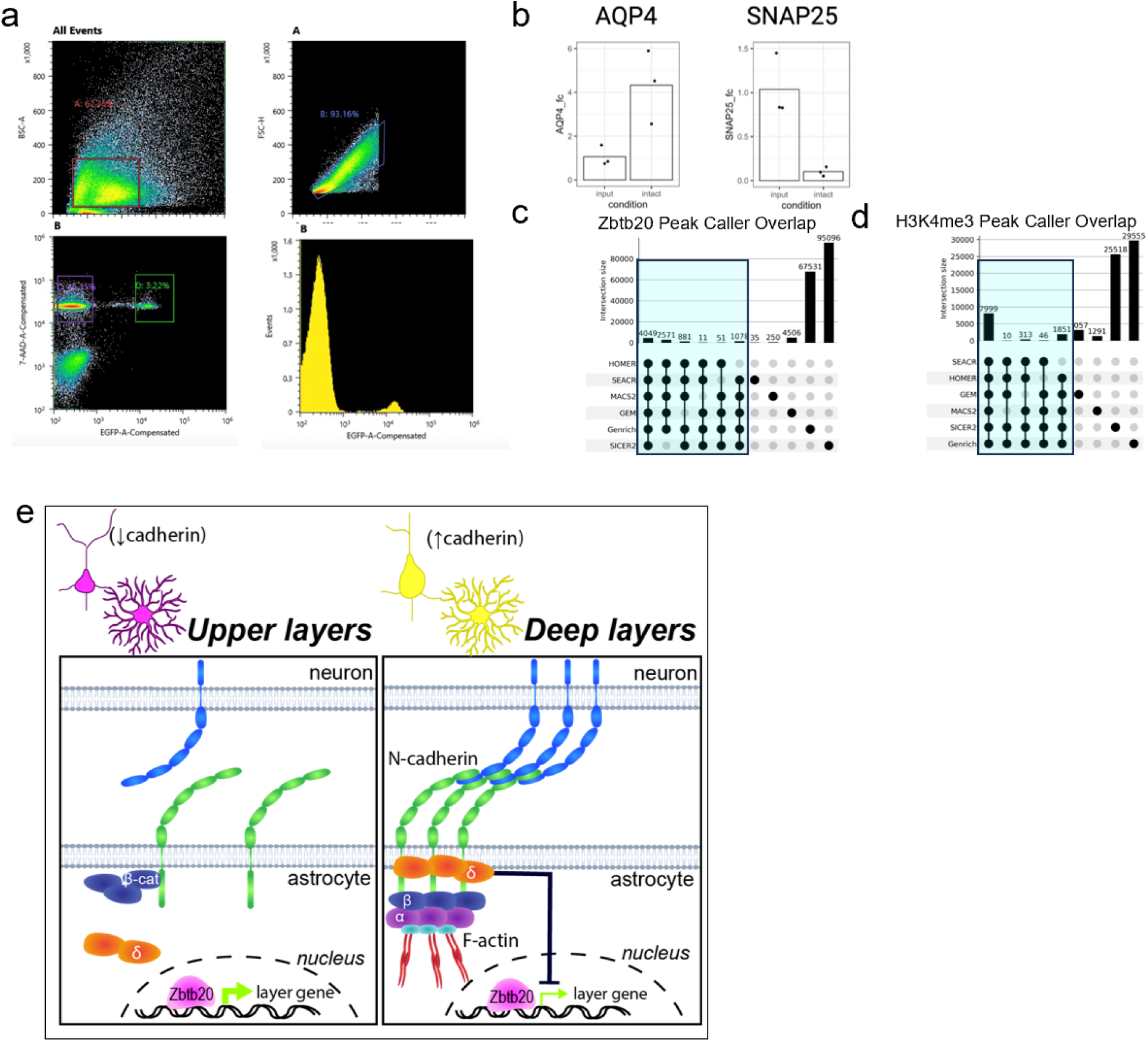
Quality control and additional astrocyte-specific CUT&RUN analyses (related to Figure 6). A) Fluorescence-activated nuclei sorting (FANS) gating strategy to isolate GFP+ singlet astrocyte nuclei from P21 cortex labeled by injecting CAG-Sun1/sfGFP mice with astrocyte-specific Cre AAV. Far left, debris exclusion gate; near left, singlet gate; near right, 7AAD+/GFP+ nuclei gate; far right, GFP histogram. B) Barplots showing mRNA expression of *Aqp4* and *Snap25* in GFP+ astrocyte nuclei isolated from p21 cortex by FANS. Values are fold-change compared to unsorted input nuclei. N=3 independent biological replicates.C-D) UpSet plots showing overlap between the called peaks for Zbtb20 (C) and H3K4me3 (D) CUT&RUN across all peak callers in the CARAS pipeline. E) Summary and working model for δ-catenin-dependent layer specificity of Zbtb20 transcriptional regulation in astrocytes.

### Supplementary Tables

**Table S1.** Statistical analyses for main and supplemental figures, related to Figures 1-6 and S1-S8.

| Panel | Quantity | Test | P-value and N |
| --- | --- | --- | --- |
| 1G | Density of DAPI+ nuclei by cortical bin | 2-way ANOVA | N: 8 littermate pairs (4 WT-M, 4 WT-F, 3 KO-M, 5 KO-F)<br>P(bin): <2e-16; P(genotype): 0.5144;<br>P(bin:genotype): 0.6976 |
| 1G | Density of NeuN+ nuclei by cortical bin | 2-way ANOVA | N: 8 littermate pairs (4 WT-M, 4 WT-F, 3 KO-M, 5 KO-F)<br>P(bin): 0.007065; P(genotype): 0.221239;<br>P(bin:genotype): 0.937642 |
| 1G | Density of Sox9+ nuclei by cortical bin | 2-way ANOVA | N: 8 littermate pairs (4 WT-M, 4 WT-F, 3 KO-M, 5 KO-F)<br>P(bin): 8.272e-13; P(genotype): 0.7825;<br>P(bin:genotype): 0.6566 |
| 1G | Density of Olig2+ nuclei by cortical bin | 2-way ANOVA | N: 8 littermate pairs (4 WT-M, 4 WT-F, 3 KO-M, 5 KO-F)<br>P(bin): 5.332e-13; P(genotype): 0.7913;<br>P(bin:genotype): 0.2505 |
| 1I | VC RNA sequencing log2 fold-change of gene expression in KO compared to WT visual cortex | Wald test with Benjamini–Hochberg’s correction | Criteria for DEG selection: log2 fold change (FC) less than -0.15 or greater than 0.15; false discovery rate (FDR) < 0.05 |
| 1J | Gene Ontology biological process category enrichment among upregulated genes | over-representation analysis using a hypergeometric distribution | Criteria for over-represented category: FDR < 0.05, q-value < 0.2 |
| 1K | Read count of upregulated genes in stemness-related GO categories | read count centered and scaled in the row direction |  |
| 1L | Gene Ontology biological process category enrichment among downregulated genes | over-representation analysis using a hypergeometric distribution | Criteria for over-represented category: FDR < 0.05, q-value < 0.2 |
| 1M | Read count of downregulated genes in oxphos-related GO categories | read count centered and scaled in the row direction |  |
| S1A | Genotype distribution of offspring in 10 independent litters from *Ctnnd2*-Het breeding pairs | Chi-square test of independence | N: 75 animals total (22 WT, 37 HET, 16 KO) |
| S1B | Sex distribution of offspring in 10 independent litters from *Ctnnd2*-Het breeding pairs | Chi-square test of independence | N: 38 animals total (10 WT-M, 12 WT-F, 9 KO-M, 7 KO-F) |
| S1F | Mouse body weight at P7, P8, P9, and P30 | Multiple unpaired two-tailed t-tests with Welch’s correction | N: 4-8 animals per genotype per age (8 WT and 6 KO at P7, P8, and P9; 4 WT and 4 KO at P30)<br>P(P7):6.03e-4; P(P8): p=1.8e-6; P(P9): p=6e-6; |
| | | | P(P30): $p=3.5e-2$ |
| S1G | Mouse brain weight at P30 | Welch's t-test | N: 4 sex-matched littermate pairs<br>P: <0.0001 |
| S1H | Mouse brain weight normalized to body weight at P30 | Welch's t-test | N: 4 sex-matched littermate pairs<br>P: 0.5873 |
| S1I | Overall density of DAPI | Welch's t-test | N: 8 littermate pairs (4 WT-M, 4 WT-F, 3 KO-M, 5 KO-F)<br>P: 0.6460 |
| S1I | Overall density of NeuN | Welch's t-test | N: 8 littermate pairs (4 WT-M, 4 WT-F, 3 KO-M, 5 KO-F)<br>P: 0.3981 |
| S1I | Overall density of Olig2 | Welch's t-test | N: 8 littermate pairs (4 WT-M, 4 WT-F, 3 KO-M, 5 KO-F)<br>P: 0.8350 |
| S1I | Overall density of Sox9 | Welch's t-test | N: 8 littermate pairs (4 WT-M, 4 WT-F, 3 KO-M, 5 KO-F)<br>P: 0.8861 |
| S2C | PFC RNA sequencing log2 fold-change of gene expression in KO compared to WT visual cortex | Wald test with Benjamini-Hochberg's correction | Criteria for DEG selection: log2 fold change (FC) less than -0.15 or greater than 0.15; false discovery rate (FDR) < 0.05 |
| 2E | Width of binocular zone by condition and genotype | 2-way ANOVA with Tukey's multiple comparisons test | N: 7 NR Ctrl (6 WT, 1 Het), 6 MD Ctrl (2 WT, 4 Het), 5 NR KO, 6 MD KO<br>ANOVA: P(genotype): 0.6773; P(condition): <0.0001; P(interaction): 0.6530<br>Tukey: P(KO:NR vs. KO:MD): 0.0043; P(CTRL:NR vs. CTRL:MD): 0.0340 |
| 2G | Width of binocular zone by condition and genotype | 2-way ANOVA with Tukey's multiple comparisons test | N: 5 NR Ctrl (1 WT, 5 Het), 8 MD Ctrl (2 WT, 6 Het), 6 NR KO, 6 MD KO<br>ANOVA: P(genotype): 0.3523; P(condition): 0.0160; P(interaction): 0.1844<br>Tukey: P(KO:NR vs. KO:MD): 0.8107; P(CTRL:NR vs. CTRL:MD): 0.0481 |
| 3B | Unbiased Leiden clustering of nuclei in visual cortex from 3 WT and 3 KO animals | Leiden algorithm | N: 99966 nuclei total |
| 3C | snRNA sequencing log2 fold-change of gene expression in KO compared to WT glutamatergic nuclei | Wilcoxon rank-sum | N: 53340 nuclei (31958 WT, 21382 KO) |
| 3C | snRNA sequencing log2 fold-change of gene expression in KO compared to WT GABAergic nuclei | Wilcoxon rank-sum | N: 7576 nuclei (4673 WT, 2903 KO) |
| 3C | snRNA sequencing log2 fold-change of gene expression in KO compared to WT oligodendroglia nuclei | Wilcoxon rank-sum | N: 16958 nuclei 8477 WT, 8481 KO) |
| 3C | snRNA sequencing log2 fold-change of gene expression in KO compared to WT astrocyte nuclei | Wilcoxon rank-sum | N: 12157 nuclei 6110 WT, 6047 KO) |
| 3D | Gene Ontology biological process category enrichment among downregulated genes | over-representation analysis using a hypergeometric distribution | Criteria for over-represented category: FDR < 0.05, q-value < 0.2 |
| 3E | Subtype distribution of glutamatergic nuclei in visual cortex from 3 WT and 3 KO animals | Chi-square test of independence | N: 53340 nuclei (31958 WT, 21382 KO) |
| 3F | Subtype distribution of GABAergic nuclei in visual cortex from 3 WT and 3 KO animals | Chi-square test of independence | N: 7576 nuclei (4673 WT, 2903 KO) |
| 3G | Subtype distribution of oligodendroglia nuclei in visual cortex from 3 WT and 3 KO animals | Chi-square test of independence | N: 16958 nuclei (8477 WT, 8481 KO) |
| 3H | Subtype distribution of astrocyte nuclei in visual cortex from 3 WT and 3 KO animals | Chi-square test of independence | N: 12157 nuclei (6110 WT, 6047 KO) |
| S4B | Spatial frequency distribution of Cux1+/RORB-/Ctip2- nuclei | 2-way ANOVA | N: 4 littermate pairs (1 WT-M, 3 WT-F, 0 KO-M, 4 KO-F)<br>P(bin): <2e-16; P(genotype): 1.0000;<br>P(bin:genotype): 0.3805 |
| S4B | Spatial frequency distribution of Cux1-/RORB+/Ctip2- and Cux1+/RORB+/Ctip2- nuclei | 2-way ANOVA | N: 4 littermate pairs (1 WT-M, 3 WT-F, 0 KO-M, 4 KO-F)<br>P(bin): <2e-16; P(genotype): 1.0000;<br>P(bin:genotype): 0.9615 |
| S4B | Spatial frequency distribution of Cux1-/RORB-/Ctip2+ and Cux1+/RORB-/Ctip2+ nuclei | 2-way ANOVA | N: 4 littermate pairs (1 WT-M, 3 WT-F, 0 KO-M, 4 KO-F)<br>P(bin): <2e-16; P(genotype): 1.0000;<br>P(bin:genotype): 0.9557 |
| S4C | Subtype distribution of glutamatergic nuclei in visual cortex from 4 WT and 4 KO animals | 2-way ANOVA | N: 12095 nuclei (5745 WT, 6350 KO)<br>Chi-square: 22.28, df: 2, P: <0.0001 |
| S4D | Density of Cux1+/RORB-/Ctip2- nuclei within L2/3 ROI | Welch's t-test | N: 4 littermate pairs (1 WT-M, 3 WT-F, 0 KO-M, 4 KO-F)<br>P: 0.4444 |
| S4D | Density of Cux1-/RORB+/Ctip2- and Cux1+/RORB+/Ctip2- within L4 ROI | Welch's t-test | N: 4 littermate pairs (1 WT-M, 3 WT-F, 0 KO-M, 4 KO-F)<br>P: 0.1270 |
| S4D | Density of Cux1-<br>/RORB-/Ctip2+ and<br>Cux1+/RORB-/Ctip2+<br>nuclei within L5/6 ROI | Welch's t-test | N: 4 littermate pairs (1 WT-M, 3 WT-F, 0 KO-M, 4 KO-F)<br>P: 0.0933 |
| S4F | Spatial frequency<br>distribution of Olig2+<br>nuclei | 2-way ANOVA | N: 3 littermate pairs (1 WT-M, 2 WT-F, 2 KO-M, 1 KO-F)<br>P(bin): 6.526e-13; P(genotype): 1.0000;<br>P(bin:genotype): 0.8088 |
| S4F | Spatial frequency<br>distribution of<br>Olig2+/Pdgfra+/Aspa-<br>nuclei | 2-way ANOVA | N: 3 littermate pairs (1 WT-M, 2 WT-F, 2 KO-M, 1 KO-F)<br>P(bin): 0.003315; P(genotype): 1.000000;<br>P(bin:genotype): 0.791523 |
| S4F | Spatial frequency<br>distribution of<br>Olig2+/Pdgfra-/Aspa+<br>nuclei | 2-way ANOVA | N: 3 littermate pairs (1 WT-M, 2 WT-F, 2 KO-M, 1 KO-F)<br>P(bin): 1.472e-13; P(genotype): 1.000;<br>P(bin:genotype): 0.479 |
| S4G | Subtype distribution of<br>oligodendrocyte nuclei<br>in visual cortex from 3<br>WT and 3 KO animals | 2-way ANOVA | N: 4046 cells (2028 WT, 2018 KO)<br>Chi-square: 63.84, df: 2, P: <0.0001 |
| S4H | Overall density of<br>Olig2+ nuclei | Welch's t-test | N: 3 littermate pairs (1 WT-M, 2 WT-F, 2 KO-M, 1 KO-F)<br>P: 0.8695 |
| S4H | Overall density of<br>Olig2+/Pdgfra+/Aspa-<br>nuclei | Welch's t-test | N: 3 littermate pairs (1 WT-M, 2 WT-F, 2 KO-M, 1 KO-F)<br>P: 0.1794 |
| S4H | Overall density of<br>Olig2+/Pdgfra-/Aspa+<br>nuclei | Welch's t-test | N: 3 littermate pairs (1 WT-M, 2 WT-F, 2 KO-M, 1 KO-F)<br>P: 0.3264 |
| S4J | Spatial frequency<br>distribution of<br>Bcas1+/MBP- cells | 2-way ANOVA | N:6 littermate pairs (3 WT-M, 3 WT-F, 5 KO-M, 1 KO-F)<br>P(bin): 1.496e-12; P(genotype): 0.7947;<br>P(bin:genotype): 0.7745 |
| S4J | Spatial frequency<br>distribution of<br>Bcas1+/MBP+ cells | 2-way ANOVA | N:6 littermate pairs (3 WT-M, 3 WT-F, 5 KO-M, 1 KO-F)<br>P(bin): <2e-16; P(genotype): 0.8083;<br>P(bin:genotype): 0.7717 |
| S4K | Subtype distribution of<br>Bcas1+ cells in visual<br>cortex from 6 WT and 6<br>KO animals | Chi-square test of<br>independence | N: 515 cells (251 WT, 264 KO)<br>Chi-square: 0.3524, df: 1, P: 0.5528 |
| 4D | Gene expression in<br>*Ctnnd2*-KO compared to<br>WT samples in V1 ROI | Wilcoxon rank-sum test<br>with Benjamini<br>Hochberg correction<br>comparing KO to WT | 20 highest Wilcoxon scores |
| 4E | Gene expression in<br>*Ctnnd2*-KO compared to<br>WT samples in V1 ROI | Wilcoxon rank-sum test<br>with Benjamini<br>Hochberg correction<br>comparing KO to WT | 20 lowest Wilcoxon scores |
| 4G | Subtype distribution of<br>astrocytes in V1 ROI<br>from 2 WT and 2 KO<br>animals | Chi-square test of<br>independence | N: 3117 nuclei total (1576 WT, 1541 KO)<br>Chi-square: 13.24, df: 2, P: 0.0013 |
| 4H | Subtype distribution of astrocytes in V1 bin 1 ROI from 2 WT and 2 KO animals | Chi-square test of independence | N: 1073 nuclei total (527 WT, 546 KO)<br>Chi-square: 11.8987, df: 2, P: 0.0026 |
| 4H | Subtype distribution of astrocytes in V1 bin 2 ROI from 2 WT and 2 KO animals | Chi-square test of independence | N: 688 nuclei total (343 WT, 345 KO)<br>Chi-square: 3.3452, df: 2, P: 0.1878 |
| 4H | Subtype distribution of astrocytes in V1 bin 3 ROI from 2 WT and 2 KO animals | Chi-square test of independence | N: 625 nuclei total (316 WT, 309 KO)<br>Chi-square: 11.9400, df: 2, P: 0.0026 |
| 4H | Subtype distribution of astrocytes in V1 bin 4 ROI from 2 WT and 2 KO animals | Chi-square test of independence | N: 727 nuclei total (388 WT, 339 KO)<br>Chi-square: 1.9180, df: 2, P: 0.38 |
| S5F | Subtype distribution of glutamatergic neurons in V1 ROI from 2 WT and 2 KO animals | Chi-square test of independence | N: 18264 nuclei total (8380 WT, 9884 KO)<br>Chi-square: 43.59, df: 3, P: <0.0001 |
| S5H | Subtype distribution of GABAergic neurons in V1 ROI from 2 WT and 2 KO animals | Chi-square test of independence | N: 3538 nuclei total (1850 WT, 1688 KO)<br>Chi-square: 2.971, df: 3, P: 0.3961 |
| S6B | Subtype distribution of oligodendroglia in V1 ROI from 2 WT and 2 KO animals | Chi-square test of independence | N: 3117 nuclei total (1576 WT, 1541 KO)<br>Chi-square: 0.4067, df: 3, P: 0.9388 |
| S6C | Subtype distribution of oligodendroglia in V1 bin 1 ROI from 2 WT and 2 KO animals | Chi-square test of independence | N: 526 nuclei total (278 WT, 248 KO)<br>Chi-square: 0.6876, df: 3, P: 0.8761 |
| S6C | Subtype distribution of oligodendroglia in V1 bin 2 ROI from 2 WT and 2 KO animals | Chi-square test of independence | N: 613 nuclei total (313 WT, 300 KO)<br>Chi-square: 2.7748, df: 3, P: 0.4277 |
| S6C | Subtype distribution of oligodendroglia in V1 bin 3 ROI from 2 WT and 2 KO animals | Chi-square test of independence | N: 914 nuclei total (479 WT, 435 KO)<br>Chi-square: 9.5061, df: 3, P: 0.02327 |
| S6C | Subtype distribution of oligodendroglia in V1 bin 4 ROI from 2 WT and 2 KO animals | Chi-square test of independence | N: 1481 nuclei total (778 WT, 703 KO)<br>Chi-square: 2.8181, df: 3, P: 0.4205 |
| 5E | Genomic feature distribution of high confidence peaks in 4 WT and 4 KO samples | Chi-square test of independence | N: 17724 loci (4349 WT 13375 KO)<br>Chi-square: 283.6, df: 6, P: <0.0001 |
| 5J | Motif enrichment among WT-only Zbtb20-bound promoters | over-representation analysis using a hypergeometric distribution | N: 2103 WT promoters, 17807 background promoters<br>Criteria for enriched motif: p-value < 0.05 |
| 5J | Motif enrichment among KO-only Zbtb20-bound promoters | over-representation analysis using a hypergeometric distribution | N: 951 WT promoters, 18868 background promoters<br>Criteria for enriched motif: p-value < 0.05 |
| 5K | DEG enrichment among genes with Zbtb20-bound promoters | Fisher's exact test | N: 35795 genes total (32227 overall, 3568 Zbtb20-bound) |
| 6E | Gene expression in *Ctnnd2*-KO compared to WT astrocyte nuclei | Wilcoxon rank-sum test with Benjamini Hochberg correction comparing KO to WT |  |
| 6F | DEG enrichment among genes with layer-enriched expression in astrocytes | Fisher's exact test | N: 3343 genes total (471 DEG, 2872 non-DEG)<br>P: <0.0001 |
| 6I | Normalized mean fluorescent intensity of *Tnc* by cortical bin | 2-way ANOVA | N: 3 littermate pairs (2 WT-M, 1 WT-F, 2 KO-M, 1 KO-F)<br>P(bin): 9.157e-15; P(genotype): 0.1090;<br>P(bin:genotype): 0.7546 |
| 6J | Normalized mean fluorescent intensity of *Id4* by cortical bin | 2-way ANOVA | N: 3 littermate pairs (2 WT-M, 1 WT-F, 2 KO-M, 1 KO-F)<br>P(bin): 9.431e-16; P(genotype): 0.6188;<br>P(bin:genotype): 0.7279 |
| 6K | Normalized mean fluorescent intensity of *Mfge8* by cortical bin | 2-way ANOVA with Tukey's multiple comparisons test | N: 3 littermate pairs (2 WT-M, 1 WT-F, 2 KO-M, 1 KO-F)<br>ANOVA: P(bin): 2.462e-12; P(genotype): 0.022735;<br>P(bin:genotype): 0.004651<br>Tukey: P(bin1): 0.0321; P(bin7): 0.0355; P(bin9): 0.0138; P(bin10): 0.0004 |

**Table S2.** Differentially expressed genes from bulk RNAseq, related to Figure 1

**Table S3.** Differentially expressed genes from snRNAseq, related to Figures 3, 4, 5, and 6

**Table S4.** Astrocyte subcluster-defining genes from snRNAseq, related to Figures 3 and 4

**Table S5.** Custom Xenium panel information

**Table S6.** Astrocyte subcluster-defining genes from Xenium, related to Figure 4

**Table S7.** Differentially expressed genes from Xenium, related to Figure 4

**Table S8.** Homer enriched motifs, related to Figure 5

